# Proteomic Analysis of *Plasmodium* Merosomes: The Link Between Liver and Blood Stages in Malaria

**DOI:** 10.1101/580266

**Authors:** Melanie J Shears, Raja Sekhar Nirujogi, Kristian E Swearingen, Santosh Renuse, Satish Mishra, Panga Jaipal Reddy, Robert L Moritz, Akhilesh Pandey, Photini Sinnis

## Abstract

The pre-erythrocytic liver stage of the malaria parasite, comprising sporozoites and the liver stages into which they develop, remains one of the least understood parts of the lifecycle, in part owing to the low numbers of parasites. Nonetheless, it is recognized as an important target for anti-malarial drugs and vaccines. Here we provide the first proteomic analysis of merosomes, which define the final phase of the liver stage and are responsible for initiating the blood stage of infection. We identify a total of 1879 parasite proteins, and a core set of 1188 proteins quantitatively detected in every biological replicate, providing an extensive picture of the protein repertoire of this stage. This unique dataset will allow us to explore key questions about the biology of merosomes and hepatic merozoites.

**Highlights:**

- First proteome of the merosome stage of malaria parasites
- Quantitative detection of 1188 parasite proteins across 3 biological replicates
- Comparison to blood stage proteomes identifies shared and unique proteins
- Discovery of cleaved PEXEL motifs highlights liver stage protein export

**In Brief:** The merosome stage that links malaria liver and blood stage infection is poorly understood. Here we provide the first proteome of this life cycle stage using the *Plasmodium berghei* rodent malaria model.

**Graphical Abstract:** 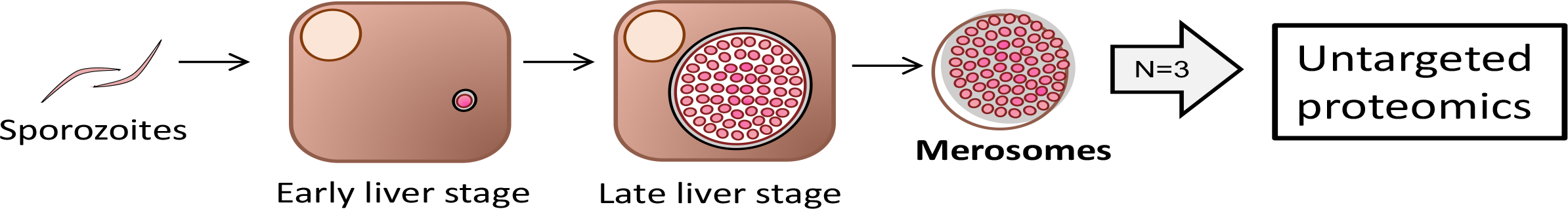

## Introduction

Malaria remains one of the most important infectious diseases in the world, impacting approximately a third of the world’s population and causing an estimated 400,000 deaths annually (1). The initial stage of malaria infection, comprised of infectious sporozoites inoculated by mosquitoes and the liver stages into which they develop, is clinically asymptomatic yet has been validated as a vaccine target in both rodent malaria models (2, 3) and human Phase III clinical trials (4, 5). The low numbers of parasites during this pre-erythrocytic stage combined with their decreased allelic diversity compared to blood stage parasites, explain, at least in part, why targeting this stage is advantageous [reviewed in (6)]. However, the limitations of current pre-erythrocytic stage vaccines indicate that additional strategies or targets are still required. Building upon this success will necessitate identifying new pre-erythrocytic stage targets and expanding the repertoire of antigens to include late liver stage proteins (7). In this study we focus on the final phase of the *Plasmodium* pre-erythrocytic stage, the merosomes, characterizing their proteome to obtain a better understanding of their biology and provide data to inform target selection for future interventions.

Merosomes and the hepatic merozoites they contain are the link between the asymptomatic pre-erythrocytic stage and the clinically important erythrocytic stage of malaria infection. They are the product of an impressive replication: A single sporozoite that has successfully entered a hepatocyte can transform and give rise to 5000 to 10,000 hepatic merozoites in a matter of days (8). Development of liver stage parasites occurs within a parasitophorous vacuole (PV), a parasite-derived structure that separates the newly-invaded parasite from the host cell cytoplasm (9). Within this vacuole, the parasite acquires nutrients from its host necessary for growth and replication (8, 10, 11). Once mature, the PV membrane ruptures, releasing the hepatic merozoites into the host cell cytoplasm (12). Merozoites bud from the host hepatocyte in packets called merosomes, each containing 10 to 1000 hepatic merozoites and surrounded by modified host cell membrane (13). These merosomes enter the bloodstream and release hepatic merozoites to initiate blood stage of infection. Iterative cycles of replication then give rise to erythrocytic merozoites, which are released and invade naïve erythrocytes, leading to high parasite numbers and clinical disease.

Hepatic and erythrocytic merozoites are morphologically similar, sharing specialized apical organelles, motility apparatus, and surface proteins necessary for erythrocyte invasion. Though frequently assumed to be identical, their different tissue origins, together with the unique mechanism of hepatic merozoite release, suggests these two merozoite populations may differ in biologically important ways. To date, only two studies have examined the differences between these merozoite populations: Hepatic and erythrocytic merozoites were shown to express different members of the Py235 protein family (14), and more recently, the discovery that deletion of the cysteine protease bergheipain-1 differentially impacts these distinct merozoite populations (15). Whether other differences exist between hepatic and erythrocytic merozoites is currently unknown, and questions also remain concerning the host-parasite interactions required for merosome formation and release.

Our understanding of the biology of *Plasmodium* liver stages, including the late-stage merosomes and hepatic merozoites, has been hampered by technical difficulties stemming from their inaccessibility and low numbers of parasites. Rodent malaria models such as *P. berghei* and *P. yoelii* offer easier access to liver stage parasites, and as such have been invaluable for defining the biology of this malaria life cycle stage (16). Using the *P. yoelii* model, a transcriptomic and proteomic study of the liver stage has been reported although it did not include merosomes or hepatic merozoites (17). Thus, we still lack a comprehensive picture of the protein repertoire of the final phase of the pre-erythrocytic stage of infection. Here we use the rodent malaria parasite *P. berghei* to survey the proteome of merosomes and the hepatic merozoites within. By scaling up and modifying existing liver stage culture methods, we reproducibly generate large numbers of intact merosomes for analysis. Using high-resolution Orbitrap mass spectrometry, we perform in-depth quantitative proteomic profiling of three biological replicate samples. This dataset provides an important foundation for furthering our understanding of the basic biology of merosomes and hepatic merozoites.

## Experimental Procedures

### Production of *P. berghei* Sporozoites

Sporozoites were produced by allowing *Anopheles stephensi* mosquitoes to blood feed on mice infected with *P. berghei* ANKA using standard procedures. All animal work was conducted in accordance with the National Institutes of Health Guide for the Care and Use of Laboratory Animals under an approved Institutional Animal Care and Use Committee protocol. Sporozoites were isolated at 20-21 days post infection by dissection of salivary glands into cold DMEM (Corning #10-013) containing 1000 IU Penicillin and 1000 μg/mL Streptomycin (Corning #30-002) and 25 μg/mL Amphotericin B (Corning #30-003). Salivary glands were homogenized and mosquito debris pelleted by centrifugation at 100 x *g* for 2 min at 4°C. Sporozoites were then transferred to a new tube, counted by hemocytometer, and incubated briefly on ice until use.

### HepG2 Cell Culture and Infection

HepG2 human hepatoma cells were maintained at 37°C and 5% CO_2_ in complete medium, consisting of DMEM, 100 IU Penicillin and 100 μg/mL Streptomycin, 2mM L-glutamine (Corning #25-005) and 10% heat inactivated fetal bovine serum (FBS; Biowest #S1520). Cells were plated in three 24-well plates pre-coated with collagen (Corning #354236) the day before infection at a density of 100,000 cells per well. Sporozoites were diluted in complete medium containing 1 μg/mL mycamine (Astellas Pharma), and added to cells to give 200,000 to 400,000 sporozoites per well. Plates were centrifuged at 310 x *g* for 3 min at room temperature (RT) and incubated at 37°C. After 2 h, the medium containing unattached sporozoites was removed and wells were washed twice with media as above before continued incubation. Infected cells were cultured for a total of 65 h with twice daily media changes, using media with 0.12 - 1 μg/mL mycamine for the first 48 h, and media without mycamine thereafter.

### Merosome Isolation

Culture supernatants from infected HepG2 cells were collected at 65 h post infection and pooled into a 15 mL tube. Merosomes and detached cells were pelleted by centrifugation at 2,000 x *g* for 5 min at 4°C and the supernatant was removed. The pellet was washed twice in 10 mL of Hank’s Buffered Saline Solution (HBSS; Gibco #14025-029) with centrifugation as above. The pellet was then resuspended in 100 μL HBSS, and 6 μL was taken for counting by hemocytometer and quantitative real time PCR (qPCR). All samples were snap frozen in liquid nitrogen and stored at −80°C until analysis.

### Experimental Design and Statistical Rationale

Three independent biological replicates were prepared as above for proteomic analysis. This number was selected as we have previously demonstrated that between one and six replicates provide sufficient proteomic coverage for untargeted analysis of *Plasmodium* parasites (18, 19).

### Merosome Immunofluorescence Staining for Quality Assessment

Merosomes were spun onto poly-L-lysine (Sigma #P8920) coated glass coverslips by centrifugation at 200 x *g* for 10 min at RT, then fixed in 4% paraformaldehyde in phosphate buffered saline (PBS) for 20 min at RT. Cells were permeabilized in ice-cold methanol for 1 h at −20°C, washed with PBS, and incubated in 1% bovine serum albumin (BSA) in PBS for 1 h at RT. Primary antibody incubations were performed with rabbit anti-MSP1-19 (MR4 #MRA-23, kindly provided by Scott Linder, Penn State USA) or rabbit anti-UIS4 (20) in 1% BSA/PBS for 1 h at RT. Secondary antibody incubations were performed with goat anti-rabbit AlexaFluor 488 (Molecular Probes #A11008) as above or overnight at 4°C. Nuclei were stained with Hoechst 33342 in PBS for 5 min, and coverslips were mounted using Prolong Gold Antifade mountant (Molecular Probes #P10144).

Imaging for merosome quality assessment was performed using a Zeiss AxioImager M2 fluorescence microscope fitted with a Hamamatsu ORCA-R2 C10600 digital camera and 63x oil plan-APOCHROMAT objective. Image acquisition was performed using Volocity software version 6.3.1 (Perkin Elmer). Images were deconvolved using by iterative restoration algorithm, and image registration, maximum projection and contrast adjustments made using Volocity or FIJI Image J software (21).

### Immunofluorescence Staining for Localization of PHIST Protein

For staining liver stage parasites, HepG2 cells were plated onto collagen-coated glass coverslips, infected with sporozoites and cultured for 60 h before fixation, permeabilization and blocking as above. For staining merosomes and free merozoites, merosomes were collected as described and spun onto poly-L-lysine coated glass coverslips, then fixed in 4% paraformaldehyde, permeabilized and blocked as above. Cells were then incubated with rabbit anti-UIS4 and goat anti-rabbit AlexaFluor 488, followed by overnight incubation with rabbit anti-PHIST protein [PBANKA_1145400; (22)] conjugated to TexasRed (Abcam Cat #ab195225). Nuclei were stained with Hoechst 33342 and coverslips were mounted as above.

Localization studies were performed using a Zeiss LSM 710 confocal microscope with a plan-apochromat 63x/1.40 oil objective and Zen 2012 software. Imaging parameters were as follows: a 680-pixel x 680-pixel x 30-slice Z-stack was taken at each position, with averaging of 2 and voxel dimensions of 100 nm x 100 nm x 460 nm. Image analysis was performed using Bitplane Imaris 8.3.1. Colocalization statistics for PHIST and UIS4 was calculated using the Imaris colocalization module from images of 75 liver stage parasites from two independent experiments.

### Merozoite Genome Counting by Quantitative Real Time PCR

The number of individual merozoite genomes in each replicate was quantified by qPCR using primers specific for a region of the HSP70 gene (primers 5’-TGCAGCAGATAATCAAACTC-3’ and 5’-ACTTCAATTTGTGGAACACC-3’). Snap-frozen merosome sample aliquots were diluted 1:100 or 1:1000 in deionized water and 4 μL was mixed with 21 μL of Power SYBR Green PCR Mastermix (Applied Biosystems # 4367659) containing 800 nM of each primer per reaction. PCR cycles were performed using a StepOnePlus instrument (Applied Biosystems) as follows: initial denaturation at 95°C for 10 min; 40 cycles of denaturation at 95°C for 10 s, primer annealing at 42°C for 20 s, and extension at 60°C for 40 s. Genome numbers in merosome samples were estimated by comparison to a four-point parasite genomic DNA dilution series made from blood stage parasites, which was calibrated against a standard containing a fixed number of sporozoite genomes. All samples were assayed in triplicate and the reproducibility of the standard dilution series was cross-checked between experiments.

### Sample Preparation and Mass Spectrometry

Samples were resuspended in lysis buffer containing 9 M Urea in 50 mM triethylammonium bicarbonate pH 8.5 and sonicated for 30 s for 3 cycles. Protein lysates were centrifuged at 17,000 × *g* for 10 min and the cleared lysates were reduced by adding 5 mM dithiothreitol with incubation at 56°C for 20 min and then alkylated by adding 20 mM iodoacetamide and incubating at RT in the dark for 20 min. Proteins were digested by adding sequencing grade trypsin (1:20 substrate:enzyme; Promega #5111) and incubating at 37°C for 16 h. The digestion was then quenched by adding trifluoroacetic acid to a final concentration of 1% (v/v).

Peptides were fractionated using stage tip-based strong cation exchange (SCX) fractionation as described (23). Briefly, the SCX stage tips were prepared using 20 mm syringe plunger from SCX disc (3M Empore #2251), with three layers packed in 200 µL pipette tip. The SCX stage tips were activated by adding 100% acetonitrile (Fisher Scientific #A998) with centrifugation at 1000 x *g* for 2 min. The acidified peptide digests were loaded onto the tips and then cleaned twice with 0.2% trifluoroacetic acid with centrifugation as above. Freshly prepared elution buffers were added sequentially and the eluent was collected in a new tube by centrifugation at 400 x *g* for 2 min. Fractions 1 to 5 were eluted by adding buffer containing X mM ammonium acetate, 20% (v/v) acetonitrile in 0.5% (v/v) formic acid, where “X” was 50 mM, 75 mM, 125 mM, 200 mM and 300 mM, respectively. The sixth fraction was eluted by adding 5% (v/v) ammonium hydroxide in 80% (v/v) acetonitrile. All eluted fractions were then vacuum dried and stored at −80°C until analysis.

Fractions from the three biological replicates were analyzed separately on an Orbitrap Fusion Lumos ETD mass spectrometer interfaced with Easy-nanoLC 1200 nanoflow liquid chromatography system (Thermo Scientific). Peptides from each fraction were reconstituted in 0.1% formic acid and loaded on a pre-column (PepMap C_18_, 100 μm × 2 cm, Thermo Scientific) at a flow rate of 5 μL per min. Peptides were resolved on the analytical column (PepMap C_18_, 75 μm×50 cm, 2μ, Thermo Scientific) at 300 nL/min flow rate using a step gradient of 5% to 22% solvent B (0.1% formic acid in 95% acetonitrile) over 130 min, then 22% to 32% solvent B for 25 min, followed by column wash and reconditioning to give a total run time of 180 min. The EasySpray ion source was operated at 2.3 kV. Mass spectrometry data was acquired in a data-dependent manner with a survey scan in the range of *m/z* 300 to 1500. Both MS and MS/MS were acquired and measured using Orbitrap mass analyzer. Full MS scans were measured at a resolution of 120,000 at *m/z* 200. Total scan cycle time was set to 3.5 s. MS/MS fragmentation was carried out using higher-energy collisional dissociation (HCD) method with normalized collision energy (NCE) of 32 and detected at a mass resolution of 30,000 at *m/z* 200. Automatic gain control for full MS was set to 2E5 ions and for MS/MS 5E4 ions with a maximum ion injection time of 40 ms and 150 ms, respectively. Dynamic exclusion was set to 30 s and singly charged ions and ions with unknown charge states were rejected. Internal calibration was carried out using lock mass option (*m/z* 445.120025) from ambient air.

Peak picking followed by mass spectrometry data searching was performed using MaxQuant’s (1.5.5.1 version) Andromeda search engine against a combined database of *P. berghei* ANKA (PlasmoDB v26), *H. sapiens* (NextProt February 2016 release), *B. taurus* (NCBI Refseq 104), *A. stephensi* (Vectorbase Version 2.1), and common contaminant proteins. This combined database was compiled in-house using MaxQuant and contained a total of 133,486 proteins. The search was set up by creating three replicate experiments with six SCX fractions for each, allowing for matching between runs within each fraction and between replicates. Fully tryptic peptides with a maximum of two missed cleavages were considered, and peptide lengths were set to a minimum of 7 and maximum of 25 amino acids. Carbamidomethylation of Cys was set as a fixed modification, and oxidation of Met, and phosphorylation of Ser, Thr, or Tyr set as variable modifications. First search peptide precursor tolerance was set to 20 ppm and main search tolerance to 4.5 ppm, with a maximum charge of 7 allowed. MS/MS fragment ion mass tolerance was set to 7 ppm. For peptide spectrum matches (PSMs), peptides, and proteins, a false discovery rate (FDR) of <0.01 was applied. This value was chosen as it is conservative for reliable identifications (24). The FDR was calculated using target-decoy search. To maximize the number of identifications, a second peptide search and Match Between Runs were performed with a 0.7 min and 20 min match time window and alignment time window, respectively. Label-free quantification (LFQ) and intensity-based absolute quantification (iBAQ) was carried out by enabling LFQ and iBAQ within the MaxQuant software suite as described (25) with a minimum of 1 ratio count for pair-wise comparisons. Mass spectrometry proteomics data have been deposited to the ProteomeXchange Consortium via the PRIDE (26) partner repository with the dataset identifier PXD010559.

### Data Analysis

Proteins originating from *P. berghei*, *H. sapiens*, *B. taurus*, *A. stephensi*, and common contaminant proteins were identified in all samples as expected from the choice of host cell, method of culture, and media components. For all data analysis, we focused on *P. berghei* proteins only. The percent *P. berghei* genome coverage, percent of functionally annotated proteins, and percent of proteins with syntenic *P. falciparum* orthologs were determined using the pre-defined search strategies in PlasmoDB (27). Comparison to the published *P. berghei* rhoptry proteome (28) was performed by converting proteins in the dataset to current PlasmoDB IDs and comparing to merosome proteins with Venny (http://bioinfogp.cnb.csic.es/tools/venny/). Comparisons to published *P. yoelii* (17) or *P. falciparum* proteomes (29–33) were performed by identifying syntenic *P. berghei* orthologs for each dataset and comparing to merosome proteins as above.

Gene Ontology (GO) term analysis was performed by downloading GO term annotations for merosome proteins and their syntenic *P. falciparum* orthologs from PlasmoDB. GO biological process annotations for *P. falciparum* orthologs of merosome proteins were analyzed using GO Term Mapper (34) (http://go.princeton.edu/cgi-bin/GOTermMapper) followed by manual curation to categorize proteins into groups of interest. All subsequent GO term analysis was performed using these *P. falciparum* homolog annotations. Comparison of GO metabolic process annotations to published liver and blood stage proteomes were performed using current GO term annotations for *P. falciparum* proteomic datasets (29–32) or data for syntenic *P. falciparum* orthologs if the original dataset was from a rodent parasite species (17).

Merozoite apical organelle proteins, merozoite surface proteins, merozoite proteases, and cytoskeleton or motility proteins were identified using one of the following resources: PlasmoDB, ApiLoc (http://apiloc.biochem.unimelb.edu.au/apiloc), MEROPS (https://www.ebi.ac.uk/merops/) and selected publications (18, 35–39). In each case, where localizations were inferred from studies in species other than *P. berghei*, proteins were included only if found to have one-to-one orthology to exclude potential issues with diverging protein functions in multigene families.

### Analysis of MSP4/5 Transcript Abundance and Splicing

Merosomes were cultured as above and blood stage schizont controls were produced as previously described (15). RNA was isolated from samples using a PureLink RNA Mini Kit (Invitrogen #12183018A) and DNA was removed by on-column DNAse digestion (Invitrogen #12185010). cDNA synthesis was performed using random hexamers (Invitrogen #N8080127) and MuLV reverse transcriptase (RT; Applied BioSystems #LSN8080018) with 100 ng RNA template per reaction. Samples without reverse transcriptase were also included to allow the presence of genomic DNA to be detected in the subsequent PCR reactions.

The relative total MSP4/5 transcript abundance in samples was measured by qPCR using primers against the first exon of the gene, which were previously shown to detect both spliced and unspliced forms of the transcript (40) (primers P1 5’-GAAAGCCGTAAATTACTTATCACTG-3’ and P2 5’-CCCTCATTTTGATTCGAACTAGTTG-3’). Relative abundance was calculated by comparison to the HSP70 transcript (primers 5’-TGCAGCAGATAATCAAACTC-3’ and 5’-ACTTCAATTTGTGGAACACC-3’). Spliced and unspliced forms of the MSP4/5 transcript were detected by PCR using primers that spanned the intron of the gene (primers P3 5’-GATAAAGCTGGAAGTGCTTC-3’ and P4 5’-ATCATCATCTTCATCATCTTCAG-3’). HSP70 primers were used as a control for total starting RNA amount. Controls lacking reverse transcriptase were used to allow for the presence of genomic DNA contamination to be excluded.

### Identification of Proteins with N-terminally Processed PEXEL Motifs

The identification of merosome proteins with N-terminally processed protein export elements (PEXELs) was based on a previously described method (41). Output files from mass spectrometry analysis were converted to mzML format using msConvert version 3.0.6002 (42) and searched with Comet version 2015.02 rev.0 (43). Spectra were searched against a database comprising *P. berghei* proteins [PlasmoDB v.33; (27)], *H. sapiens* proteins [UniRef reviewed reference proteome (44), proteins with 90% or greater redundancy collapsed into single entries], and the common Repository of Adventitious Proteins v.2012.01.01 (www.thegpm.org/cRAP). Decoy proteins with the residues between tryptic residues randomly shuffled were interleaved among the real entries using a tool in the Trans-Proteomic Pipeline (TPP) version 5.0.0 Typhoon (45). Precursor mass tolerance was ±10 ppm, fragment ions bins were set to a tolerance of 0.02 *m/z* and a monoisotopic mass offset of 0.0 *m/z*, and the use of flanking peaks for theoretical fragment ions was enabled. Semi-tryptic peptides and up to two missed cleavages were allowed. Search parameters included a fixed modification of +57.021464 Da at Cys for formation of S-carboxamidomethyl-Cys by iodoacetamide, and variable modifications of +15.994915 Da at Met for oxidation, +17.026549 Da at peptide N-terminal Gln for deamidation from formation of pyroGlu, +18.010565 Da at peptide N-terminal Glu for loss of water from formation of pyroGlu, −17.026549 Da at peptide N-terminal Cys for deamidation from formation of cyclized N-terminal S-carboxamidomethyl Cys, and +42.010565 Da at Lys, at peptide N-termini, and at protein N-termini (either at N-terminal Met or the N-terminal residue after cleavage of N-terminal Met) for acetylation. The MS/MS data were analyzed using the TPP, and peptide spectrum matches (PSMs) were assigned scores in PeptideProphet (46), employing accurate mass modeling to correct for systematic mass error and using the Comet expect score as the only contributor to the f-value for modeling. Localization of variable modifications was confirmed and/or corrected using a development version of PTMProphet (source code available at https://sourceforge.net/p/sashimi, SVN revision number 7605). PSMs identifying decoy proteins were assigned probabilities and used to estimate the FDR at a given PeptideProphet probability.

Putatively processed PEXEL motifs were identified by selecting peptides bearing N-terminal acetylation after being cleaved C-terminal to a Leu or Ile residue. Only PSMs identified at a PeptideProphet probability corresponding to an FDR of 0.0% among all such cleaved and acetylated peptides were taken for further analysis. (PeptideProphet probability cut-offs corresponded to FDRs of 0.21%, 0.12% and 0.08% among all PSMs in replicates 1, 2 and 3, respectively). Cleaved and acetylated peptides were then examined in the context of the originating protein for a canonical PEXEL motif, i.e. [K/R]x[L/I]x[D/E/Q] with cleavage between [K/R]x[L/I] and x[D/E/Q] and N-terminal acetylation of the resulting semi-tryptic peptide beginning with x[D/E/Q]. Alternative PEXELs of [K/R]x[L/I]xx were also considered to allow for the evolutionary divergence of *P. berghei* and *P. falciparum* motifs (47). Evidence for a cleaved and acetylated PEXEL was considered "Strong" if it was identified in all three independent replicates, "Fair" if it was identified by multiple PSMs in at least one replicate, and "Weak" if it was identified from at most a single PSM in any replicate (Supp Table 4). Proteins with “Strong” and “Fair” evidence for N-terminally processed PEXELs were considered confident identifications. These proteins were analyzed for the presence signal peptides or signal anchors using Signal P (48) and TargetP (49).

## Results

### Merosome Production and Quality Assessment

To produce sufficient quantities of merosomes for proteomics, the routine method for *P. berghei* liver stage culture was scaled up and modified to reduce potential contaminants. Quality and intactness of the cultured merosomes was assessed using light and immunofluorescence microscopy. Live light microscopy indicated samples contained a mixture of merosomes and detached cells as previously described (10), and indicated the majority of merosomes were spherical and intact. Immunofluorescence microscopy using antibodies specific for MSP1 identified individual hepatic merozoites within merosomes, and staining for the PV marker UIS4 indicated the vacuole membrane was fragmented or absent, confirming PV rupture had occurred prior to merosome formation (10) (Fig 1A).

Three independent replicates were prepared for proteomic analysis. Replicates one and two contained approximately 100,000 merosomes, and replicate three over 250,000 merosomes (Fig 1B). As merosomes may be of variable size and contain variable numbers of merozoites (10), we additionally quantified the number of hepatic merozoite genomes by qPCR. All replicates contained similar numbers of merozoite genomes (Fig 1B), suggesting the higher merosome count for replicate three was balanced by a smaller number of merozoites per merosome in that sample.

### The Merosome Proteome

As expected, untargeted proteomic analysis of merosome samples identified proteins from *P. berghei*, as well as from the host hepatocyte, mosquito vector, media components, and common contaminant proteins (Supp Table 1). Focusing our analysis on proteins of parasite origin, we identified an average of 15,159 *P. berghei* PSMs per replicate and 1,700 protein identifications per replicate (Fig 1B). Across all three replicates, we obtained a total of 1879 proteins identified by at least one peptide (Supp Table 1). To define the core protein identifications in this set, we considered only those that were represented by at least two unique peptides in every biological replicate. We observed excellent concordance between replicates in terms of protein abundance (Pearson correlation between 0.89 and 0.95; Fig 1C). Similarly, we observed strong concordance between replicates in terms of protein identifications, with over 1331 proteins identified by at least two unique peptides in each replicate, and a core set of 1188 proteins identified by at least two peptides in every replicate (Fig 1B and D). We define these 1188 proteins as the core *P. berghei* merosome proteome (Supp Table 1).

**Figure 1:**
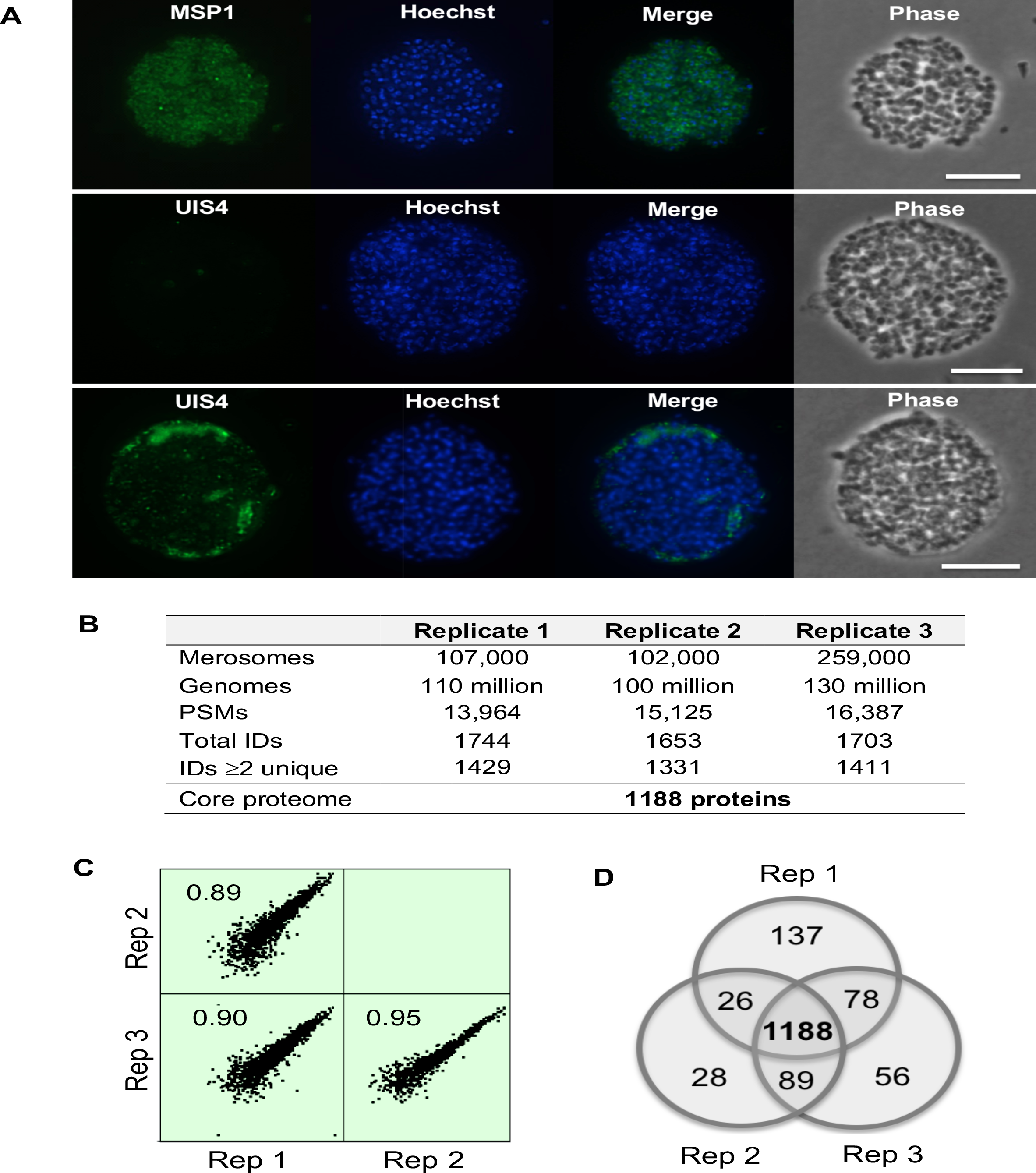
Overview of the core *P. berghei* merosome proteome. A. Immunofluorescence analysis of merosome samples used for proteomics. Top: Staining with antibodies specific for MSP1 identifies hepatic merozoites within merosomes and confirms merosomes are intact. Middle and bottom: Staining with antibodies specific for the PV marker UIS4 demonstrates the vacuole is absent or fragmented in merosomes. DNA stained with Hoechst. Scale bar 10 microns. B. Overview of the *P. berghei* merosome proteome. Genomes refer to the number of hepatic merozoite genomes estimated by quantitative PCR. PSMs refers to peptide spectrum matches. Total IDs are proteins identified by at least one peptide. IDs ≥ 2 unique are proteins identified by at least two unique peptides. C. Correlation among biological replicates. iBAQ values were log_2_ transformed to generate scatter plots showing the correlation among biological replicates. The Pearson correlation between replicates is indicated in each plot. Data shown are for proteins with >2 unique peptides and plots were generated using Perseus version 1.6.0.2078. D. Concordance between protein IDs with ≥ 2 unique peptides between replicates. We define the core merosome proteome as the 1188 proteins identified by ≥ 2 unique peptides in all three biological replicates.

To gain information about the relative abundance of proteins in the merosome proteome, we used intensity-based absolute quantification (iBAQ) to rank proteins (Supp Table 1). The most abundant proteins in the merosome proteome included merozoite surface protein 1, elongation factor alpha, actin, and common dual-function enzymes such as GAPDH and enolase (50). The PV resident protein exported protein 1 (EXP1) was also among this list, consistent with the observed presence of vacuole membrane fragments in some merosomes (Fig 1A). Other abundant proteins included ribosome components, histones, and heat shock proteins, as previously reported for other *Plasmodium* proteomes and transcript analyses (28, 51).

The core merosome proteome corresponds to 24% of predicted protein coding sequences in the *P. berghei* genome (27). This depth of coverage is similar or superior to previously published untargeted proteomes of other *Plasmodium* life stages (17, 18, 41, 52). Approximately 81% of merosome proteins have annotated protein descriptions, while the remaining 19% are hypothetical proteins or proteins of unknown function. Importantly, the vast majority of proteins in the proteome were found to have syntenic orthologs in *P. falciparum*, indicating the protein repertoire of merosomes is broadly conserved between rodent and human malaria parasite species.

### Global Comparison to Liver and Blood Stage Proteomes

Comparison of the core merosome proteome to previously published *Plasmodium* liver and blood stage proteomes reveals significant overlap with both of these stages (Fig 2A and Supp Table 2). Comparison to four different *P. falciparum* blood stage schizont and erythrocytic merozoite proteomes revealed 91.8% of merosome proteins were shared with at least one of these stages (29–32) corresponding to 1091 of the 1188 identified proteins in the core merosome proteome (Supp Table 2.1). The merosome proteome therefore bears very strong resemblance to published blood stage schizont and merozoite proteomes, consistent with observations that liver stage merozoites comprise most of the volume of merosomes (10) and that these merozoites closely mirror their blood stage counterparts.

Merosomes also share considerable similarity to the *P. yoelii* liver stage proteome, with 46.5% of merosome proteins previously identified in liver stage parasites (17), corresponding to 553 of the 1188 identified proteins in the core proteome (Fig 2A and Supp Table 2.2). It is important to note that since far fewer proteins were identified in the liver stage proteome (17), this comparison underestimates the true degree of similarity between liver stage parasites and merosomes. Indeed, of the liver stage proteome (n=664 with syntenic orthologs), 83.3% of these proteins are identified in our core merosome proteome, suggesting that when an updated liver stage proteome becomes available, the overlap will be in this range. Of note, we did observe a subset of proteins that were uniquely shared between liver stages and merosomes, which included known liver stage-specific proteins such as LISP1 and LISP2 (51, 53). The merosome proteome therefore likewise bears considerable similarity to liver stages, consistent with the continued presence of liver stage proteins in merosomes after PV breakdown and merosome budding from the host hepatocyte.

**Figure 2:**
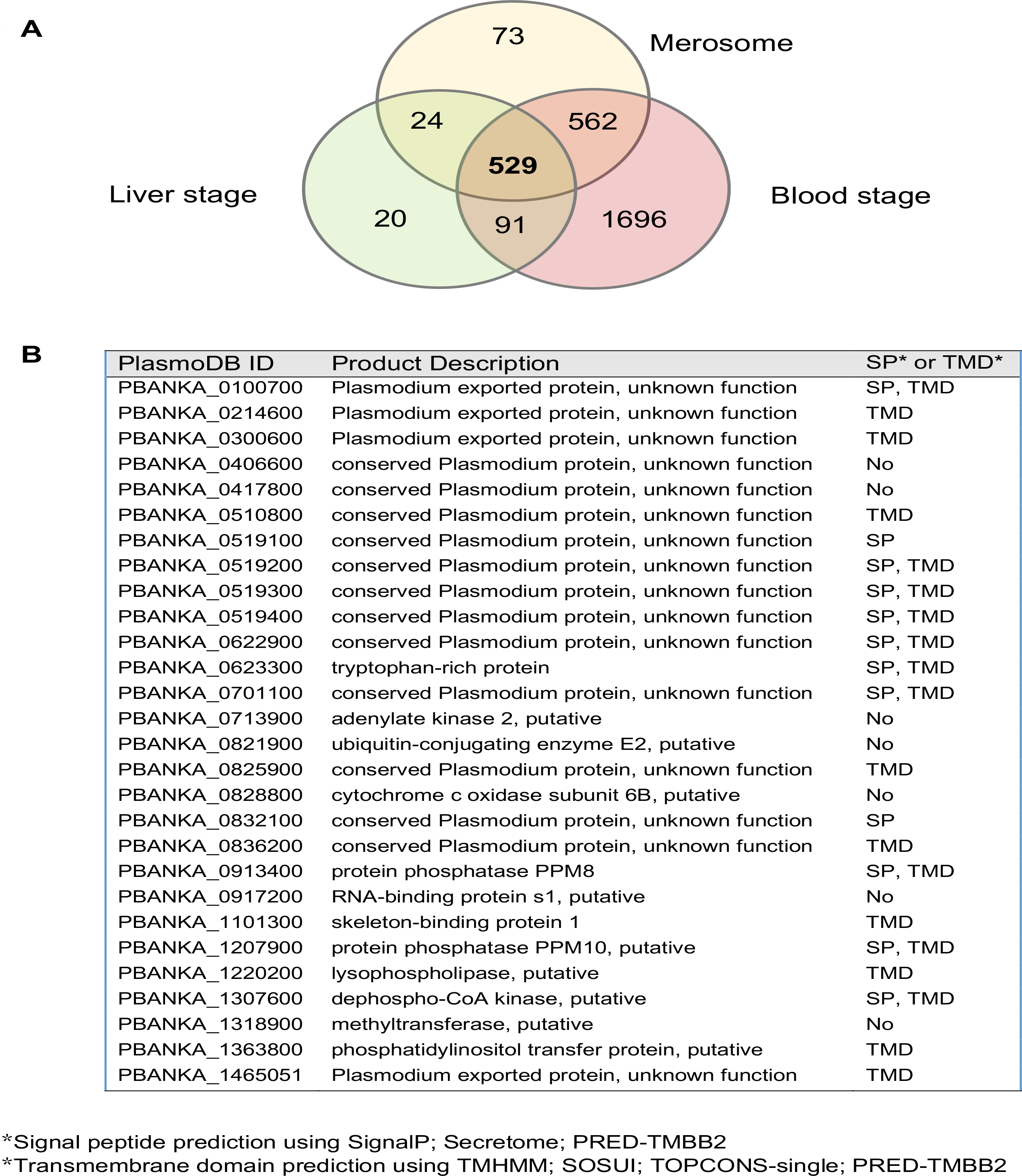
Overlap between the core merosome proteome and published liver and blood stage proteomes with potentially merosome unique proteins shown. A. Venn diagram depicting the overlap between our core merosome proteome, the *P. yoelii* liver stage proteome (17) and the combined proteomes of four *Plasmodium* blood stage schizont or merozoite proteomes (29–32). Proteins in the published proteomes were converted to their syntenic *P. berghei* orthologs for this comparison. The majority of merosome proteins have syntenic orthologs that have previously been detected in the liver or blood stage. B. Table of putative merosome unique proteins. The 73 proteins not overlapping with liver and blood stage proteomes were further analyzed: Proteins previously described in other life cycle stages or part of multigene families were removed and the remaining 28 proteins may be unique to merosomes. SP, signal peptide; TM, transmembrane domain.

To explore the biological processes represented by the proteins in this proteome, we combined GO term analysis with manual curation of the proteome, using GO terms for the *P. falciparum* orthologs because of the relative paucity of annotations for *P. berghei* (Supp Fig 1 and Supp Table 3.1). Consistent with the overall similarity of merosomes to liver and blood stage parasites, numerous metabolic pathways were shared between stages. Common processes in liver stage parasites (17) and merosomes included amino acid metabolism, carbohydrate metabolism, energy metabolism, cofactor metabolism, and lipid metabolism, most notably the fatty acid synthesis enzymes known to be upregulated in the liver stage relative to the blood stage (17, 54, 55). Similarly, most of the annotated merosome metabolic proteins were also shared with blood stage parasites (29–32). Therefore, at least for canonical metabolic pathways that are well annotated, merosomes share the majority of metabolic processes with liver and/or blood stage parasites.

### Shared Liver and Blood Stage Merozoite Proteins

#### Apical Organelle Proteins

To identify individual proteins shared between hepatic and erythrocytic merozoites, we searched the merosome proteome for proteins known to be involved in red blood cell invasion in erythrocytic merozoites. We identified forty-seven proteins that had previously been localized to the apical organelles (rhoptries or micronemes), dense granules, or merozoite apex in erythrocytic merozoites (Table 1). Examining this list, we observed similar abundance rankings of known complex-forming proteins RAP1 and RAP2/3, and RhopH2, RhopH3 and Clag, consistent with their predicted or known binding ratios in blood stage parasites (56, 57). We detected one highly abundant p235 reticulocyte binding protein and several lesser abundant p235s, mirroring findings from transcriptomic studies of these proteins in the blood stage (58). However, because p235 family members do not have strict one-to-one orthology across *Plasmodium* species, we could not determine if the most abundant merosome p235 protein was the closest ortholog of the reported *P. yoelii* hepatic merozoite-specific Py235 transcript (14). We also identified all five components of the *Plasmodium* Translocon of Exported Proteins (PTEX translocon) (59) (PTEX150, HSP101, EXP2, TRX2 and PTEX88), consistent with observed transfer of these components from the merozoite apical organelles to the developing PV upon red blood cell invasion (60, 61). Thus, the repertoire and relative abundance of known apical organelle proteins are largely conserved between hepatic and erythrocytic merozoites.

**Table 1:** Apical organelle and merozoite surface proteins in merosomes

| **PlasmoDB ID** | **Protein Description** | **Rank<sup>a</sup>** |
| --- | --- | --- |
| **Microneme** |  |  |
| PBANKA_0702800 | protein disulfide isomerase | 27 |
| PBANKA_0915000 | apical membrane antigen 1 (AMA1) | 136 |
| PBANKA_1332700 | duffy-binding protein | 395 |
| PBANKA_0701900 | GPI-anchored micronemal antigen (GAMA), putative | 550 |
| PBANKA_0933500 | rhomboid protease (ROM1) | 754 |
| PBANKA_1215100 | Rh5 interacting protein, putative | 774 |
| PBANKA_0514200 | claudin-like apicomplexan microneme protein, putative | 1096 |
| **Rhoptry Neck** |  |  |
| PBANKA_0501400 | rhoptry neck protein 12 (RON12), putative | 92 |
| PBANKA_0932000 | rhoptry neck protein 4 (RON4) | 156 |
| PBANKA_0100400 | reticulocyte binding protein (Pb235), putative | 209 |
| PBANKA_0713100 | rhoptry neck protein 5 (RON5), putative | 261 |
| PBANKA_1315700 | rhoptry neck protein 2 (RON2) | 320 |
| PBANKA_0311700 | rhoptry neck protein 6 (RON6), putative | 519 |
| PBANKA_1003600 | apical sushi protein (ASP), putative | 709 |
| PBANKA_0721800 | apical merozoite protein (Pf34), putative | 740 |
| PBANKA_1464900 | rhoptry neck protein 3 (RON3), putative | 741 |
| PBANKA_0216761 | reticulocyte binding protein (Pb235), putative | 982 |
| PBANKA_1400091 | reticulocyte binding protein (Pb235), putative | 1131 |
| PBANKA_0316600 | reticulocyte binding protein (Pb235), putative | 1154 |
| **Rhoptry** |  |  |
| PBANKA_1032100 | rhoptry-associated protein 1 (RAP1), putative | 53 |
| PBANKA_1101400 | rhoptry-associated protein 2/3 (RAP2/3) | 55 |
| PBANKA_0919100 | parasitophorous vacuolar protein 1 (PV1), putative | 62 |
| PBANKA_0830200 | high molecular weight rhoptry protein 2 (RhopH2), putative | 150 |
| PBANKA_0416000 | high molecular weight rhoptry protein 3 (RhopH3), putative | 161 |
| PBANKA_1400600 | cytoadherence linked asexual protein (Clag), putative | 191 |
| PBANKA_1210600 | rhoptry associated adhesin, putative | 256 |
| PBANKA_0804500 | rhoptry-associated membrane antigen (RAMA), putative | 283 |
| PBANKA_0619700 | rhoptry-associated leucine zipper-like protein 1 (RALP1), putative | 288 |
| PBANKA_0716900 | armadillo-domain containing rhoptry protein, putative | 301 |

| PlasmoDB ID | Protein Description | Rank<sup>a</sup> |
| --- | --- | --- |
| **Rhoptry (continued)** |  |  |
| PBANKA_0813000 | inhibitor of cysteine proteases | 563 |
| PBANKA_0111600 | rhoptry protein ROP14, putative (ROP14) | 996 |
| PBANKA_1330300 | rhoptry protein, putative | 1147 |
| **Dense Granule** |  |  |
| PBANKA_1008500 | translocon component PTEX150 (PTEX150) | 196 |
| PBANKA_0931200 | heat shock protein 101 (HSP101) | 249 |
| PBANKA_1334300 | exported protein 2 (EXP2) | 264 |
| PBANKA_0911700 | subtilisin-like protease 2 (SUB2) | 977 |
| **Apical Region** |  |  |
| PBANKA_1002600 | 6-cysteine protein (p41) | 218 |
| PBANKA_1107600 | 6-cysteine protein (p38) | 316 |
| PBANKA_1358000 | thioredoxin 2 (TRX2) | 388 |
| PBANKA_0501200 | early transcribed membrane protein (UIS4) | 493 |
| PBANKA_0941300 | translocon component PTEX88 | 593 |
| PBANKA_1433600 | thrombospondin-related apical membrane protein (TRAMP) | 766 |
| PBANKA_1400800 | early transcribed membrane protein (UIS3) | 811 |
| PBANKA_1107100 | subtilisin-like protease 1 (SUB1) | 805 |
| PBANKA_0934500 | protein tyrosine phosphatase | 904 |
| PBANKA_0209100 | thrombospondin related sporozoite protein (TRSP) | 958 |
| PBANKA_1321700 | berghepain-1 (BP1) | 1078 |
| **Merozoite Surface** |  |  |
| PBANKA_0831000 | merozoite surface protein 1 (MSP1) | 14 |
| PBANKA_1349000 | MSP7-like protein (MSRP2) | 21 |
| PBANKA_1349100 | MSP7-like protein (MSP7) | 41 |
| PBANKA_1102200 | merozoite surface protein 8 (MSP8) | 159 |
| PBANKA_1002600 | 6-cysteine protein (P41) | 218 |
| PBANKA_0111000 | 6-cysteine protein (P12) | 250 |
| PBANKA_1107600 | 6-cysteine protein (P38) | 316 |
| PBANKA_1119600 | merozoite surface protein 10, putative (MSP10) | 721 |

<sup>a</sup> Intensity-based absolute quantification (iBAQ) ranking

A comparison with the available parasite organelle-enriched proteomes indicated merosomes likewise share significant overlap with these datasets (Supp Table 2.4, 2.5 and 2.6). Merosomes contained approximately 70% of the 36 proteins identified in the rhoptry-enriched fraction of blood stage parasites (28), and a similar fraction of the ~300 proteins identified in the microneme-enriched fraction ookinetes (62), the motile stage that infects the mosquito midgut. Merosomes also had orthologs of many proteins identified in *P. falciparum* blood stage extracellular vesicle fraction, which likely includes both apical organelle and secreted proteins (33). While only a subset of these proteins have had their localizations experimentally confirmed (63), these findings nonetheless suggest that hepatic merozoites may share tens to hundreds of apical organelle or co-fractionating proteins with erythrocytic merozoites and motile stages more broadly.

#### Cytoskeleton, Motility and Invasion Proteins

To identify cytoskeleton and motility proteins shared between hepatic and erythrocytic merozoites, we searched the merosome proteome for proteins known to be involved in these processes (Supp Table 3.2). As observed in blood stage parasites (29), we detected key components of the actin-myosin motor such as actin, myosin A, and myosin A tail domain interacting protein (MTIP) in considerable abundance, ranked 13^th^, 84^th^ and 198^th^ in merosomes, respectively. Merosomes possessed numerous other important motility-related and cytoskeletal components such as actin depolymerizing factor 1, profilin, aldolase, GAP45, GAPM1, GAPM2, GAPM3, and tubulin (35, 36). The actin nucleation protein formin 1 (64) was also detected in two of three merosome replicates, suggesting it may be present at levels near the limit of detection (Supp Table 1).

#### Merozoite Surface Proteins and Proteases

To identify merozoite surface proteins shared between hepatic and erythrocytic merozoites, we searched the merosome proteome for proteins known to be surface-localized in erythrocytic merozoites (Table 1). The known merozoite surface proteins (MSPs) found in merosomes were MSP1, MSP7-like proteins, MSP8, MSP9, MSP10 and the 6-cysteine family members p12, p38 and p41 (37, 38). Consistent with findings from blood stage parasites (65), MSP1 and MSP7-like proteins were among the most abundant proteins in merosomes, ranked 14^th^, 21^st^ and 41^st^ respectively. As proteolytic processing of surface and secreted proteins plays a critical role in egress and invasion (38), we also searched the merosome proteome for proteases (Supp Table 3.3). We observed several proteases known to have roles during merozoite priming, egress and invasion, including members of the serine repeat antigen (SERA) family, the subtilisin-like (SUB) protease family, and the rhomboid (ROM) protease family (38). Thus, for most characterized merozoite surface proteins and proteases, hepatic and erythrocytic merozoites appear highly similar.

### Unique Features of Merosomes and Liver Stage Merozoites

#### Putatively Unique Merosome Proteins

While comparison to the published liver and blood stage proteomes suggests that the majority of merosome proteins are shared with one or both of these stages (17, 29–32), it also identified a small subset of proteins that had not previously been detected or did not have syntenic orthologs in these liver or blood stage proteomes. Specifically, we found 6.1% of the merosome proteome or 73 proteins in this group (Fig 2A and Supp Table 2.3). Closer examination of this list reveals that only a subset of these proteins is likely to be unique to merosomes. Indeed, several proteins are members of multigene families that do not have strict one-to-one orthology across *Plasmodium* species, such as the p235 proteins, duffy binding proteins, and fam-a or fam-c proteins (Supp Table 2.3). These proteins are therefore not likely unique to merosomes, but were identified because we considered only proteins with clear orthologs in our global comparisons. Another group are liver stage proteins whose orthologs were not observed in the only other published proteomic analysis of liver stages, performed on mid-and late-stage *P. yoelii* liver stage forms (17). As discussed, fewer proteins were identified in that work compared to our proteome, in part because of interference from host proteins, and in part because the mass spectrometers available at the time were more limited in sensitivity, duty cycle, and resolution than the instrument used in our study. In light of this, many of the putatively unique merosome proteins we identify may also turn out to be present in liver stages if such a study were re-visited with modern instrumentation. Finally, this subset also includes proteins previously detected in sporozoites (18, 19), or organelle-fraction proteomes of various life cycle stages (62, 63). Given these many caveats, we have manually annotated the list of potentially unique merosome proteins to highlight those 28 proteins that may be unique to merosomes versus those that were prematurely labeled as unique for one of the reasons outlined above (Fig 2B and Supp Table 2.3). We anticipate that future studies will confirm some of these proteins to be truly unique to merosomes.

#### Abundant Hypothetical Proteins

Examining the merosome proteome as a whole, we noted several hypothetical proteins were among the most abundant proteins in our dataset (Supp Table 1). Since we suspected the unique biology of merosomes may be accompanied by unique or divergent proteins, we undertook further analysis of these proteins. The top 8 most abundant hypothetical proteins were selected for analysis, as all were ranked among the top 100 proteins in merosomes (Supp Table 1). Examination of available expression data for these proteins showed evidence for expression at multiple other life stages for all except the most abundant, suggesting the majority do not have unique roles in the liver stage or merosomes (Supp Table 3.4).

Nonetheless, the most abundant hypothetical protein (PBANKA_0518900) was noteworthy because it appears to be restricted to the late liver stage and merosomes based on available expression data. Clear homologs of the protein are found only in rodent malaria parasites, so less data is available than for the proteins that have *P. falciparum* orthologs. Nevertheless, the available RNA-seq data indicate little or no expression in blood stage parasites [PlasmoDB.org; (66, 67)], and the protein is conspicuously absent from the *P. berghei* blood stage proteome (52), despite being detected in the *P. yoelii* liver stage proteome (17). Furthermore, our data indicate the protein is more abundant in merosomes than the abundant merozoite proteins MSP1 and actin (Supp Table 1). As this hypothetical protein does not have any annotated domains or genetic modification data to date, it is difficult to speculate its possible function in merosomes. However, its abundance in merosomes and restriction to the liver stage suggest this would be a good candidate for further study.

#### Differential Expression of Merozoite Surface Protein 4/5

While searching for known merozoite proteins in merosomes allowed us to identify numerous common hepatic and erythrocytic merozoite proteins, there was a notable absence of peptides derived from MSP4/5 detected in merosomes. In blood stage parasites, MSP4/5 expression is known to be regulated by alternative splicing, with an unspliced transcript detected in early blood stages when no MSP4/5 protein is detected, but a spliced transcript detected in mature blood stages where MSP4/5 protein is normally expressed (68). To explore if alternative splicing was responsible for the lack of detection of MSP4/5 in merosomes and hepatic merozoites, we performed quantitative reverse transcriptase PCR to quantify total MSP4/5 transcript (Fig 3A), using blood stage schizonts as a control. We found total MSP4/5 transcript abundance was comparable in schizonts and merosomes (Fig 3B), supporting the hypothesis that MSP4/5 protein expression was translationally regulated.

We next performed reverse transcriptase PCR using primers that bound either side of the alternatively spliced intron, allowing us to visualize MSP4/5 mRNA splice variants present at each stage. We detected both the spliced and unspliced mRNA forms in schizonts, but only the unspliced form in merosomes, confirming the transcript was differentially spliced between these stages (Fig 3C). As the unspliced form contains a premature stop codon and does not produce detectable protein in early blood stage parasites (68), we conclude the absence of MSP4/5 in merosomes is likewise due to the lack of a functional spliced transcript. Thus, we identify differential expression of MSP4/5 as the third known difference between hepatic and erythrocytic stage merozoites (Table 2).

**Figure 3.**
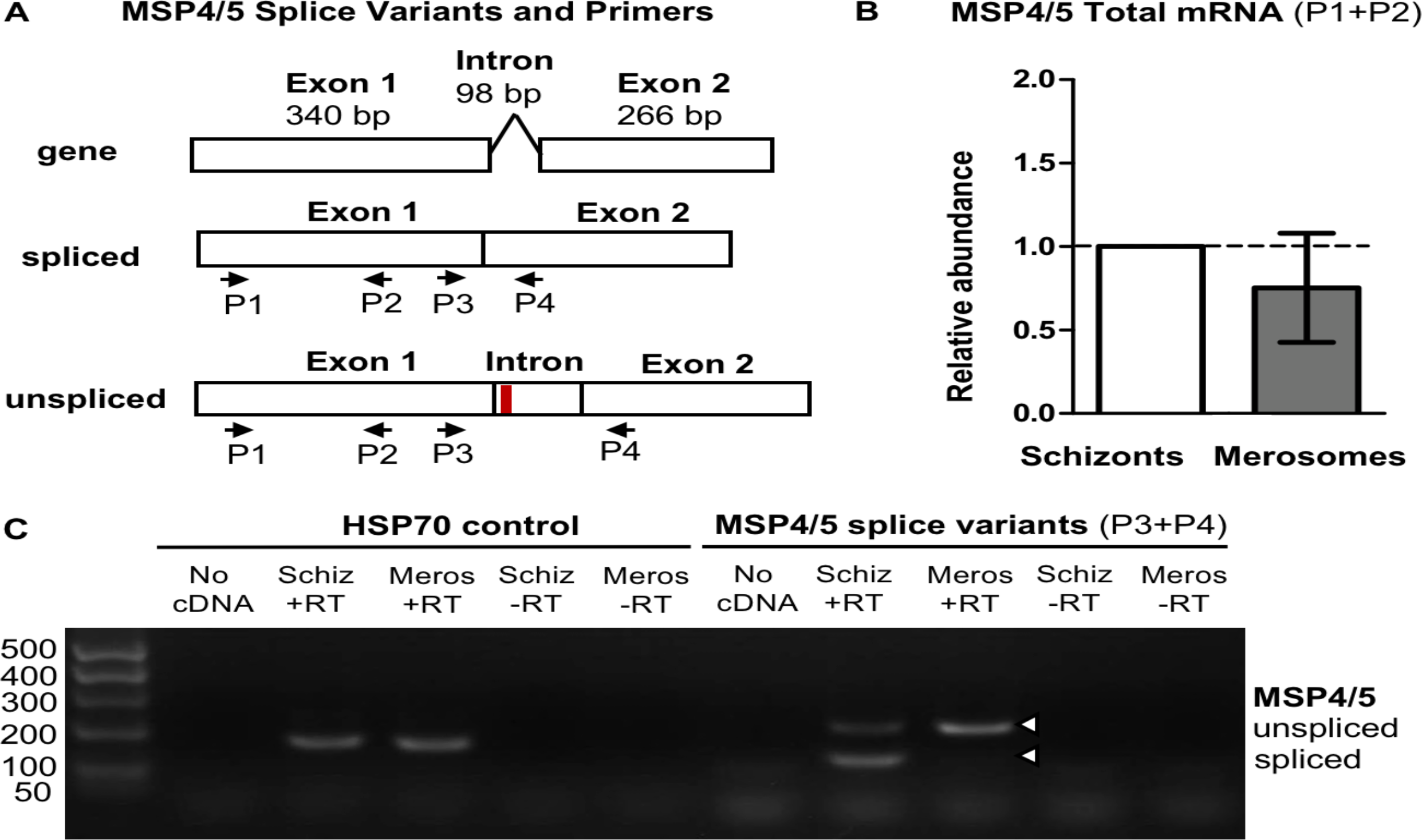
The absence of merozoite surface protein 4/5 in merosomes is associated with alternative splicing of the mRNA transcript. A. Diagram of the *P. berghei* MSP4/5 gene structure, spliced mRNA, and unspliced mRNA. Unspliced mRNA contains a premature stop codon (red line). Primers used to quantify total MSP4/5 mRNA (P1+P2) and detect splice variants (P3+P4) are shown. B. Quantitative reverse transcriptase PCR demonstrates relative abundance of total MSP4/5 mRNA is comparable in blood stage schizonts and merosomes. Relative abundance calculated by comparison to HSP70. Relative abundance of MSP4/5 was normalized to 1.0 for schizonts. Error bars show mean ± standard deviation. C. Non-quantitative reverse transcriptase PCR demonstrates MSP4/5 is alternatively spliced in schizonts and merosomes. Abbreviations: Schiz, schizonts; Meros, merosomes; +RT, with reverse transcriptase; -RT, without reverse transcriptase.

**Table 2.** Known differences between hepatic and erythrocytic merozoites

| PlasmoDB ID | Protein | Difference | Ref |
| --- | --- | --- | --- |
| PBANKA_0304400 | MSP4/5 | Protein absent from hepatic merozoites | This work |
| PBANKA_1321700 | berghepain-1 | Knockout has more impact on hepatic merozoites | (15) |
| Multiple | Py235 family | Different members in hepatic vs erythrocytic merozoites | (14) |

#### Putatively Exported Proteins

Finally, since the existence of a functional PTEX complex in liver stages remains somewhat controversial, we explored whether the proteome of intact merosomes might contain proteins that were exported into the host hepatocyte prior to merosome release. There is evidence that the PTEX translocon may be involved in protein export in the liver stage, either for trafficking proteins to the PV, or beyond the PV into the host hepatocyte (51). Protein export via the PTEX has been extensively studied in blood stage parasites, and is commonly mediated by a pentameric amino acid sequence called a PEXEL motif (59). PEXEL motifs are located downstream of a signal sequence, and after cleavage by Plasmepsin V and N-terminal acetylation in the ER, are sufficient to mediate protein export to the host cell (69–72). To investigate PEXEL mediated export in the liver stage, and to search for exported proteins that remain in remnant host cell cytoplasm of mature merosomes, we searched our proteome for cleaved, N-terminally acetylated PEXEL peptides using the parameters previously described (41).

We confidently identified processed PEXEL motifs for nine merosome proteins, and detected single PSMs corresponding to processed motifs for a further eight (Table 3 and Supp Table 4). Annotated MS/MS spectra of cleaved and acetylated PEXEL motifs can be found in Supplementary Figure 2. The confidently identified proteins included the characterized liver stage-specific exported protein LISP2, which localizes to the host hepatocyte cytosol (51), and three known dual blood and liver stage exported proteins IBIS1, SMAC and UIS2, which localize either to the liver stage PV or the associated tubulovesicular network (73–75). The list also included two proteins previously shown be exported to the PV or host cell by blood stage parasites, but had not yet been studied in liver stage parasites, namely the StAR-related lipid transport protein and a PHIST domain protein (22, 76–79). The remaining three proteins had previously been predicted to be exported, annotated as putatively exported proteins of unknown function (PBANKA_1400700 and PBANKA_0700700), and a putatively exported fam-c domain protein (PBANKA_1465000) (66). These findings support the hypothesis that exported proteins can be retained in the remnant host cell cytoplasm after PV breakdown and merosome release, and suggests the as-yet-uncharacterized PEXEL-containing proteins may likewise be exported to the PV or host cell in the liver stage.

**Table 3:**
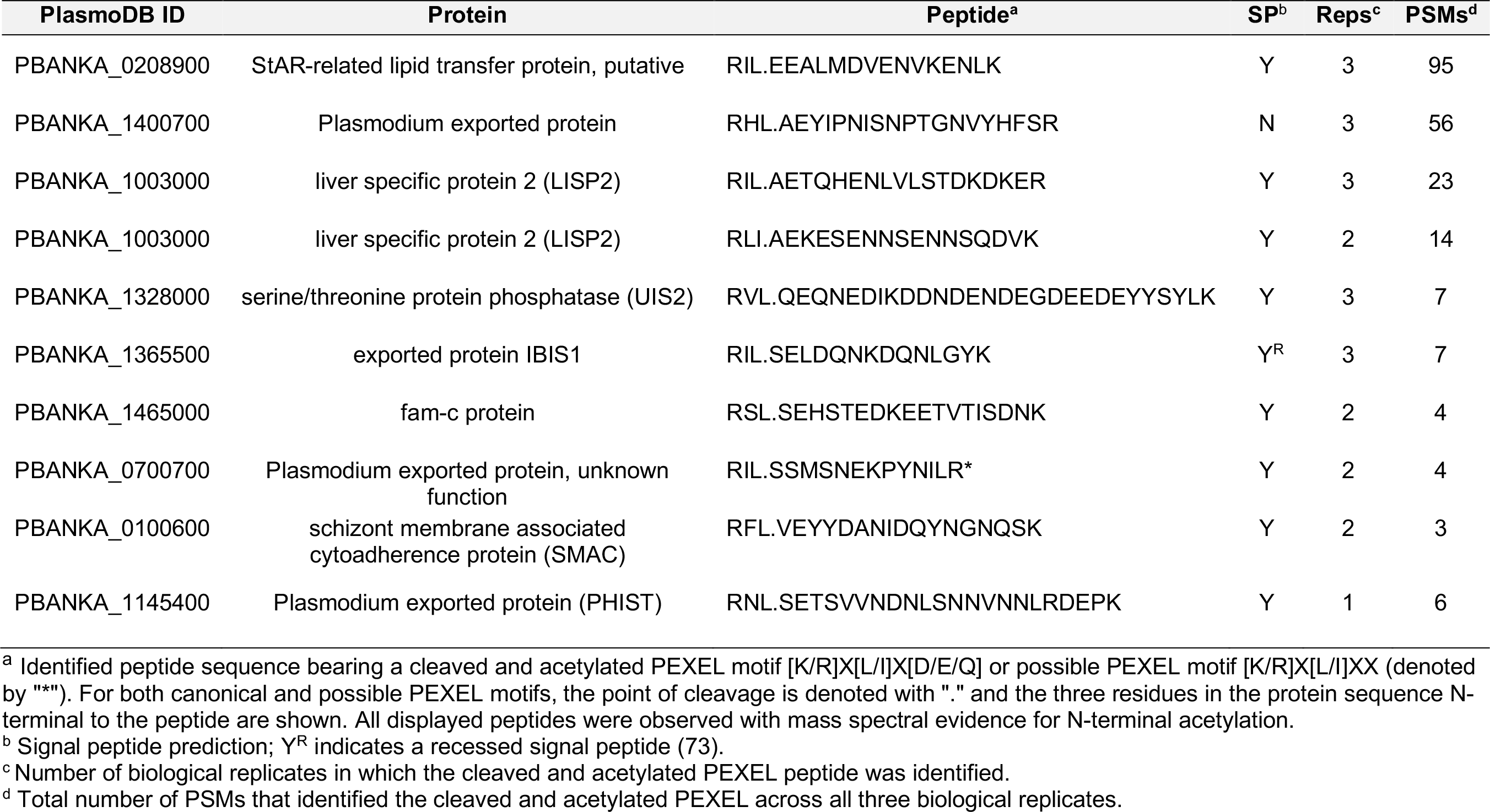
Proteins with processed PEXEL protein export motifs

Analysis of this group of proteins revealed several findings consistent with the known mechanism of PEXEL-mediated protein export. We observed all but one protein had a predicted signal peptide or signal anchor, and in most cases the PEXEL motif matched the canonical ‘RxLxE’ consensus (59) (Table 3 and Supp Table 4). We also found that most of the PEXEL motifs were positioned in proximity to the signal sequence as expected (59). The exception was LISP2, a large protein in which the putative PEXEL motif RILAE appears once and a second putative PEXEL motif RLIAE appears sixteen times dispersed throughout a repeat region. In addition to finding strong evidence for processing of the single RILAE motif, we also detected processing of the RLIAE motif as the peptide n_42_AEKESENNSENNSQDVK (n_42_ indicates where the PEXEL motif was cleaved and the protein was N-terminally acetylated). This exact peptide sequence is repeated nine different times in the protein, so it was not possible to determine if the protein in the sample was processed at a single site or multiple sites. However, we speculate this multiplicity of PEXEL cleavage sites in LISP2 may explain the atypical processing previously reported for this protein (51). Consistent with these data, we identified both Plasmepsin V and the putative N-acetyltransferase in merosomes (Supp Table 1), confirming the enzymes responsible for PEXEL processing (80) were present at this stage.

To further validate these findings, we localized the PHIST protein (PBANKA_1145400) by confocal immunofluorescence microscopy in late liver stage parasites and merosomes. A previous study found this protein was highly expressed in *P. berghei* blood stage schizonts, and demonstrated it was exported into the host red blood cell cytoplasm (22). In late liver stage parasites, we observed punctate staining for the PHIST protein that partially colocalized with the PV marker UIS4, suggesting it was likewise exported at this stage (Fig 4A). To better quantify this partial colocalization, we performed 3D image analysis of z-stacks and calculated the percent of PHIST positive pixels that were positive for UIS4. Analysis of 75 individual parasites indicated over half of all PHIST positive pixels were UIS4 positive (mean colocalization 56.0% ± 15.1 standard deviation). Thus, in late liver stage parasites a large fraction of the PHIST protein is trafficked to the PV membrane, consistent with PEXEL-mediated export of the protein.

In merosomes, where the PV membrane is absent, we observed PHIST staining associated with individual merozoites (Fig 4B). This finding was further supported by the pattern of PHIST staining in free hepatic merozoites, where much of the “teardrop” shaped merozoite was stained (Fig 4C). Thus, it appears that the localization of the PHIST protein changes during liver stage development, being exported to puncta at or near the PV in late liver stages, then localizing on or within individual merozoites after merosomes are formed.

**Figure 4:**
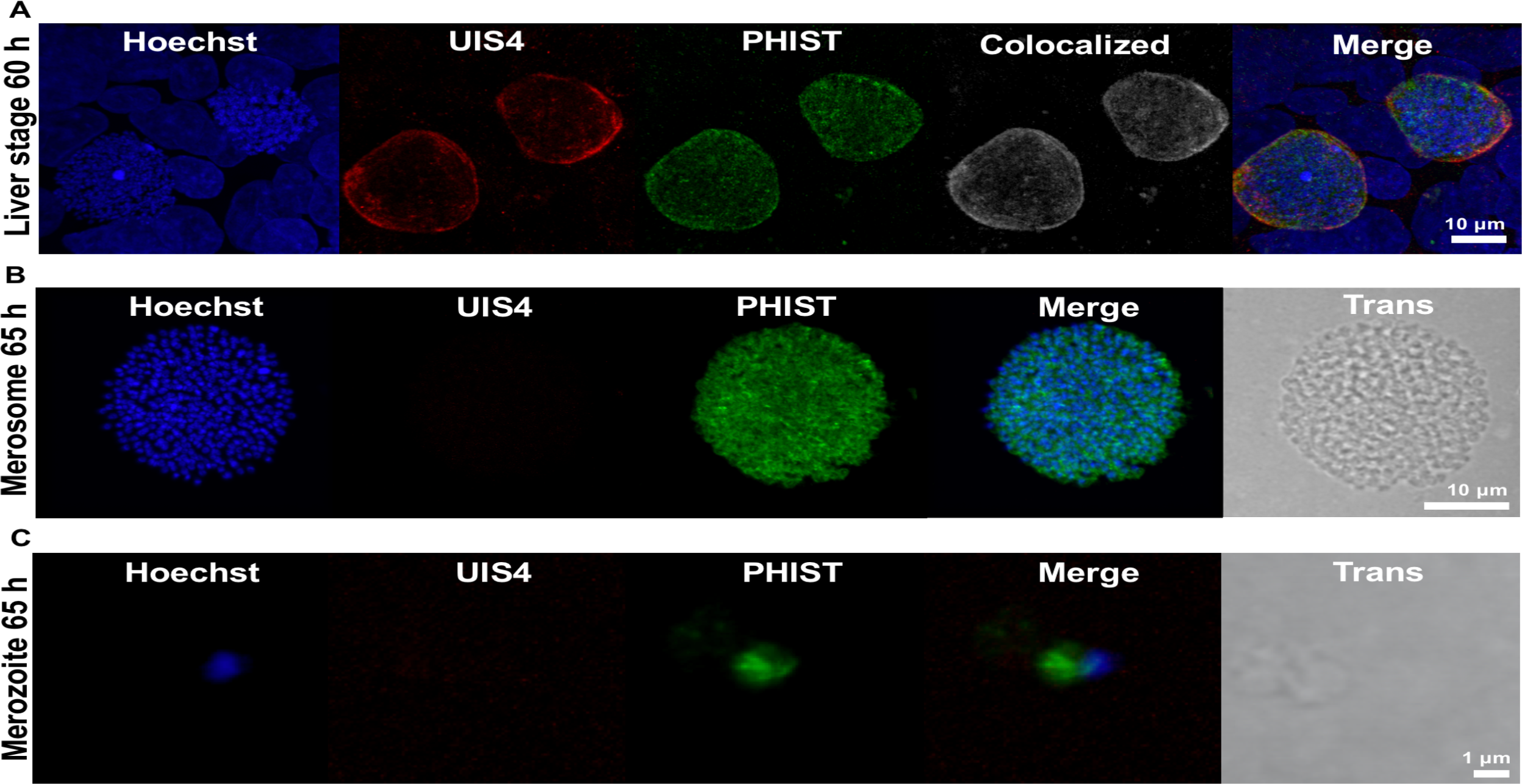
Localization of the PHIST protein in late liver stage parasites and merosomes. A. Immunofluorescence staining of late liver stage parasites at 60 h post infection indicates the PHIST protein (PBANKA_1145400) partially colocalizes with the parasitophorous vacuole marker UIS4. Colocalized pixels in white. DNA stained with Hoechst. B and C. Immunofluorescence staining of merosomes and free merozoites at 65 h post infection demonstrates the PHIST protein associates with individual liver stage merozoites.

## Discussion

We describe the first proteomic analysis of *Plasmodium* merosomes, the stage that initiates blood stage infection. Due to the relatively low numbers of hepatic merozoites and greater barrier to accessibility in comparison to other *Plasmodium* life cycle stages, few studies of merosomes and hepatic merozoites have been performed (16). Nonetheless, sensitive mass spectrometry techniques together with improvements to culture techniques have enabled us to identify a total of 1879 *P. berghei* proteins from merosomes, and a core set of 1188 proteins quantitatively detected across all replicates, producing the most comprehensive late liver stage dataset to date.

Comparison of the merosome proteome to published *Plasmodium* liver and blood stage proteomes demonstrates that merosomes share considerable similarity to both liver and blood stages. Considering merosomes are the final stage of liver stage development and the initial stage of infection in the blood, this overlap is consistent with merosomes being poised at this nexus. The similarity of merosomes to blood stages points to similar structural and functional features of hepatic and erythrocytic merozoites. Indeed, hepatic and erythrocytic merozoites were found to share the vast majority of known apical organelle proteins, cytoskeleton and motility proteins, and surface proteins. Merosomes also share considerable similarity to liver stage parasites, sharing known liver stage-specific proteins and a number of hypothetical proteins. Interestingly, this includes the highly abundant hypothetical protein (PBANKA_0518900), specific to liver stage parasites and whose function remains unknown. Comparison of the merosome proteome to previously published proteomes further identified a subset of proteins potentially unique to merosomes. Though a proportion of the proteins in this group would be expected to be liver stage proteins that were not identified in the earlier liver stage proteome (17), some are likely to be bona fide unique hepatic merozoite or merosome proteins, and are high value targets for follow-up studies.

Despite the overall similarity between erythrocytic and hepatic merozoites, previous studies focusing on particular proteins or protein families have demonstrated some differences between these two populations of merozoites. *P. yoelii* hepatic merozoites were previously reported to express a unique Py235 rhoptry protein (14) and we had previously found that hepatic merozoites have a greater requirement for the cysteine protease berghepain-1 compared to erythrocytic merozoites (15). Here we identify differential expression of MSP4/5 as a third difference between these two merozoite populations. We found MSP4/5 was absent from merosomes demonstrating that it is not expressed in hepatic merozoites, in contrast to erythrocytic merozoites where the protein is readily detected (68). As reported for early blood stage parasites (68), lack of the MSP4/5 protein in merosomes correlated with alternative splicing of the transcript, suggesting a common mechanism of translational regulation is conserved across both life stages. Why MSP4/5 might be expressed in erythrocytic merozoites but not hepatic merozoites is currently unclear, as its function in blood stage parasites is not known (38). Since MSP4/5 has been pursued as a vaccine antigen (81–83) and vaccines targeting both hepatic and erythrocytic merozoites would likely be more effective, it would be important to determine if the orthologous proteins are likewise differentially expressed in *P. falciparum*.

Our findings also support a role for the PTEX translocon in liver stage parasites. The identification of nine proteins with cleaved acetylated PEXEL motifs and concurrent detection of the enzymes responsible for their cleavage suggests that PEXEL processing in liver stage parasites can occur as it does in blood stages. Furthermore, the discovery of processed PEXELs in known liver stage exported proteins argues that this processing is relevant to their trafficking. One of the exported proteins we identified is the PHIST protein, PBANKA_1145400. *P. berghei* has two PHIST-domain proteins, and a previous study charactering these proteins in blood stage parasites found both were exported to the host erythrocyte cytoplasm (22). Additionally, PBANKA_1145400 could not be deleted and was predicted to be essential (22). Here we show that this PHIST protein partially localizes to the PV membrane in the late liver stage, confirming that it is indeed exported at this stage, and present at the host-parasite interface. Given the relatively low number of known PV proteins and the clear importance of this structure for host parasite interactions in the liver stage (11), we propose further characterization of this PHIST protein in the liver stage may be warranted.

In conclusion, this work provides a valuable resource for one of the least characterized stages of malaria parasites, the merosome. At the juncture between liver and blood stages, merosomes have a unique biology and initiate blood stage infection with relatively low numbers of parasites, both factors that could be exploited in the generation of new drugs and vaccines. This dataset and some of the insight gained from it should help inform future studies and interventions.

## Abbreviations

(PV): parasitophorous vacuole
(DMEM): Dulbecco’s Modified Eagle’s Medium
(FBS): fetal bovine serum
(RT): room temperature
(HBSS): Hank’s Buffered Saline Solution
(qPCR): quantitative PCR
(SCX): strong cation exchange
(HCD): higher-energy collisional dissociation
(NCE): normalized collision energy
(PSM): peptide spectrum match
(FDR): false discovery rate
(LFQ): label-free quantification
(GO): Gene Ontology
(RT): reverse transcriptase
(PEXEL): protein export element
(TPP): Trans-Proteomic Pipeline
(iBAQ): intensity-based absolute quantification
(EXP1): exported protein 1
(PTEX translocon): Plasmodium Translocon of Exported Proteins
(MTIP): myosin A tail domain interacting protein
(MSP): merozoite surface protein
(SERA): serine repeat antigen
(SUB): subtilisin-like protease
(ROM): rhomboid protease

## Acknowledgements

We thank the Insectary and Parasitology Core Facilities at the Johns Hopkins Malaria Research Institute and in particular Dr. Abhai Tripathi, Dr. Godfree Mlambo and Chris Kizito for their outstanding work. We are grateful to The Bloomberg Family Foundation for supporting these Core Facilities. We thank Dr. Sean Prigge for his assistance with the merosome genome quantitative PCR experiments, and Dr. Scott Lindner for sharing antibodies. We thank the Johns Hopkins School of Medicine Microscopy Facility for assistance with imaging experiments, and Dr. Hoku West-Foyle for his guidance with confocal microscopy and image analysis.

This work was supported by a Johns Hopkins Malaria Research Institute Pilot Grant to Drs. Photini Sinnis and Akhilesh Pandey, NIH Grant R01 AI056840 (P.S.), and a postdoctoral fellowship to Dr. Melanie Shears from the Johns Hopkins Provost’s Office. This work was also funded in part by the National Institutes of Health, grants R01GM087221 and 1S10RR027584 (R.M.), and K25AI119229 (K.S.). Funding for the Zeiss LSM 710 microscope was provided by NIH Grant S10 RR024550.

## Footnote

The content is solely the responsibility of the authors and does not necessarily represent the official views of the National Institutes of Health.

