## Supplementary Figures for "Proteomic Analysis of *Plasmodium* Merosomes: The Link Between Liver and Blood Stages in Malaria"

### Supplementary Figure 1

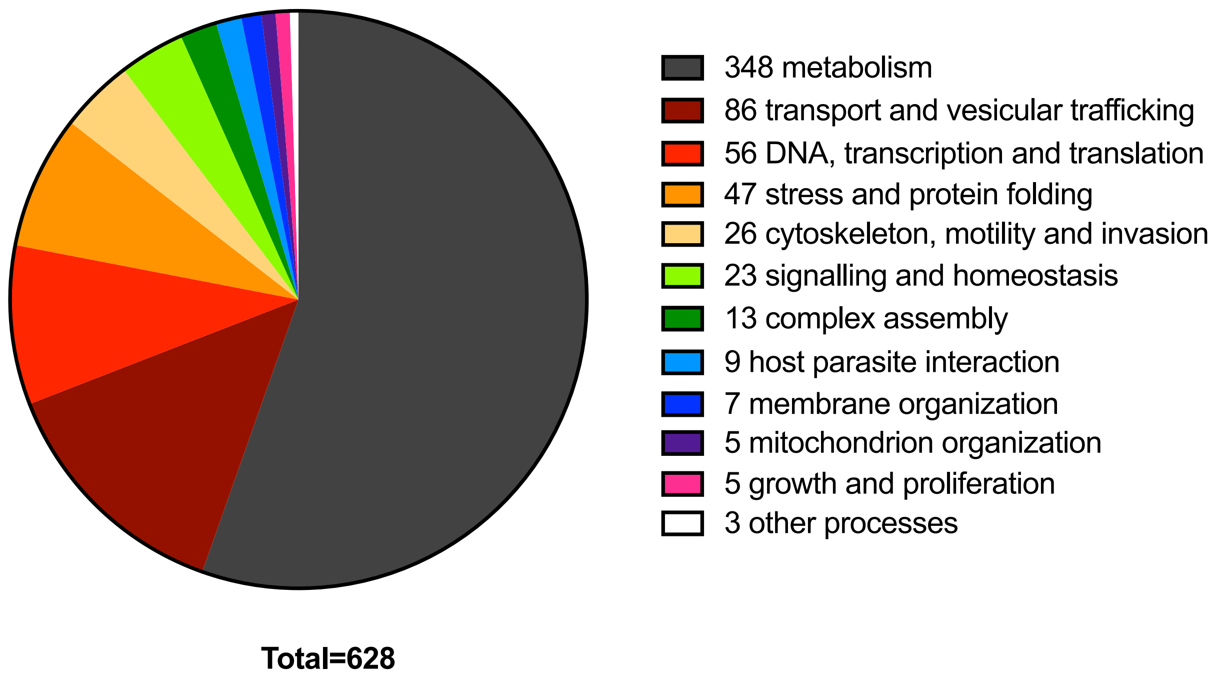


**Supplementary Figure 1. GO Biological Process terms for proteins in the core merosome proteome.** GO Biological Process terms were inferred from *P. falciparum* orthologs due to the relative paucity of annotations for *P. berghei*. Raw data for this graphic can be found in Supplementary Table 3. For this figure, GO terms were manually combined into groups of interest.

**Supplementary Figure 2. Annotated representative MS/MS spectra identifying putatively cleaved and acetylated PEXEL motifs.**

| Gene | Description | Peptide with cleaved & acetylated PEXEL |
|---|---|---|
| PBANKA_0208900 | StAR-related lipid transfer protein, putative | RIL.EEALMDVENVKENLK |

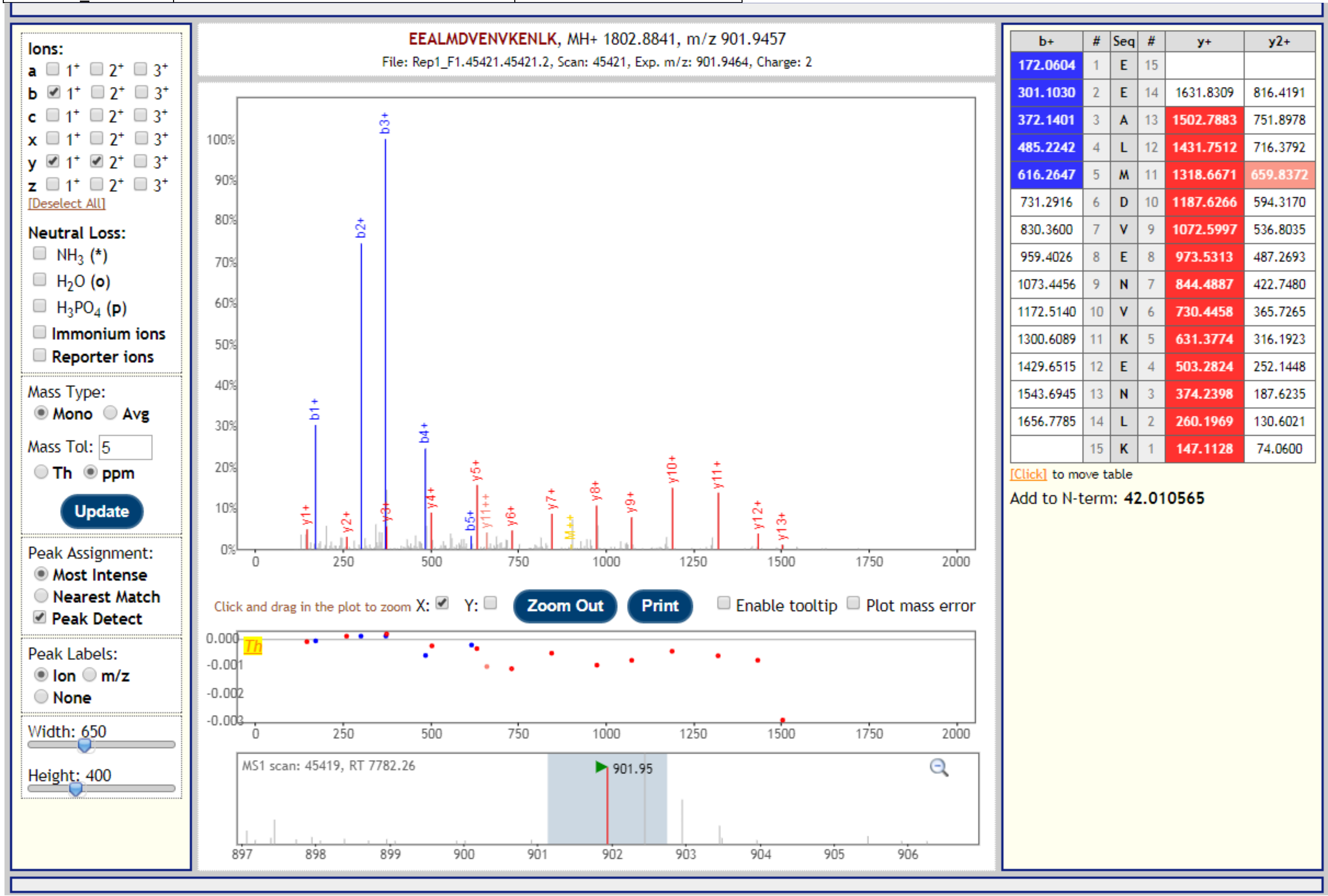

| Gene | Description | Peptide with cleaved & acetylated PEXEL |
|---|---|---|
| PBANKA_0208900 | StAR-related lipid transfer protein, putative | RIL.EEALMDVENVKENLK |

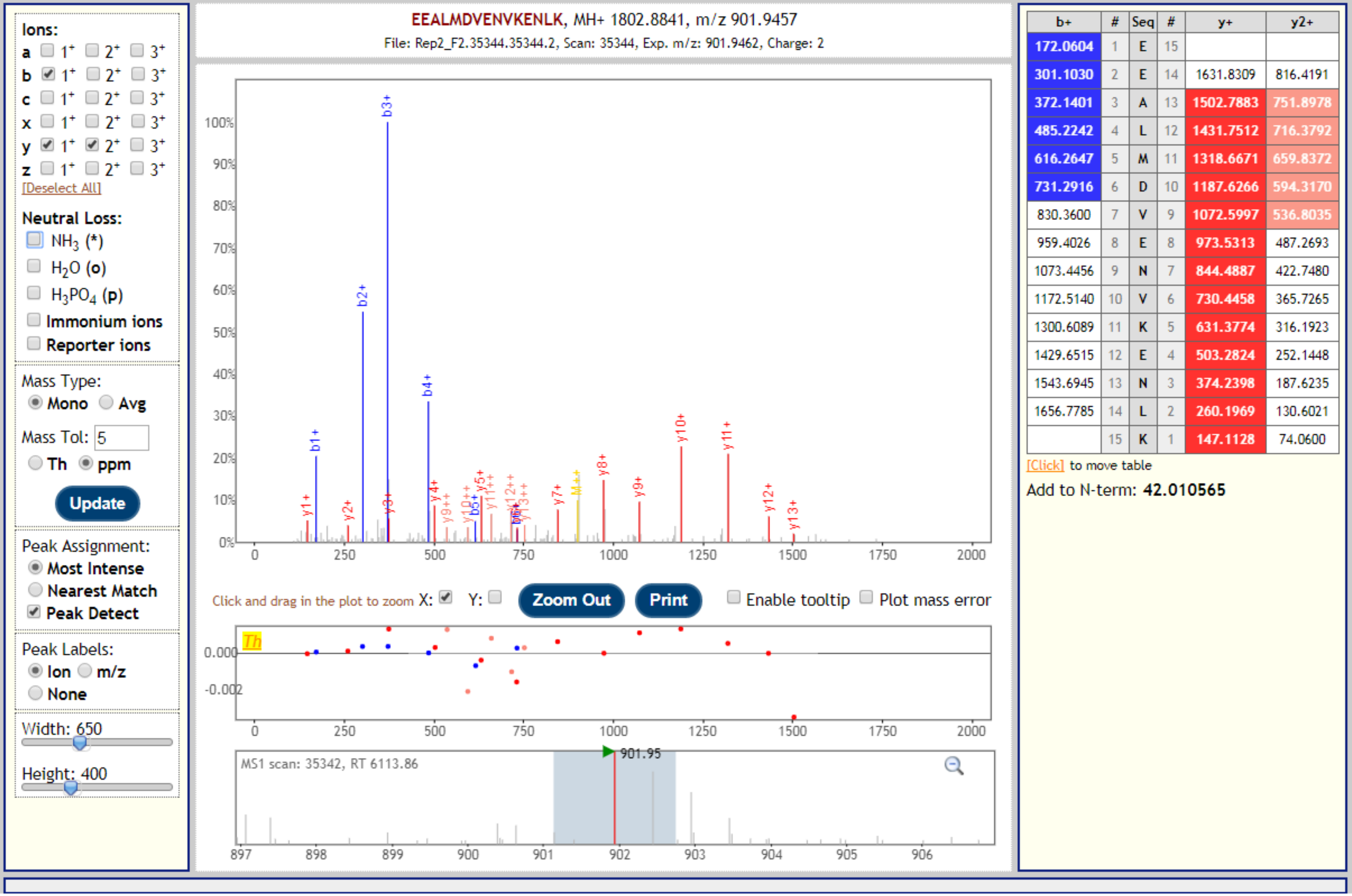


| b+ | # | Seq | # | y+ | y2+ |
|---|---|---|---|---|---|
| 172.0604 | 1 | E | 15 | | |
| 301.1030 | 2 | E | 14 | 1631.8309 | 816.4191 |
| 372.1401 | 3 | A | 13 | 1502.7883 | 751.8978 |
| 485.2242 | 4 | L | 12 | 1431.7512 | 716.3792 |
| 616.2647 | 5 | M | 11 | 1318.6671 | 659.8372 |
| 731.2916 | 6 | D | 10 | 1187.6266 | 594.3170 |
| 830.3600 | 7 | V | 9 | 1072.5997 | 536.8035 |
| 959.4026 | 8 | E | 8 | 973.5313 | 487.2693 |
| 1073.4456 | 9 | N | 7 | 844.4887 | 422.7480 |
| 1172.5140 | 10 | V | 6 | 730.4458 | 365.7265 |
| 1300.6089 | 11 | K | 5 | 631.3774 | 316.1923 |
| 1429.6515 | 12 | E | 4 | 503.2824 | 252.1448 |
| 1543.6945 | 13 | N | 3 | 374.2398 | 187.6235 |
| 1656.7785 | 14 | L | 2 | 260.1969 | 130.6021 |
| | 15 | K | 1 | 147.1128 | 74.0600 |

| Gene | Description | Peptide with cleaved & acetylated PEXEL |
|---|---|---|
| PBANKA_0208900 | StAR-related lipid transfer protein, putative | RIL.EEALMDVENVKENLK |

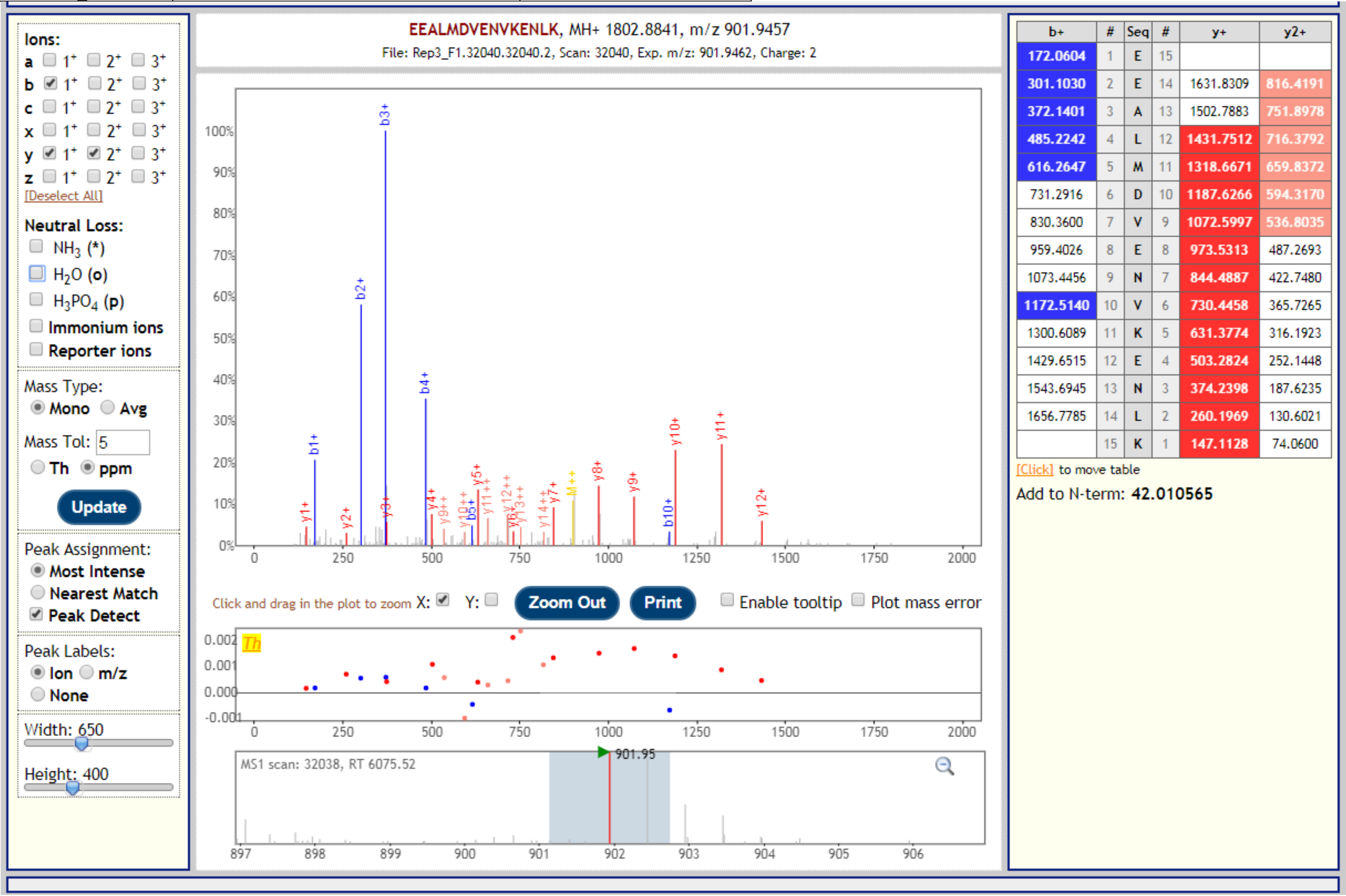

| Gene | Description | Peptide with cleaved & acetylated PEXEL |
|---|---|---|
| PBANKA_0208900 | StAR-related lipid transfer protein, putative | RIL.EEALMDVENVKENLKYVQQAK |

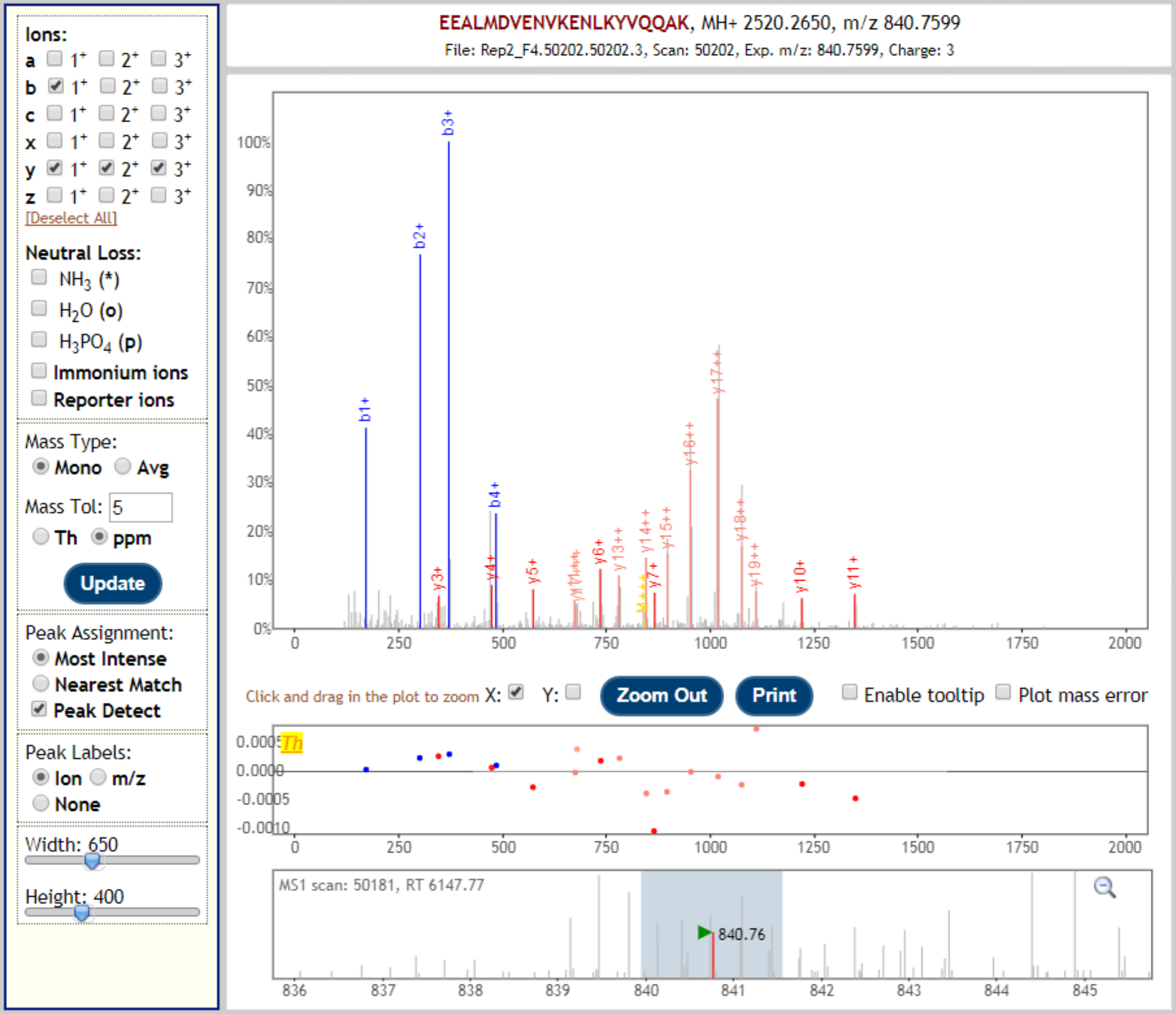


| b+ | # | Seq | # | y+ | y2+ | y3+ |
|---|---|---|---|---|---|---|
| 172.0604 | 1 | E | 21 | | | |
| 301.1030 | 2 | E | 20 | 2349.2119 | 1175.1096 | 783.7421 |
| 372.1401 | 3 | A | 19 | 2220.1693 | 1110.5883 | 740.7279 |
| 485.2242 | 4 | L | 18 | 2149.1322 | 1075.0697 | 717.0489 |
| 616.2647 | 5 | M | 17 | 2036.0481 | 1018.5277 | 679.3542 |
| 731.2916 | 6 | D | 16 | 1905.0076 | 953.0074 | 635.6741 |
| 830.3600 | 7 | V | 15 | 1789.9807 | 895.4940 | 597.3317 |
| 959.4026 | 8 | E | 14 | 1690.9123 | 845.9598 | 564.3089 |
| 1073.4456 | 9 | N | 13 | 1561.8697 | 781.4385 | 521.2947 |
| 1172.5140 | 10 | V | 12 | 1447.8267 | 724.4170 | 483.2804 |
| 1300.6089 | 11 | K | 11 | 1348.7583 | 674.8828 | 450.2576 |
| 1429.6515 | 12 | E | 10 | 1220.6634 | 610.8353 | 407.5593 |
| 1543.6945 | 13 | N | 9 | 1091.6208 | 546.3140 | 364.5451 |
| 1656.7785 | 14 | L | 8 | 977.5778 | 489.2926 | 326.5308 |
| 1784.8735 | 15 | K | 7 | 864.4938 | 432.7505 | 288.8361 |
| 1947.9368 | 16 | Y | 6 | 736.3988 | 368.7030 | 246.1378 |
| 2047.0052 | 17 | V | 5 | 573.3355 | 287.1714 | 191.7833 |
| 2175.0638 | 18 | Q | 4 | 474.2671 | 237.6372 | 158.7605 |
| 2303.1224 | 19 | Q | 3 | 346.2085 | 173.6079 | 116.0743 |
| 2374.1595 | 20 | A | 2 | 218.1499 | 109.5786 | 73.3882 |
| | 21 | K | 1 | 147.1128 | 74.0600 | 49.7091 |

[Click] to move table

Add to N-term: **42.010565**

| Gene | Description | Peptide with cleaved & acetylated PEXEL |
|---|---|---|
| PBANKA_0208900 | StAR-related lipid transfer protein, putative | RIL.EEALMDVENVKENLKYVQQAK |

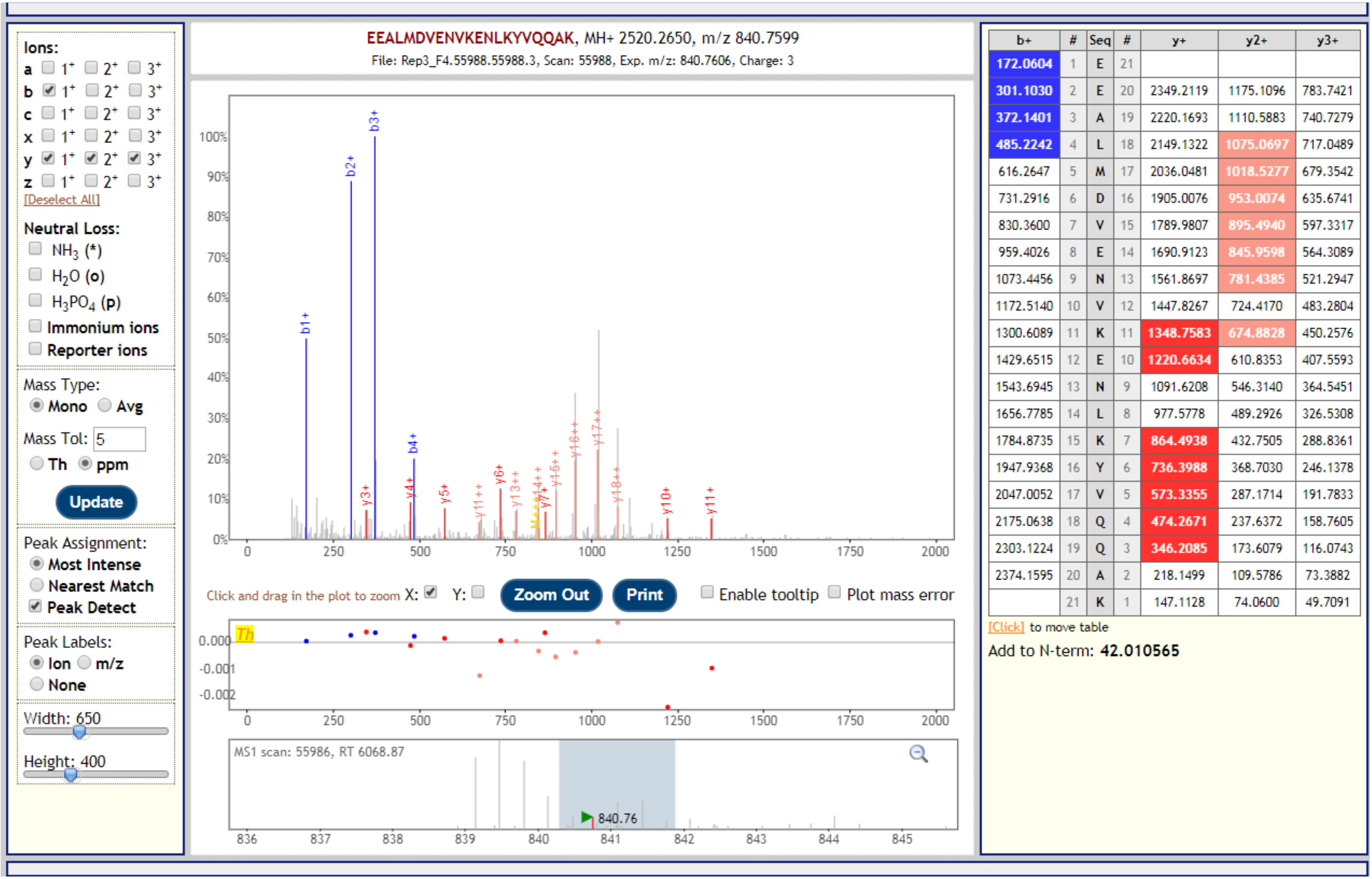

| Gene | Description | Peptide with cleaved & acetylated PEXEL |
|---|---|---|
| PBANKA_0208900 | StAR-related lipid transfer protein, putative | RIL.EEALMDVENVK |

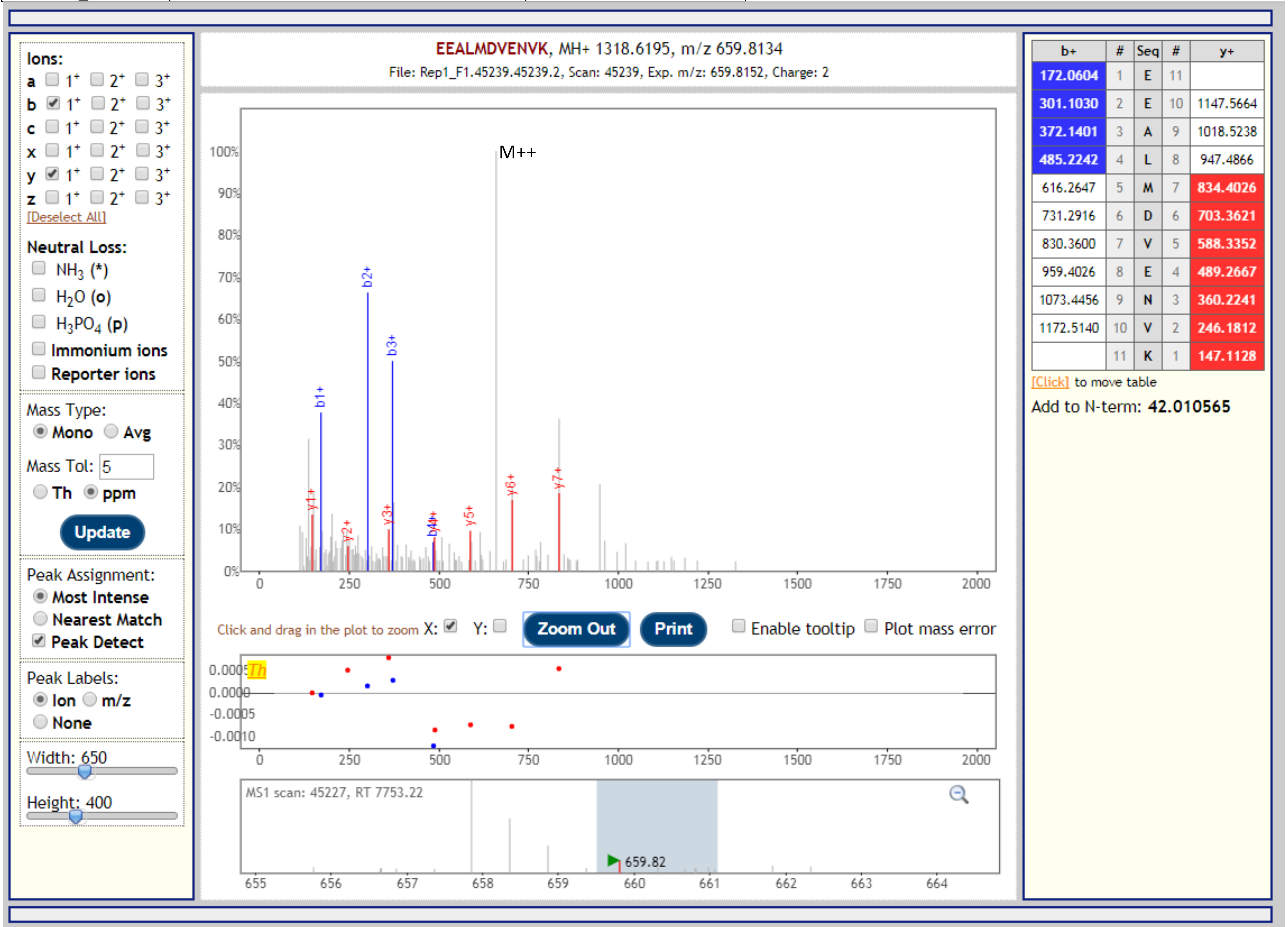

| Gene | Description | Peptide with cleaved & acetylated PEXEL |
|---|---|---|
| PBANKA_1400700 | Plasmodium exported protein, unknown function | RHL.AEYIPNISNPTGNVYHFSR |

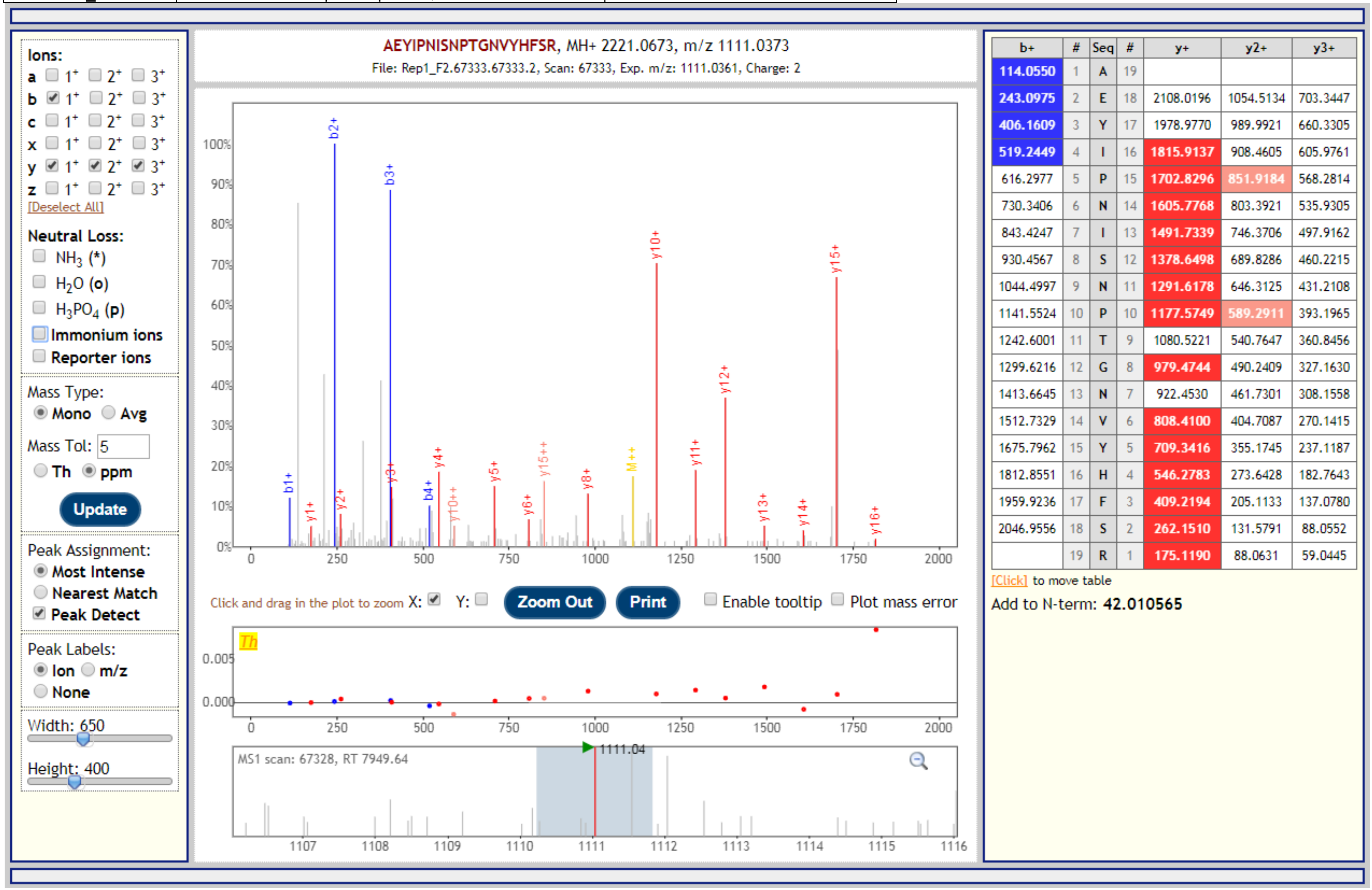


| b+ | # | Seq | # | y+ | y2+ | y3+ |
|---|---|---|---|---|---|---|
| 114.0550 | 1 | A | 19 | | | |
| 243.0975 | 2 | E | 18 | 2108.0196 | 1054.5134 | 703.3447 |
| 406.1609 | 3 | Y | 17 | 1978.9770 | 989.9921 | 660.3305 |
| 519.2449 | 4 | I | 16 | 1815.9137 | 908.4605 | 605.9761 |
| 616.2977 | 5 | P | 15 | 1702.8296 | 851.9184 | 568.2814 |
| 730.3406 | 6 | N | 14 | 1605.7768 | 803.3921 | 535.9305 |
| 843.4247 | 7 | I | 13 | 1491.7339 | 746.3706 | 497.9162 |
| 930.4567 | 8 | S | 12 | 1378.6498 | 689.8286 | 460.2215 |
| 1044.4997 | 9 | N | 11 | 1291.6178 | 646.3125 | 431.2108 |
| 1141.5524 | 10 | P | 10 | 1177.5749 | 589.2911 | 393.1965 |
| 1242.6001 | 11 | T | 9 | 1080.5221 | 540.7647 | 360.8456 |
| 1299.6216 | 12 | G | 8 | 979.4744 | 490.2409 | 327.1630 |
| 1413.6645 | 13 | N | 7 | 922.4530 | 461.7301 | 308.1558 |
| 1512.7329 | 14 | V | 6 | 808.4100 | 404.7087 | 270.1415 |
| 1675.7962 | 15 | Y | 5 | 709.3416 | 355.1745 | 237.1187 |
| 1812.8551 | 16 | H | 4 | 546.2783 | 273.6428 | 182.7643 |
| 1959.9236 | 17 | F | 3 | 409.2194 | 205.1133 | 137.0780 |
| 2046.9556 | 18 | S | 2 | 262.1510 | 131.5791 | 88.0552 |
| | 19 | R | 1 | 175.1190 | 88.0631 | 59.0445 |

| Gene | Description | Peptide with cleaved & acetylated PEXEL |
|---|---|---|
| PBANKA_1400700 | Plasmodium exported protein, unknown function | RHL.AEYIPNISNPTGNVYHFSR |

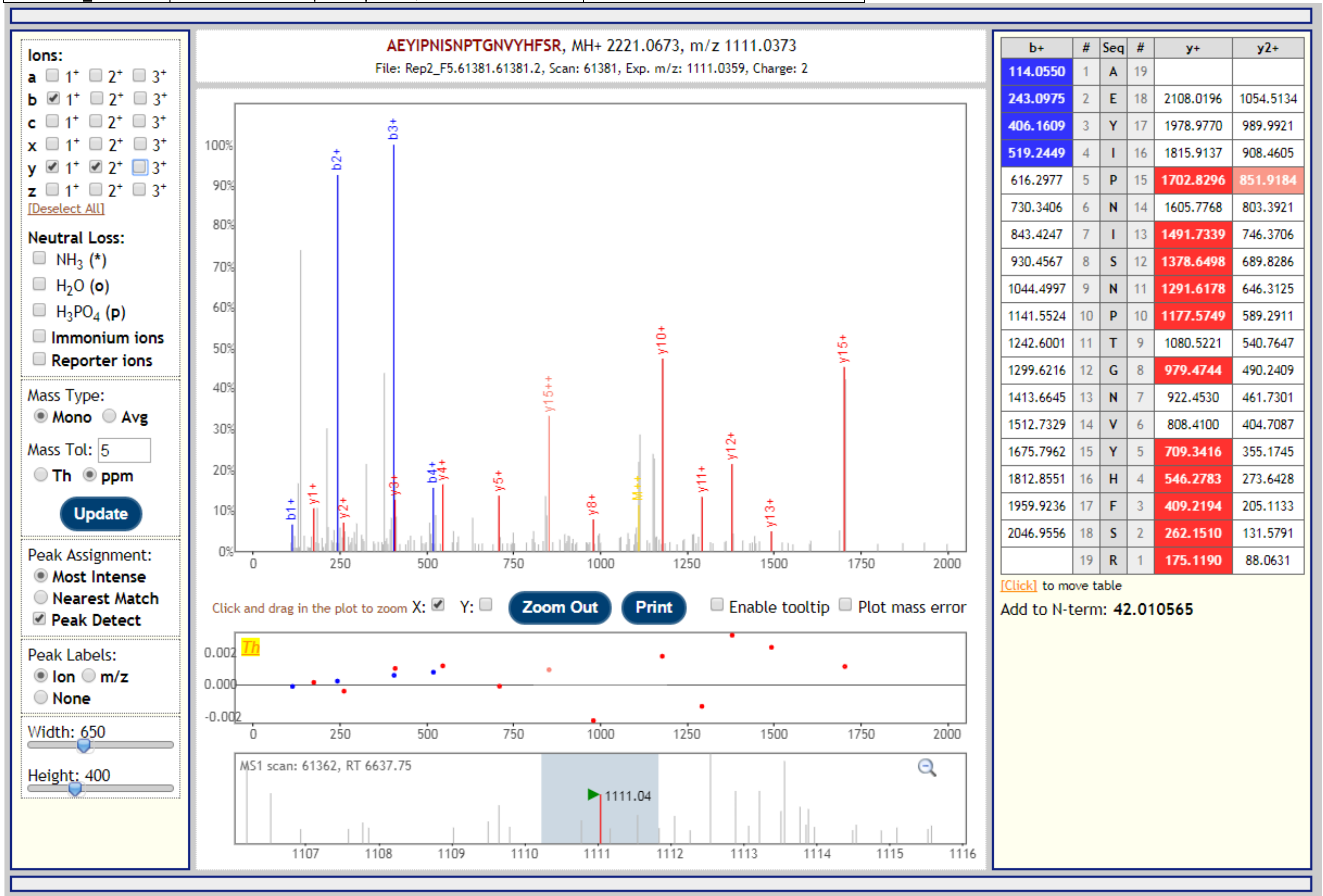


| b+ | # | Seq | # | y+ | y2+ |
|---|---|---|---|---|---|
| 114.0550 | 1 | A | 19 | | |
| 243.0975 | 2 | E | 18 | 2108.0196 | 1054.5134 |
| 406.1609 | 3 | Y | 17 | 1978.9770 | 989.9921 |
| 519.2449 | 4 | I | 16 | 1815.9137 | 908.4605 |
| 616.2977 | 5 | P | 15 | 1702.8296 | 851.9184 |
| 730.3406 | 6 | N | 14 | 1605.7768 | 803.3921 |
| 843.4247 | 7 | I | 13 | 1491.7339 | 746.3706 |
| 930.4567 | 8 | S | 12 | 1378.6498 | 689.8286 |
| 1044.4997 | 9 | N | 11 | 1291.6178 | 646.3125 |
| 1141.5524 | 10 | P | 10 | 1177.5749 | 589.2911 |
| 1242.6001 | 11 | T | 9 | 1080.5221 | 540.7647 |
| 1299.6216 | 12 | G | 8 | 979.4744 | 490.2409 |
| 1413.6645 | 13 | N | 7 | 922.4530 | 461.7301 |
| 1512.7329 | 14 | V | 6 | 808.4100 | 404.7087 |
| 1675.7962 | 15 | Y | 5 | 709.3416 | 355.1745 |
| 1812.8551 | 16 | H | 4 | 546.2783 | 273.6428 |
| 1959.9236 | 17 | F | 3 | 409.2194 | 205.1133 |
| 2046.9556 | 18 | S | 2 | 262.1510 | 131.5791 |
| | 19 | R | 1 | 175.1190 | 88.0631 |

| Gene | Description | Peptide with cleaved & acetylated PEXEL |
|---|---|---|
| PBANKA_1400700 | Plasmodium exported protein, unknown function | RHL.AEYIPNISNPTGNVYHFSR |

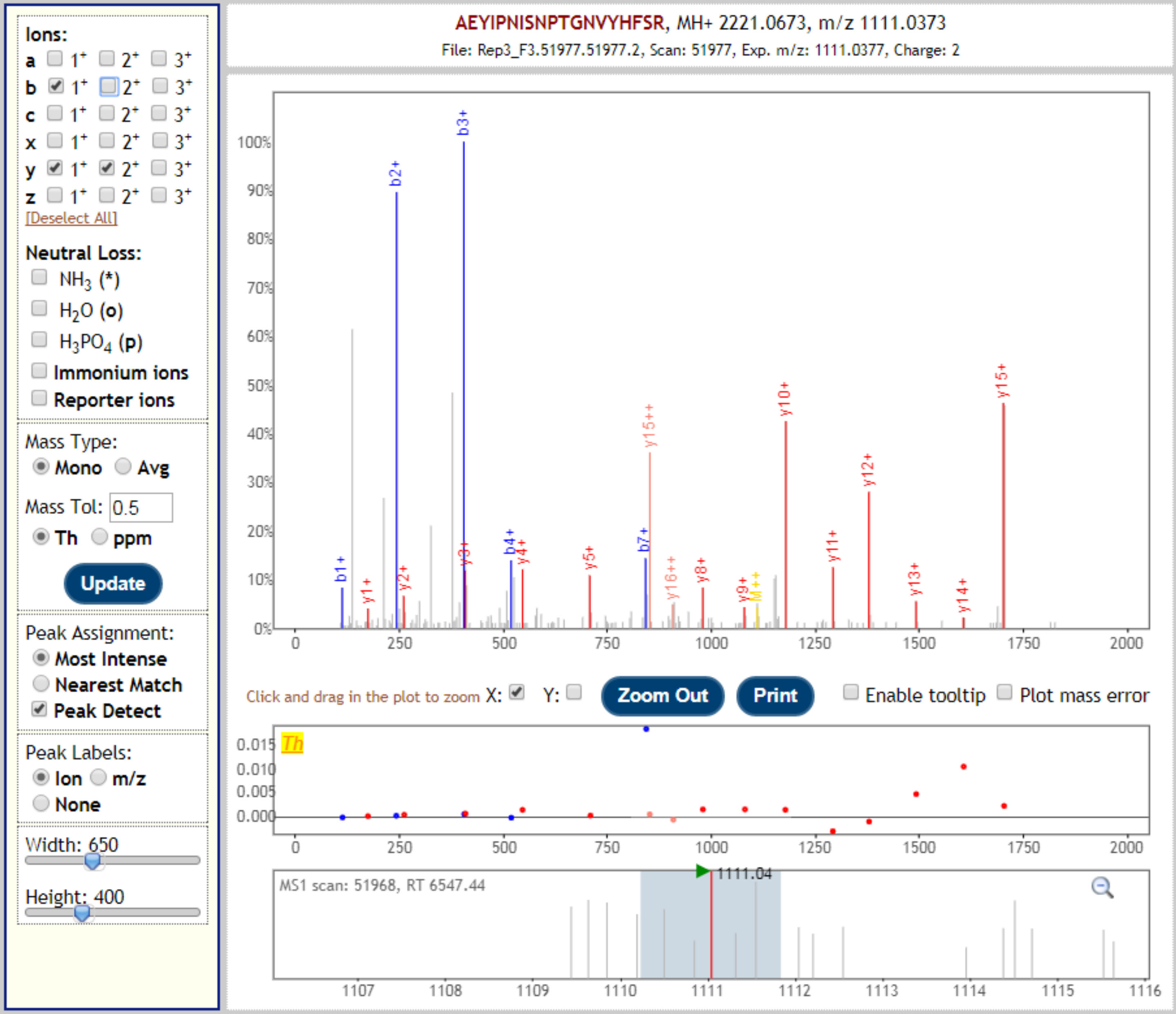


| b+ | # | Seq | # | y+ | y2+ |
|---|---|---|---|---|---|
| 114.0550 | 1 | A | 19 | | |
| 243.0975 | 2 | E | 18 | 2108.0196 | 1054.5134 |
| 406.1609 | 3 | Y | 17 | 1978.9770 | 989.9921 |
| 519.2449 | 4 | I | 16 | 1815.9137 | 908.4605 |
| 616.2977 | 5 | P | 15 | 1702.8296 | 851.9184 |
| 730.3406 | 6 | N | 14 | 1605.7768 | 803.3921 |
| 843.4247 | 7 | I | 13 | 1491.7339 | 746.3706 |
| 930.4567 | 8 | S | 12 | 1378.6498 | 689.8286 |
| 1044.4997 | 9 | N | 11 | 1291.6178 | 646.3125 |
| 1141.5524 | 10 | P | 10 | 1177.5749 | 589.2911 |
| 1242.6001 | 11 | T | 9 | 1080.5221 | 540.7647 |
| 1299.6216 | 12 | G | 8 | 979.4744 | 490.2409 |
| 1413.6645 | 13 | N | 7 | 922.4530 | 461.7301 |
| 1512.7329 | 14 | V | 6 | 808.4100 | 404.7087 |
| 1675.7962 | 15 | Y | 5 | 709.3416 | 355.1745 |
| 1812.8551 | 16 | H | 4 | 546.2783 | 273.6428 |
| 1959.9236 | 17 | F | 3 | 409.2194 | 205.1133 |
| 2046.9556 | 18 | S | 2 | 262.1510 | 131.5791 |
| | 19 | R | 1 | 175.1190 | 88.0631 |

[Click] to move table

Add to N-term: **42.010565**

| Gene | Description | Peptide with cleaved & acetylated PEXEL |
|---|---|---|
| PBANKA_1003000 | liver specific protein 2 | RIL.AETQHENLVLSTDKDKER |

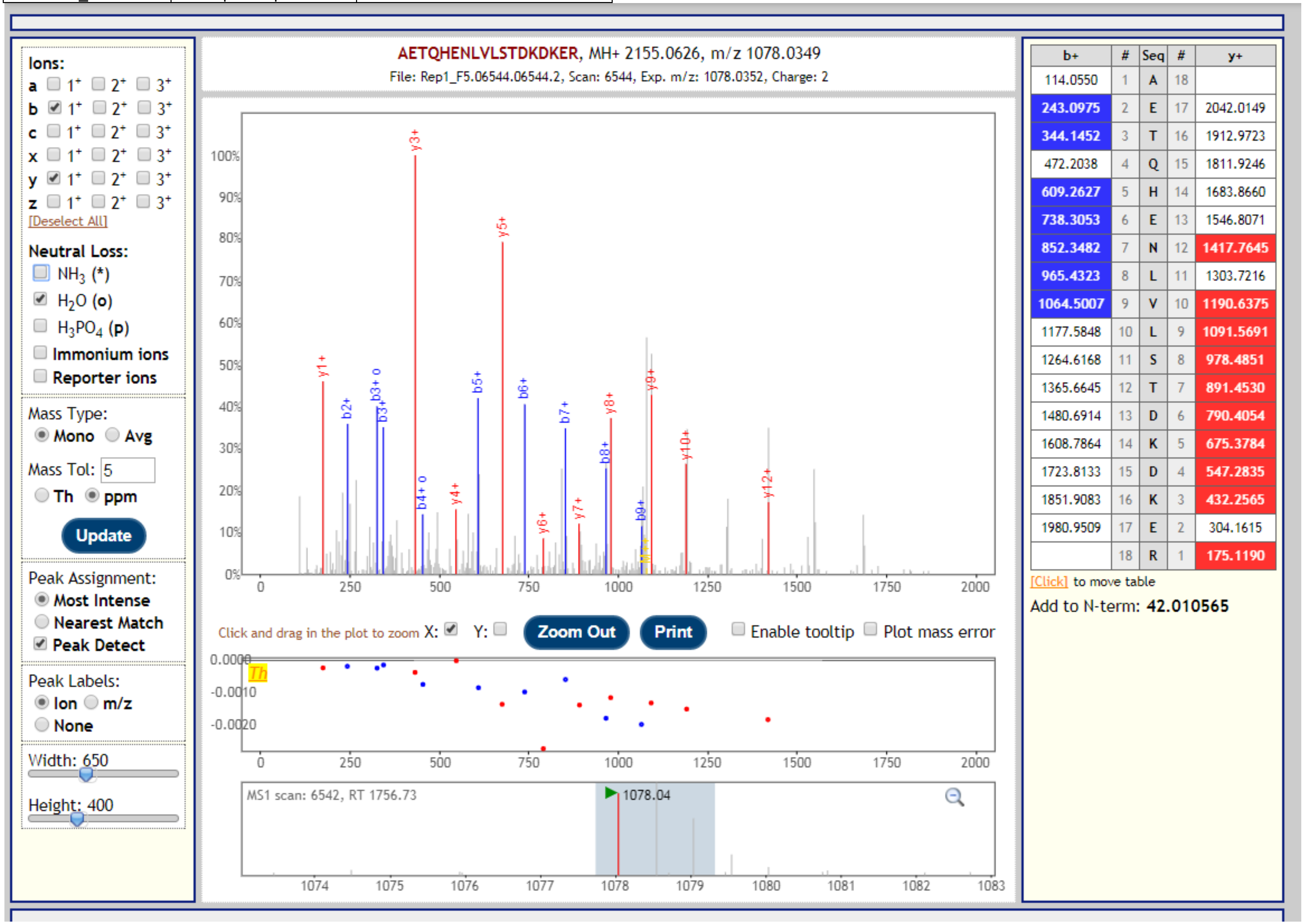

| Gene | Description | Peptide with cleaved & acetylated PEXEL |
|---|---|---|
| PBANKA_1003000 | liver specific protein 2 | RIL.AETQHENLVLSTDKDKER |

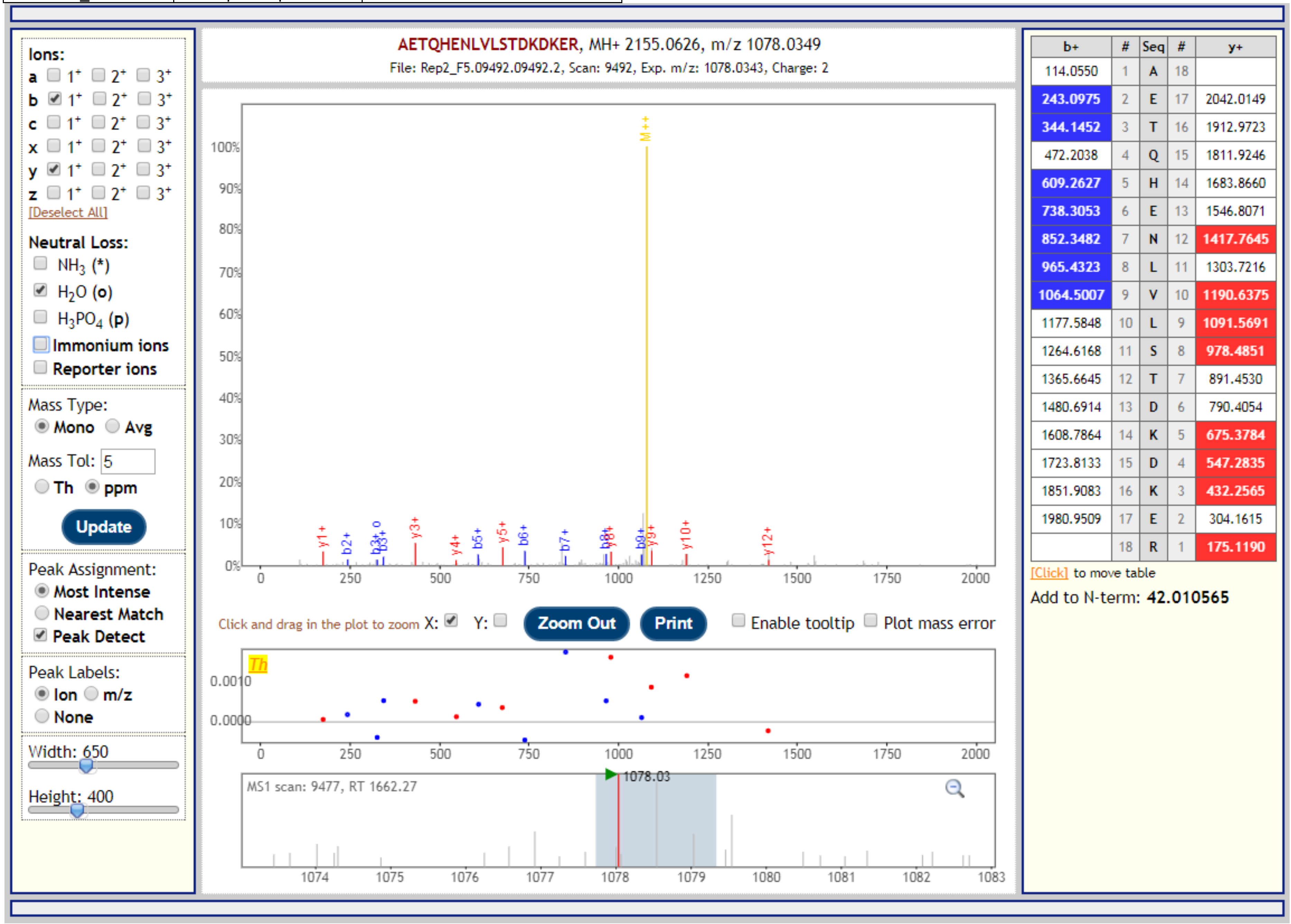


| b+ | # | Seq | # | y+ |
|---|---|---|---|---|
| 114.0550 | 1 | A | 18 | |
| 243.0975 | 2 | E | 17 | 2042.0149 |
| 344.1452 | 3 | T | 16 | 1912.9723 |
| 472.2038 | 4 | Q | 15 | 1811.9246 |
| 609.2627 | 5 | H | 14 | 1683.8660 |
| 738.3053 | 6 | E | 13 | 1546.8071 |
| 852.3482 | 7 | N | 12 | 1417.7645 |
| 965.4323 | 8 | L | 11 | 1303.7216 |
| 1064.5007 | 9 | V | 10 | 1190.6375 |
| 1177.5848 | 10 | L | 9 | 1091.5691 |
| 1264.6168 | 11 | S | 8 | 978.4851 |
| 1365.6645 | 12 | T | 7 | 891.4530 |
| 1480.6914 | 13 | D | 6 | 790.4054 |
| 1608.7864 | 14 | K | 5 | 675.3784 |
| 1723.8133 | 15 | D | 4 | 547.2835 |
| 1851.9083 | 16 | K | 3 | 432.2565 |
| 1980.9509 | 17 | E | 2 | 304.1615 |
| | 18 | R | 1 | 175.1190 |

| Gene | Description | Peptide with cleaved & acetylated PEXEL |
|---|---|---|
| PBANKA_1003000 | liver specific protein 2 | RIL.AETQHENLVLSTDKDKER |

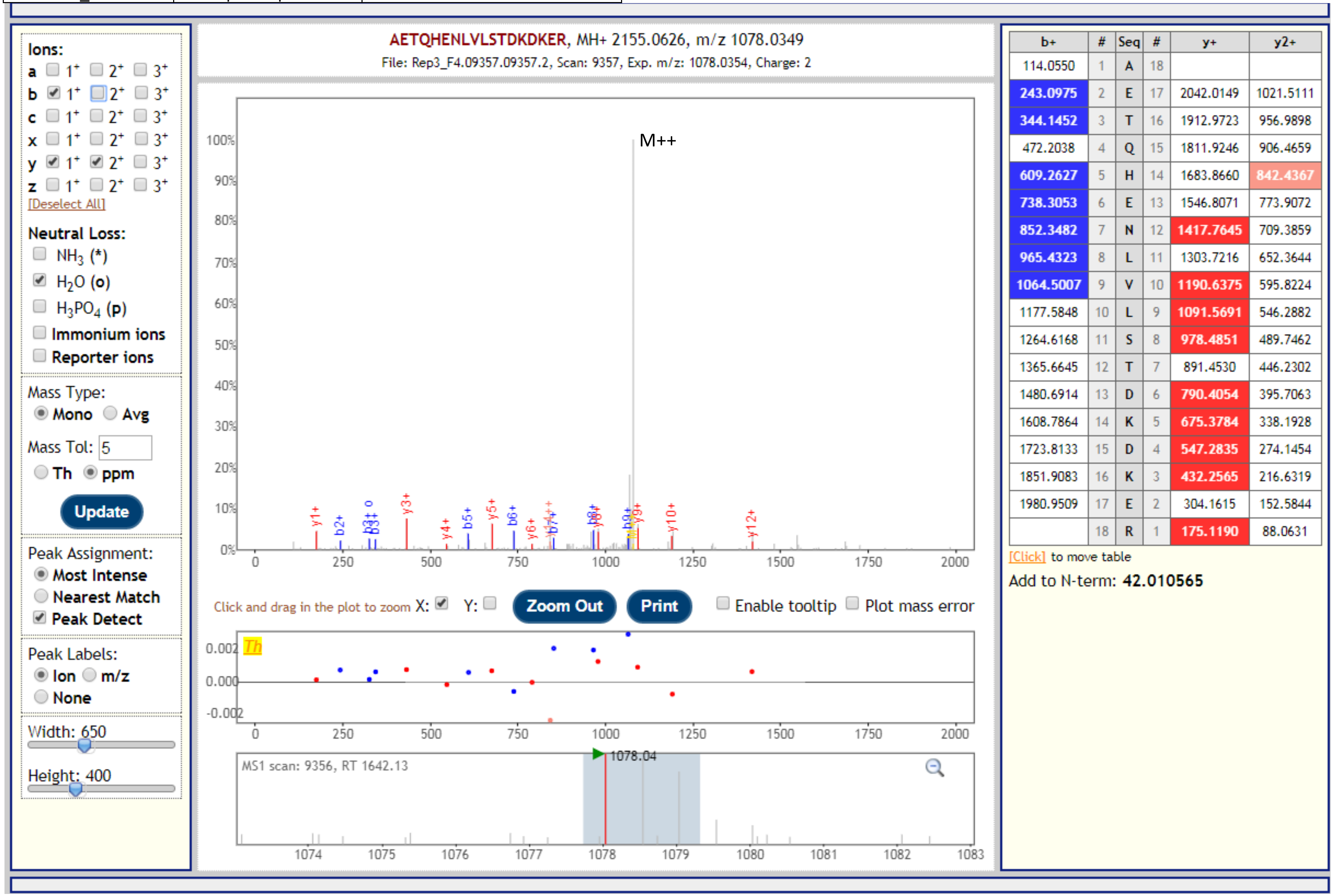


| b+ | # | Seq | # | y+ | y2+ |
|---|---|---|---|---|---|
| 114.0550 | 1 | A | 18 | | |
| 243.0975 | 2 | E | 17 | 2042.0149 | 1021.5111 |
| 344.1452 | 3 | T | 16 | 1912.9723 | 956.9898 |
| 472.2038 | 4 | Q | 15 | 1811.9246 | 906.4659 |
| 609.2627 | 5 | H | 14 | 1683.8660 | 842.4367 |
| 738.3053 | 6 | E | 13 | 1546.8071 | 773.9072 |
| 852.3482 | 7 | N | 12 | 1417.7645 | 709.3859 |
| 965.4323 | 8 | L | 11 | 1303.7216 | 652.3644 |
| 1064.5007 | 9 | V | 10 | 1190.6375 | 595.8224 |
| 1177.5848 | 10 | L | 9 | 1091.5691 | 546.2882 |
| 1264.6168 | 11 | S | 8 | 978.4851 | 489.7462 |
| 1365.6645 | 12 | T | 7 | 891.4530 | 446.2302 |
| 1480.6914 | 13 | D | 6 | 790.4054 | 395.7063 |
| 1608.7864 | 14 | K | 5 | 675.3784 | 338.1928 |
| 1723.8133 | 15 | D | 4 | 547.2835 | 274.1454 |
| 1851.9083 | 16 | K | 3 | 432.2565 | 216.6319 |
| 1980.9509 | 17 | E | 2 | 304.1615 | 152.5844 |
| | 18 | R | 1 | 175.1190 | 88.0631 |

| Gene | Description | Peptide with cleaved & acetylated PEXEL |
|---|---|---|
| PBANKA_1003000 | liver specific protein 2 | RLI.AEKESENNSENNSQDVK |

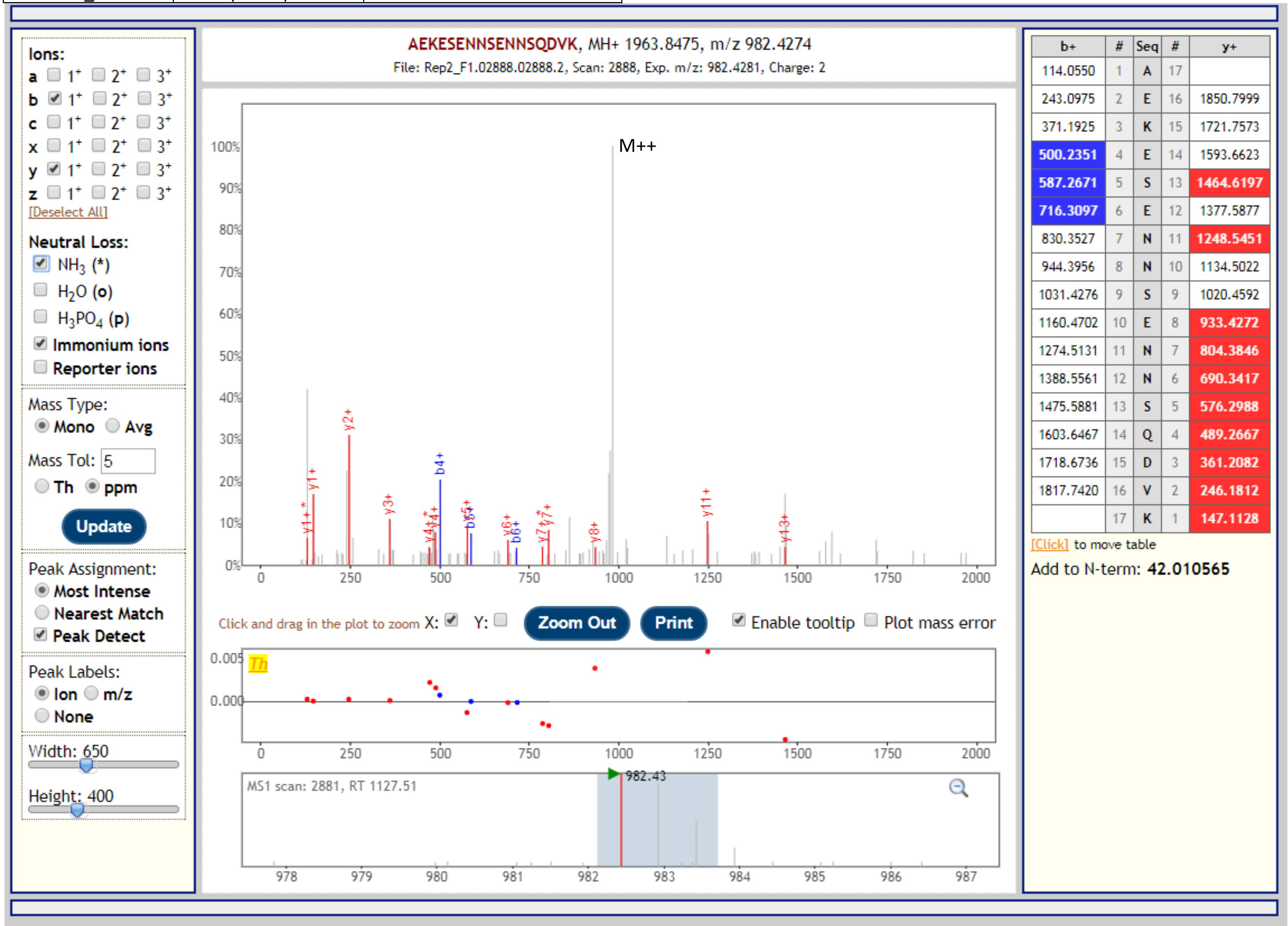

| Gene | Description | Peptide with cleaved & acetylated PEXEL |
|---|---|---|
| PBANKA_1003000 | liver specific protein 2 | RLI.AEKESENNSENNSQDVK |

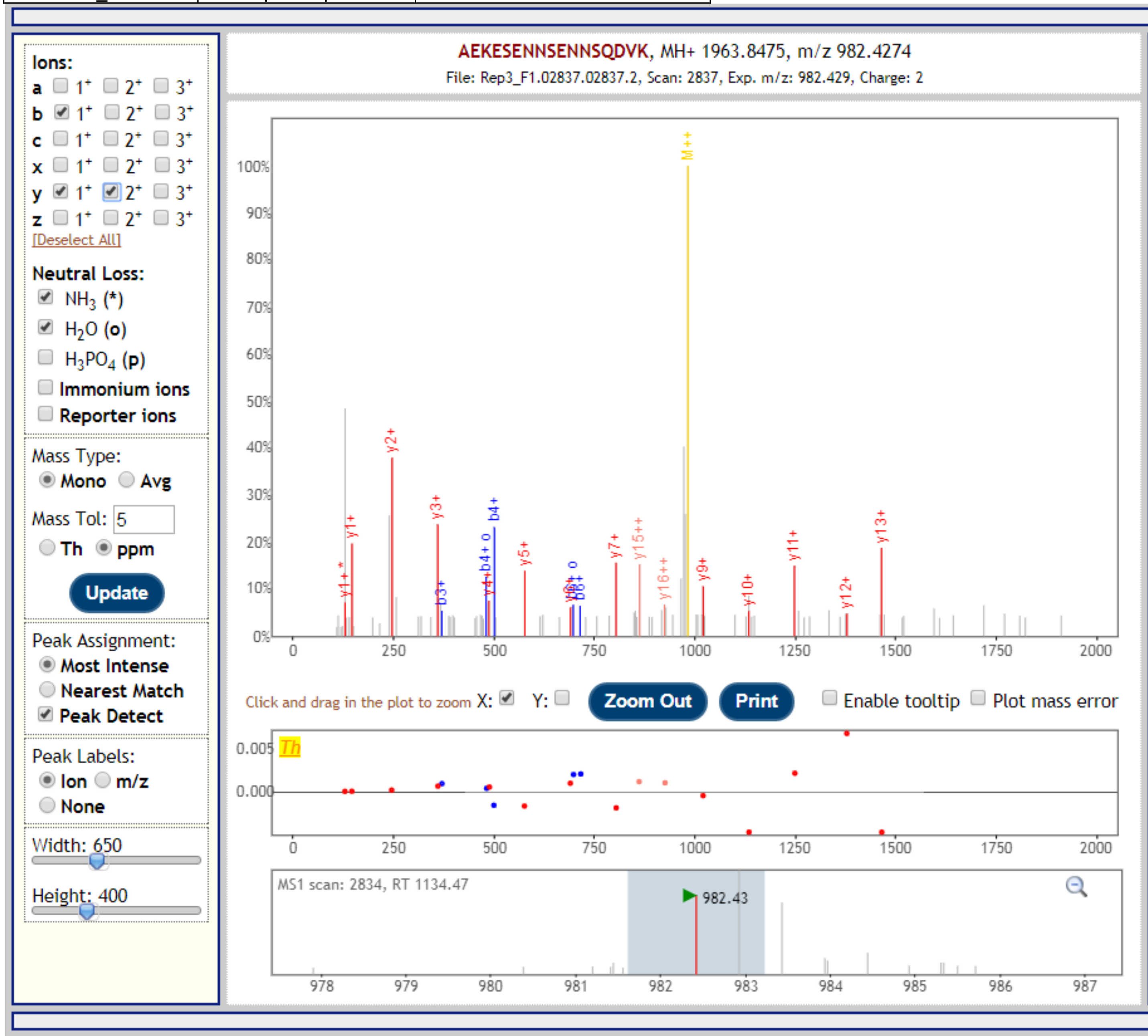


| b+ | # | Seq | # | y+ | y2+ |
|---|---|---|---|---|---|
| 114.0550 | 1 | A | 17 | | |
| 243.0975 | 2 | E | 16 | 1850.7999 | 925.9036 |
| 371.1925 | 3 | K | 15 | 1721.7573 | 861.3823 |
| 500.2351 | 4 | E | 14 | 1593.6623 | 797.3348 |
| 587.2671 | 5 | S | 13 | 1464.6197 | 732.8135 |
| 716.3097 | 6 | E | 12 | 1377.5877 | 689.2975 |
| 830.3527 | 7 | N | 11 | 1248.5451 | 624.7762 |
| 944.3956 | 8 | N | 10 | 1134.5022 | 567.7547 |
| 1031.4276 | 9 | S | 9 | 1020.4592 | 510.7333 |
| 1160.4702 | 10 | E | 8 | 933.4272 | 467.2172 |
| 1274.5131 | 11 | N | 7 | 804.3846 | 402.6959 |
| 1388.5561 | 12 | N | 6 | 690.3417 | 345.6745 |
| 1475.5881 | 13 | S | 5 | 576.2988 | 288.6530 |
| 1603.6467 | 14 | Q | 4 | 489.2667 | 245.1370 |
| 1718.6736 | 15 | D | 3 | 361.2082 | 181.1077 |
| 1817.7420 | 16 | V | 2 | 246.1812 | 123.5942 |
| | 17 | K | 1 | 147.1128 | 74.0600 |

[Click] to move table

Add to N-term: **42.010565**

| Gene | Description | Peptide with cleaved & acetylated PEXEL |
|---|---|---|
| PBANKA_1003000 | liver specific protein 2 | RLI.AEKESENNSENNSQDVKQDK |

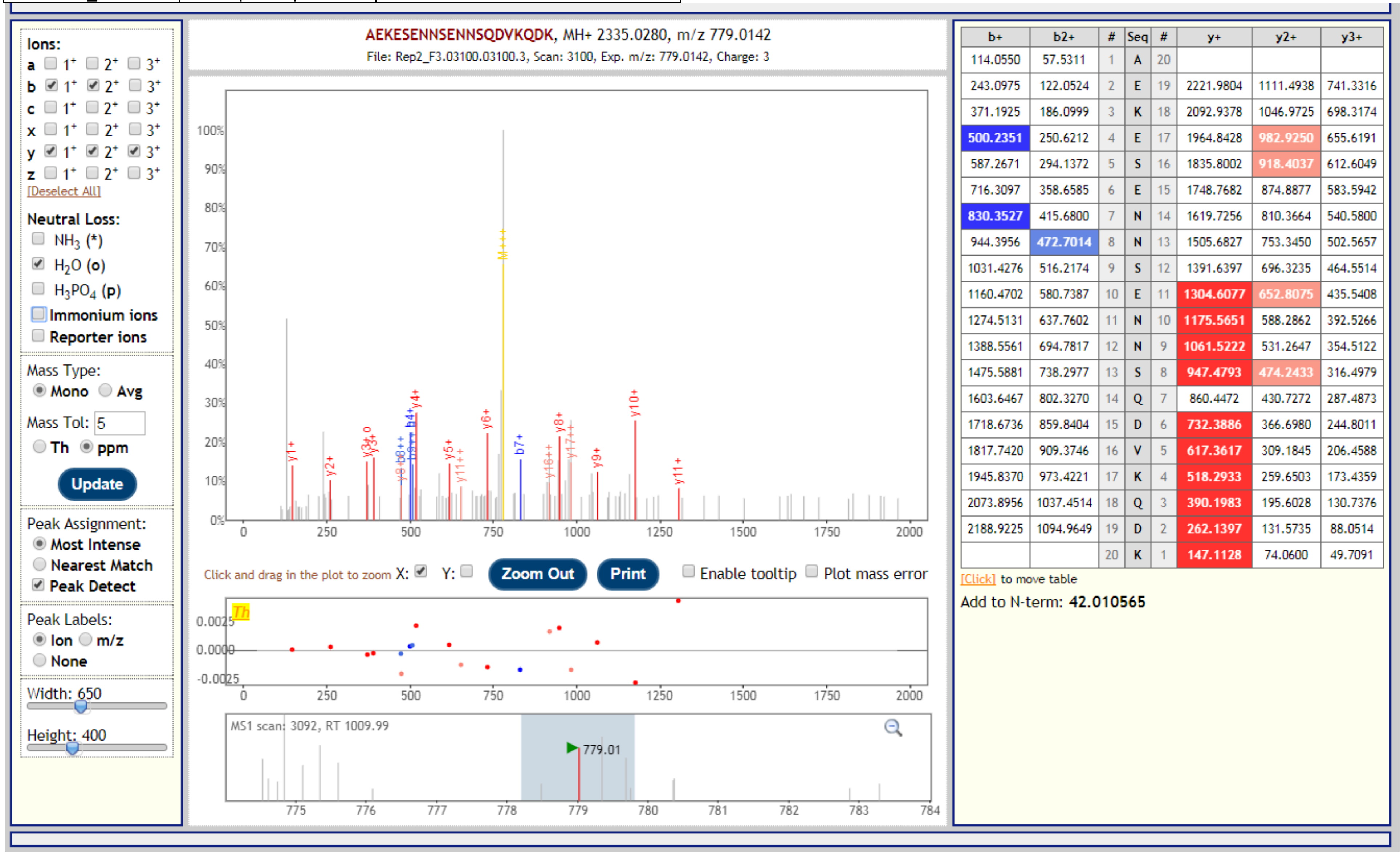


| b+ | b2+ | # | Seq | # | y+ | y2+ | y3+ |
|---|---|---|---|---|---|---|---|
| 114.0550 | 57.5311 | 1 | A | 20 | | | |
| 243.0975 | 122.0524 | 2 | E | 19 | 2221.9804 | 1111.4938 | 741.3316 |
| 371.1925 | 186.0999 | 3 | K | 18 | 2092.9378 | 1046.9725 | 698.3174 |
| 500.2351 | 250.6212 | 4 | E | 17 | 1964.8428 | 982.9250 | 655.6191 |
| 587.2671 | 294.1372 | 5 | S | 16 | 1835.8002 | 918.4037 | 612.6049 |
| 716.3097 | 358.6585 | 6 | E | 15 | 1748.7682 | 874.8877 | 583.5942 |
| 830.3527 | 415.6800 | 7 | N | 14 | 1619.7256 | 810.3664 | 540.5800 |
| 944.3956 | 472.7014 | 8 | N | 13 | 1505.6827 | 753.3450 | 502.5657 |
| 1031.4276 | 516.2174 | 9 | S | 12 | 1391.6397 | 696.3235 | 464.5514 |
| 1160.4702 | 580.7387 | 10 | E | 11 | 1304.6077 | 652.8075 | 435.5408 |
| 1274.5131 | 637.7602 | 11 | N | 10 | 1175.5651 | 588.2862 | 392.5266 |
| 1388.5561 | 694.7817 | 12 | N | 9 | 1061.5222 | 531.2647 | 354.5122 |
| 1475.5881 | 738.2977 | 13 | S | 8 | 947.4793 | 474.2433 | 316.4979 |
| 1603.6467 | 802.3270 | 14 | Q | 7 | 860.4472 | 430.7272 | 287.4873 |
| 1718.6736 | 859.8404 | 15 | D | 6 | 732.3886 | 366.6980 | 244.8011 |
| 1817.7420 | 909.3746 | 16 | V | 5 | 617.3617 | 309.1845 | 206.4588 |
| 1945.8370 | 973.4221 | 17 | K | 4 | 518.2933 | 259.6503 | 173.4359 |
| 2073.8956 | 1037.4514 | 18 | Q | 3 | 390.1983 | 195.6028 | 130.7376 |
| 2188.9225 | 1094.9649 | 19 | D | 2 | 262.1397 | 131.5735 | 88.0514 |
| | | 20 | K | 1 | 147.1128 | 74.0600 | 49.7091 |

| Gene | Description | Peptide with cleaved & acetylated PEXEL |
|---|---|---|
| PBANKA_1003000 | liver specific protein 2 | RLI.AEKESENNSENNSQDVKQDK |

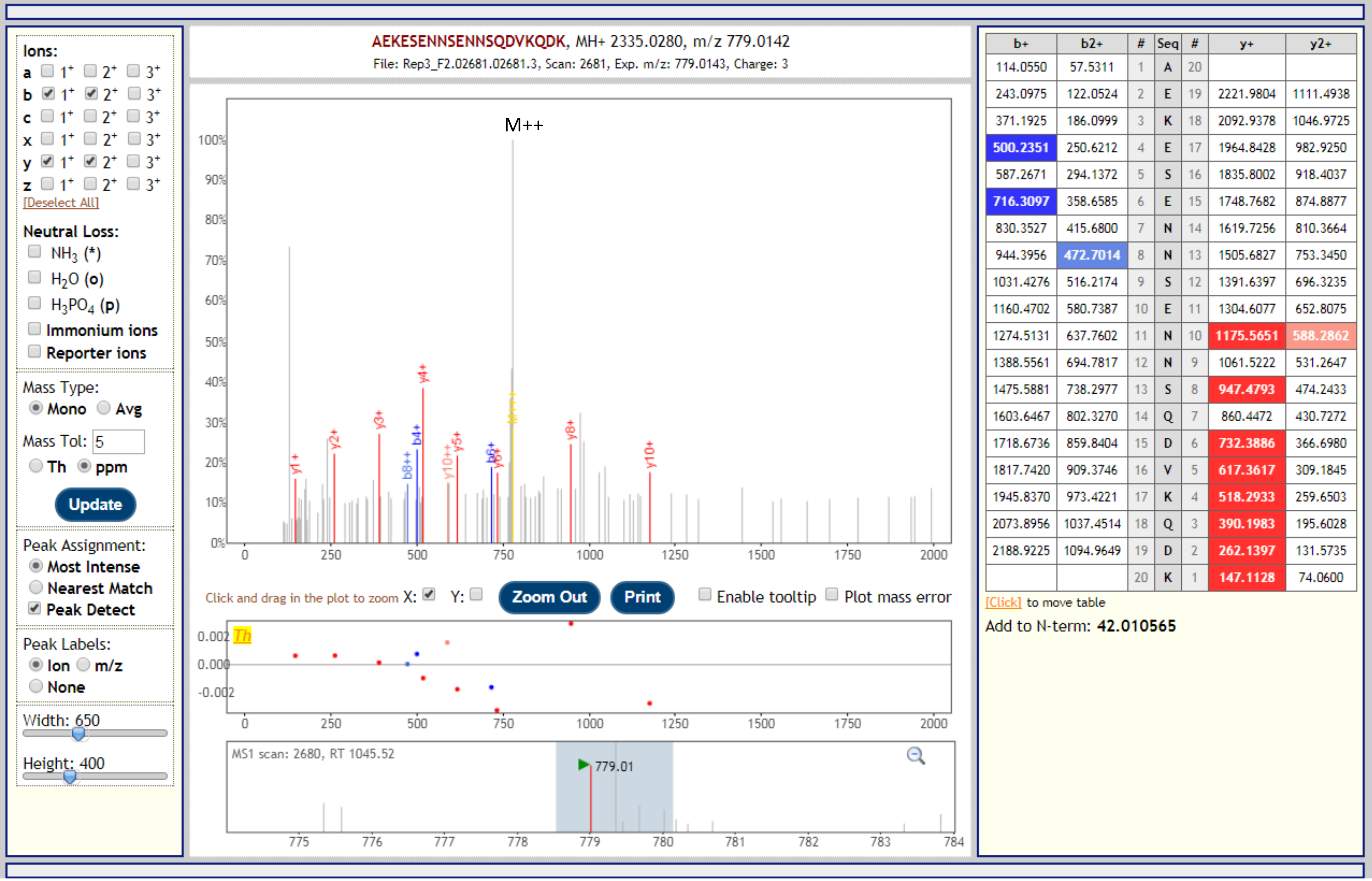


| b+ | b2+ | # | Seq | # | y+ | y2+ |
|---|---|---|---|---|---|---|
| 114.0550 | 57.5311 | 1 | A | 20 | | |
| 243.0975 | 122.0524 | 2 | E | 19 | 2221.9804 | 1111.4938 |
| 371.1925 | 186.0999 | 3 | K | 18 | 2092.9378 | 1046.9725 |
| 500.2351 | 250.6212 | 4 | E | 17 | 1964.8428 | 982.9250 |
| 587.2671 | 294.1372 | 5 | S | 16 | 1835.8002 | 918.4037 |
| 716.3097 | 358.6585 | 6 | E | 15 | 1748.7682 | 874.8877 |
| 830.3527 | 415.6800 | 7 | N | 14 | 1619.7256 | 810.3664 |
| 944.3956 | 472.7014 | 8 | N | 13 | 1505.6827 | 753.3450 |
| 1031.4276 | 516.2174 | 9 | S | 12 | 1391.6397 | 696.3235 |
| 1160.4702 | 580.7387 | 10 | E | 11 | 1304.6077 | 652.8075 |
| 1274.5131 | 637.7602 | 11 | N | 10 | 1175.5651 | 588.2862 |
| 1388.5561 | 694.7817 | 12 | N | 9 | 1061.5222 | 531.2647 |
| 1475.5881 | 738.2977 | 13 | S | 8 | 947.4793 | 474.2433 |
| 1603.6467 | 802.3270 | 14 | Q | 7 | 860.4472 | 430.7272 |
| 1718.6736 | 859.8404 | 15 | D | 6 | 732.3886 | 366.6980 |
| 1817.7420 | 909.3746 | 16 | V | 5 | 617.3617 | 309.1845 |
| 1945.8370 | 973.4221 | 17 | K | 4 | 518.2933 | 259.6503 |
| 2073.8956 | 1037.4514 | 18 | Q | 3 | 390.1983 | 195.6028 |
| 2188.9225 | 1094.9649 | 19 | D | 2 | 262.1397 | 131.5735 |
| | | 20 | K | 1 | 147.1128 | 74.0600 |

| Gene | Description | Peptide with cleaved & acetylated PEXEL |
|---|---|---|
| PBANKA_1003000 | liver specific protein 2 | RLI.AEKESENNSENNSENNSENNSENNSENNSENNSQDVK |

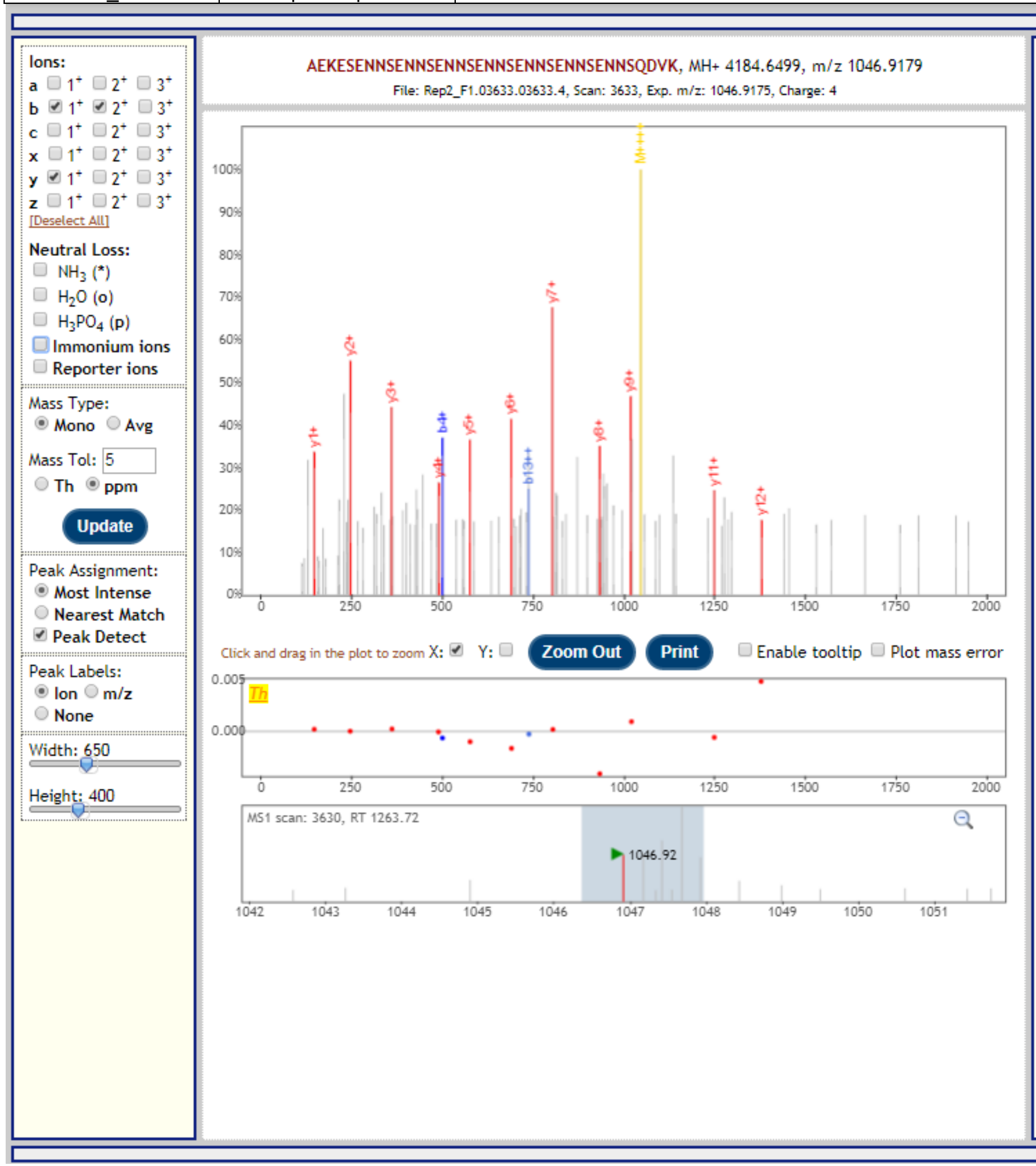


| b+ | b2+ | # | Seq | # | y+ |
|---|---|---|---|---|---|
| 114.0550 | 57.5311 | 1 | A | 37 | |
| 243.0975 | 122.0524 | 2 | E | 36 | 4071.6023 |
| 371.1925 | 186.0999 | 3 | K | 35 | 3942.5597 |
| 500.2351 | 250.6212 | 4 | E | 34 | 3814.4647 |
| 587.2671 | 294.1372 | 5 | S | 33 | 3685.4221 |
| 716.3097 | 358.6585 | 6 | E | 32 | 3598.3901 |
| 830.3527 | 415.6800 | 7 | N | 31 | 3469.3475 |
| 944.3956 | 472.7014 | 8 | N | 30 | 3355.3046 |
| 1031.4276 | 516.2174 | 9 | S | 29 | 3241.2616 |
| 1160.4702 | 580.7387 | 10 | E | 28 | 3154.2296 |
| 1274.5131 | 637.7602 | 11 | N | 27 | 3025.1870 |
| 1388.5561 | 694.7817 | 12 | N | 26 | 2911.1441 |
| 1475.5881 | 738.2977 | 13 | S | 25 | 2797.1011 |
| 1604.6307 | 802.8190 | 14 | E | 24 | 2710.0691 |
| 1718.6736 | 859.8404 | 15 | N | 23 | 2581.0265 |
| 1832.7165 | 916.8619 | 16 | N | 22 | 2466.9836 |
| 1919.7486 | 960.3779 | 17 | S | 21 | 2352.9407 |
| 2048.7912 | 1024.8992 | 18 | E | 20 | 2265.9086 |
| 2162.8341 | 1081.9207 | 19 | N | 19 | 2136.8661 |
| 2276.8770 | 1138.9421 | 20 | N | 18 | 2022.8231 |
| 2363.9090 | 1182.4582 | 21 | S | 17 | 1908.7802 |
| 2492.9516 | 1246.9795 | 22 | E | 16 | 1821.7482 |
| 2606.9946 | 1304.0009 | 23 | N | 15 | 1692.7056 |
| 2721.0375 | 1361.0224 | 24 | N | 14 | 1578.6626 |
| 2808.0695 | 1404.5384 | 25 | S | 13 | 1464.6197 |
| 2937.1121 | 1469.0597 | 26 | E | 12 | 1377.5877 |
| 3051.1550 | 1526.0812 | 27 | N | 11 | 1248.5451 |
| 3165.1980 | 1583.1026 | 28 | N | 10 | 1134.5022 |
| 3252.2300 | 1626.6186 | 29 | S | 9 | 1020.4592 |
| 3381.2726 | 1691.1399 | 30 | E | 8 | 933.4272 |
| 3495.3155 | 1748.1614 | 31 | N | 7 | 804.3846 |
| 3609.3584 | 1805.1829 | 32 | N | 6 | 690.3417 |
| 3696.3905 | 1848.6989 | 33 | S | 5 | 576.2988 |
| 3824.4490 | 1912.7282 | 34 | Q | 4 | 489.2667 |
| 3939.4760 | 1970.2416 | 35 | D | 3 | 361.2082 |
| 4038.5444 | 2019.7758 | 36 | V | 2 | 246.1812 |
| | | 37 | K | 1 | 147.1128 |

[Click] to move table

Add to N-term: 42.010565

| Gene | Description | Peptide with cleaved & acetylated PEXEL |
|---|---|---|
| PBANKA_1003000 | liver specific protein 2 | RLI.AEKESENNSENNSENNSENNSENNSENNSENNSQDVK |

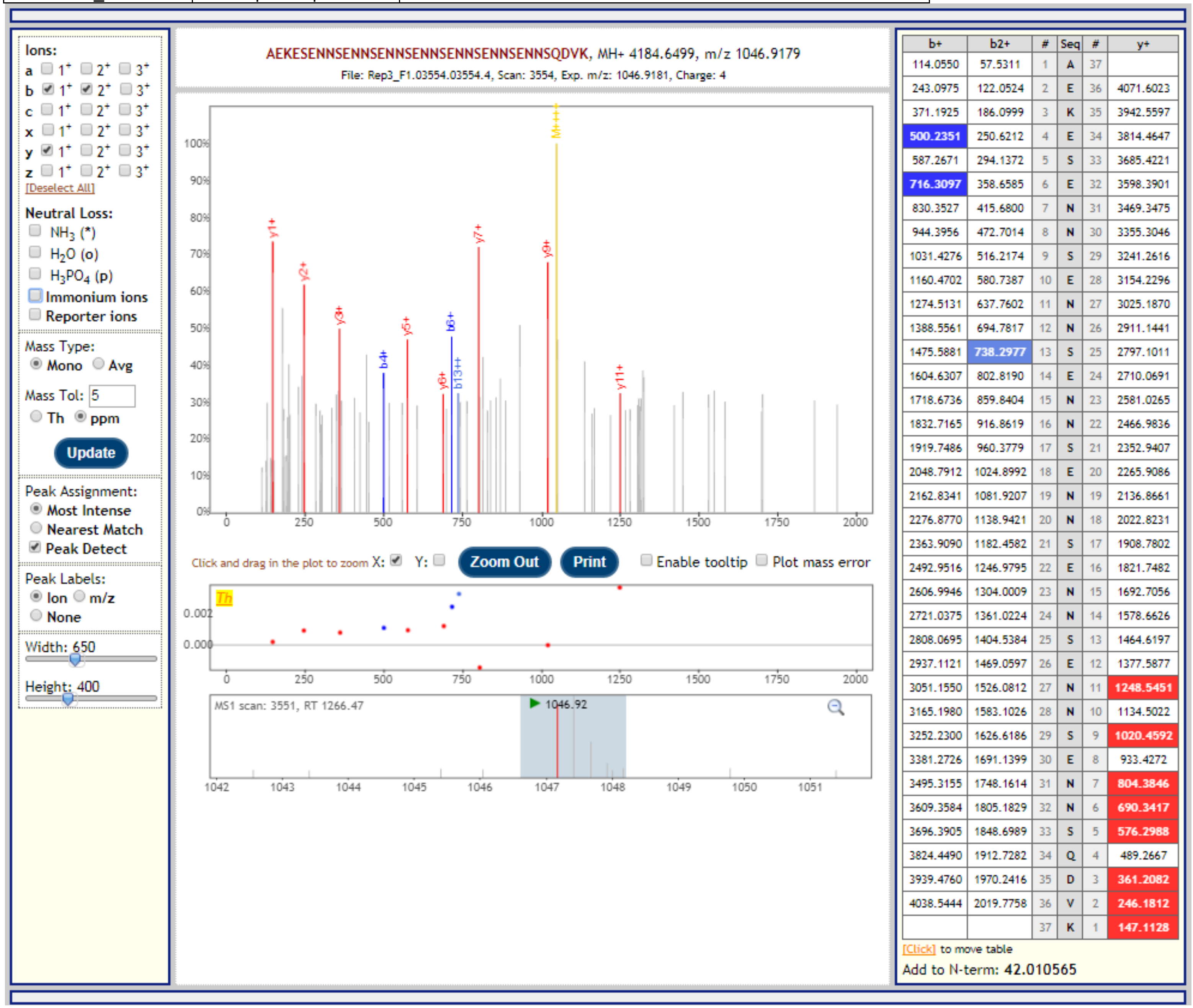


| b+ | b2+ | # | Seq | # | y+ |
|---|---|---|---|---|---|
| 114.0550 | 57.5311 | 1 | A | 37 | |
| 243.0975 | 122.0524 | 2 | E | 36 | 4071.6023 |
| 371.1925 | 186.0999 | 3 | K | 35 | 3942.5597 |
| 500.2351 | 250.6212 | 4 | E | 34 | 3814.4647 |
| 587.2671 | 294.1372 | 5 | S | 33 | 3685.4221 |
| 716.3097 | 358.6585 | 6 | E | 32 | 3598.3901 |
| 830.3527 | 415.6800 | 7 | N | 31 | 3469.3475 |
| 944.3956 | 472.7014 | 8 | N | 30 | 3355.3046 |
| 1031.4276 | 516.2174 | 9 | S | 29 | 3241.2616 |
| 1160.4702 | 580.7387 | 10 | E | 28 | 3154.2296 |
| 1274.5131 | 637.7602 | 11 | N | 27 | 3025.1870 |
| 1388.5561 | 694.7817 | 12 | N | 26 | 2911.1441 |
| 1475.5881 | 738.2977 | 13 | S | 25 | 2797.1011 |
| 1604.6307 | 802.8190 | 14 | E | 24 | 2710.0691 |
| 1718.6736 | 859.8404 | 15 | N | 23 | 2581.0265 |
| 1832.7165 | 916.8619 | 16 | N | 22 | 2466.9836 |
| 1919.7486 | 960.3779 | 17 | S | 21 | 2352.9407 |
| 2048.7912 | 1024.8992 | 18 | E | 20 | 2265.9086 |
| 2162.8341 | 1081.9207 | 19 | N | 19 | 2136.8661 |
| 2276.8770 | 1138.9421 | 20 | N | 18 | 2022.8231 |
| 2363.9090 | 1182.4582 | 21 | S | 17 | 1908.7802 |
| 2492.9516 | 1246.9795 | 22 | E | 16 | 1821.7482 |
| 2606.9946 | 1304.0009 | 23 | N | 15 | 1692.7056 |
| 2721.0375 | 1361.0224 | 24 | N | 14 | 1578.6626 |
| 2808.0695 | 1404.5384 | 25 | S | 13 | 1464.6197 |
| 2937.1121 | 1469.0597 | 26 | E | 12 | 1377.5877 |
| 3051.1550 | 1526.0812 | 27 | N | 11 | 1248.5451 |
| 3165.1980 | 1583.1026 | 28 | N | 10 | 1134.5022 |
| 3252.2300 | 1626.6186 | 29 | S | 9 | 1020.4592 |
| 3381.2726 | 1691.1399 | 30 | E | 8 | 933.4272 |
| 3495.3155 | 1748.1614 | 31 | N | 7 | 804.3846 |
| 3609.3584 | 1805.1829 | 32 | N | 6 | 690.3417 |
| 3696.3905 | 1848.6989 | 33 | S | 5 | 576.2988 |
| 3824.4490 | 1912.7282 | 34 | Q | 4 | 489.2667 |
| 3939.4760 | 1970.2416 | 35 | D | 3 | 361.2082 |
| 4038.5444 | 2019.7758 | 36 | V | 2 | 246.1812 |
| | | 37 | K | 1 | 147.1128 |

| Gene | Description | Peptide with cleaved & acetylated PEXEL |
|---|---|---|
| PBANKA_1003000 | liver specific protein 2 | RLI.AEKESENNSENNSENNSENNSENNSENNSQDVK |

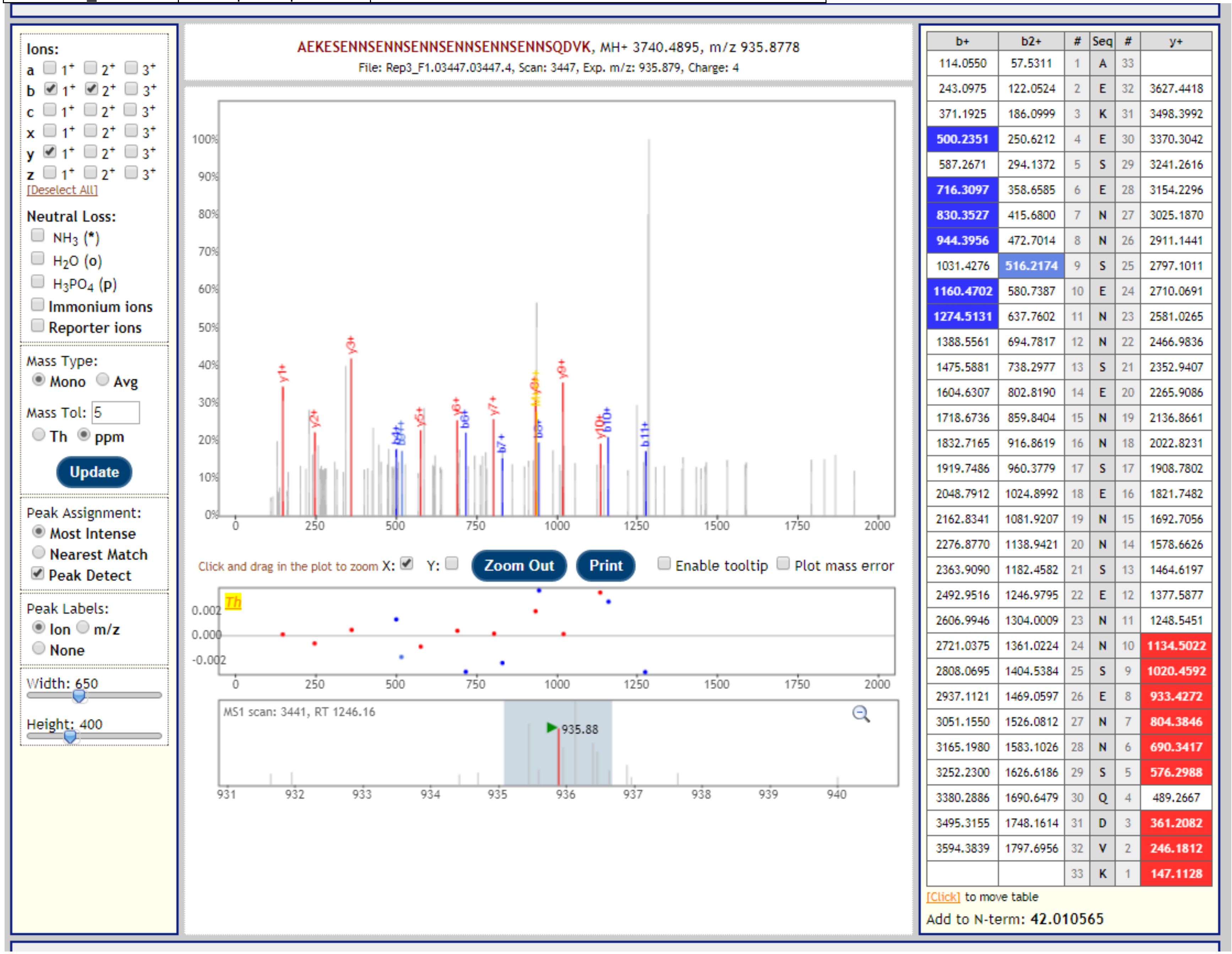


| b+ | b2+ | # | Seq | # | y+ |
|---|---|---|---|---|---|
| 114.0550 | 57.5311 | 1 | A | 33 | |
| 243.0975 | 122.0524 | 2 | E | 32 | 3627.4418 |
| 371.1925 | 186.0999 | 3 | K | 31 | 3498.3992 |
| 500.2351 | 250.6212 | 4 | E | 30 | 3370.3042 |
| 587.2671 | 294.1372 | 5 | S | 29 | 3241.2616 |
| 716.3097 | 358.6585 | 6 | E | 28 | 3154.2296 |
| 830.3527 | 415.6800 | 7 | N | 27 | 3025.1870 |
| 944.3956 | 472.7014 | 8 | N | 26 | 2911.1441 |
| 1031.4276 | 516.2174 | 9 | S | 25 | 2797.1011 |
| 1160.4702 | 580.7387 | 10 | E | 24 | 2710.0691 |
| 1274.5131 | 637.7602 | 11 | N | 23 | 2581.0265 |
| 1388.5561 | 694.7817 | 12 | N | 22 | 2466.9836 |
| 1475.5881 | 738.2977 | 13 | S | 21 | 2352.9407 |
| 1604.6307 | 802.8190 | 14 | E | 20 | 2265.9086 |
| 1718.6736 | 859.8404 | 15 | N | 19 | 2136.8661 |
| 1832.7165 | 916.8619 | 16 | N | 18 | 2022.8231 |
| 1919.7486 | 960.3779 | 17 | S | 17 | 1908.7802 |
| 2048.7912 | 1024.8992 | 18 | E | 16 | 1821.7482 |
| 2162.8341 | 1081.9207 | 19 | N | 15 | 1692.7056 |
| 2276.8770 | 1138.9421 | 20 | N | 14 | 1578.6626 |
| 2363.9090 | 1182.4582 | 21 | S | 13 | 1464.6197 |
| 2492.9516 | 1246.9795 | 22 | E | 12 | 1377.5877 |
| 2606.9946 | 1304.0009 | 23 | N | 11 | 1248.5451 |
| 2721.0375 | 1361.0224 | 24 | N | 10 | 1134.5022 |
| 2808.0695 | 1404.5384 | 25 | S | 9 | 1020.4592 |
| 2937.1121 | 1469.0597 | 26 | E | 8 | 933.4272 |
| 3051.1550 | 1526.0812 | 27 | N | 7 | 804.3846 |
| 3165.1980 | 1583.1026 | 28 | N | 6 | 690.3417 |
| 3252.2300 | 1626.6186 | 29 | S | 5 | 576.2988 |
| 3380.2886 | 1690.6479 | 30 | Q | 4 | 489.2667 |
| 3495.3155 | 1748.1614 | 31 | D | 3 | 361.2082 |
| 3594.3839 | 1797.6956 | 32 | V | 2 | 246.1812 |
| | | 33 | K | 1 | 147.1128 |

| Gene | Description | Peptide with cleaved & acetylated PEXEL |
|---|---|---|
| PBANKA_1003000 | liver specific protein 2 | RLI.AEKESENNSENNSENNSENNSENNSENNSQDVK |

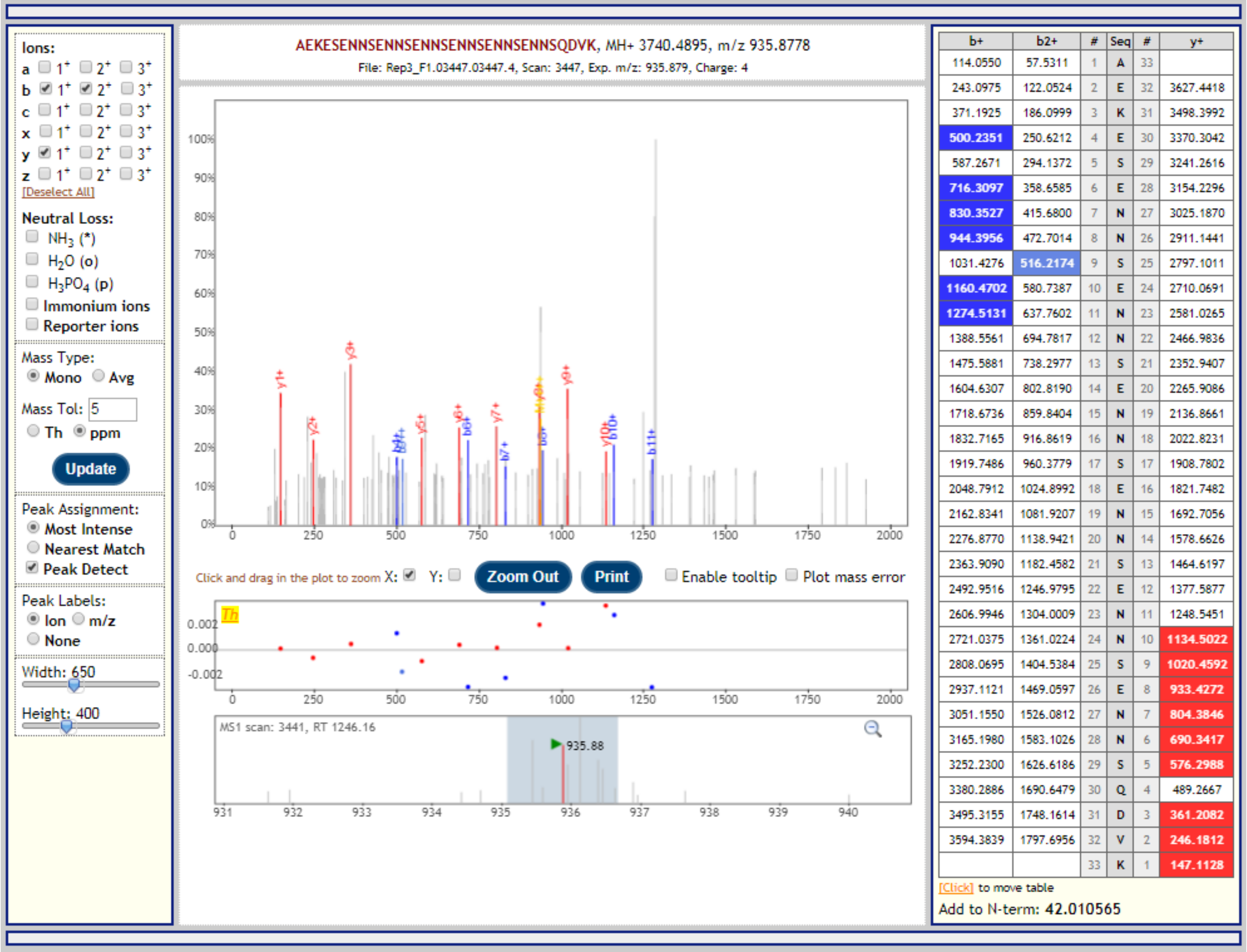


| b+ | b2+ | # | Seq | # | y+ |
|---|---|---|---|---|---|
| 114.0550 | 57.5311 | 1 | A | 33 | |
| 243.0975 | 122.0524 | 2 | E | 32 | 3627.4418 |
| 371.1925 | 186.0999 | 3 | K | 31 | 3498.3992 |
| 500.2351 | 250.6212 | 4 | E | 30 | 3370.3042 |
| 587.2671 | 294.1372 | 5 | S | 29 | 3241.2616 |
| 716.3097 | 358.6585 | 6 | E | 28 | 3154.2296 |
| 830.3527 | 415.6800 | 7 | N | 27 | 3025.1870 |
| 944.3956 | 472.7014 | 8 | N | 26 | 2911.1441 |
| 1031.4276 | 516.2174 | 9 | S | 25 | 2797.1011 |
| 1160.4702 | 580.7387 | 10 | E | 24 | 2710.0691 |
| 1274.5131 | 637.7602 | 11 | N | 23 | 2581.0265 |
| 1388.5561 | 694.7817 | 12 | N | 22 | 2466.9836 |
| 1475.5881 | 738.2977 | 13 | S | 21 | 2352.9407 |
| 1604.6307 | 802.8190 | 14 | E | 20 | 2265.9086 |
| 1718.6736 | 859.8404 | 15 | N | 19 | 2136.8661 |
| 1832.7165 | 916.8619 | 16 | N | 18 | 2022.8231 |
| 1919.7486 | 960.3779 | 17 | S | 17 | 1908.7802 |
| 2048.7912 | 1024.8992 | 18 | E | 16 | 1821.7482 |
| 2162.8341 | 1081.9207 | 19 | N | 15 | 1692.7056 |
| 2276.8770 | 1138.9421 | 20 | N | 14 | 1578.6626 |
| 2363.9090 | 1182.4582 | 21 | S | 13 | 1464.6197 |
| 2492.9516 | 1246.9795 | 22 | E | 12 | 1377.5877 |
| 2606.9946 | 1304.0009 | 23 | N | 11 | 1248.5451 |
| 2721.0375 | 1361.0224 | 24 | N | 10 | 1134.5022 |
| 2808.0695 | 1404.5384 | 25 | S | 9 | 1020.4592 |
| 2937.1121 | 1469.0597 | 26 | E | 8 | 933.4272 |
| 3051.1550 | 1526.0812 | 27 | N | 7 | 804.3846 |
| 3165.1980 | 1583.1026 | 28 | N | 6 | 690.3417 |
| 3252.2300 | 1626.6186 | 29 | S | 5 | 576.2988 |
| 3380.2886 | 1690.6479 | 30 | Q | 4 | 489.2667 |
| 3495.3155 | 1748.1614 | 31 | D | 3 | 361.2082 |
| 3594.3839 | 1797.6956 | 32 | V | 2 | 246.1812 |
| | | 33 | K | 1 | 147.1128 |

| Gene | Description | Peptide with cleaved & acetylated PEXEL |
|---|---|---|
| PBANKA_1003000 | liver specific protein 2 | RLI.AEKESENNSENNSENNSENNSQDVK |

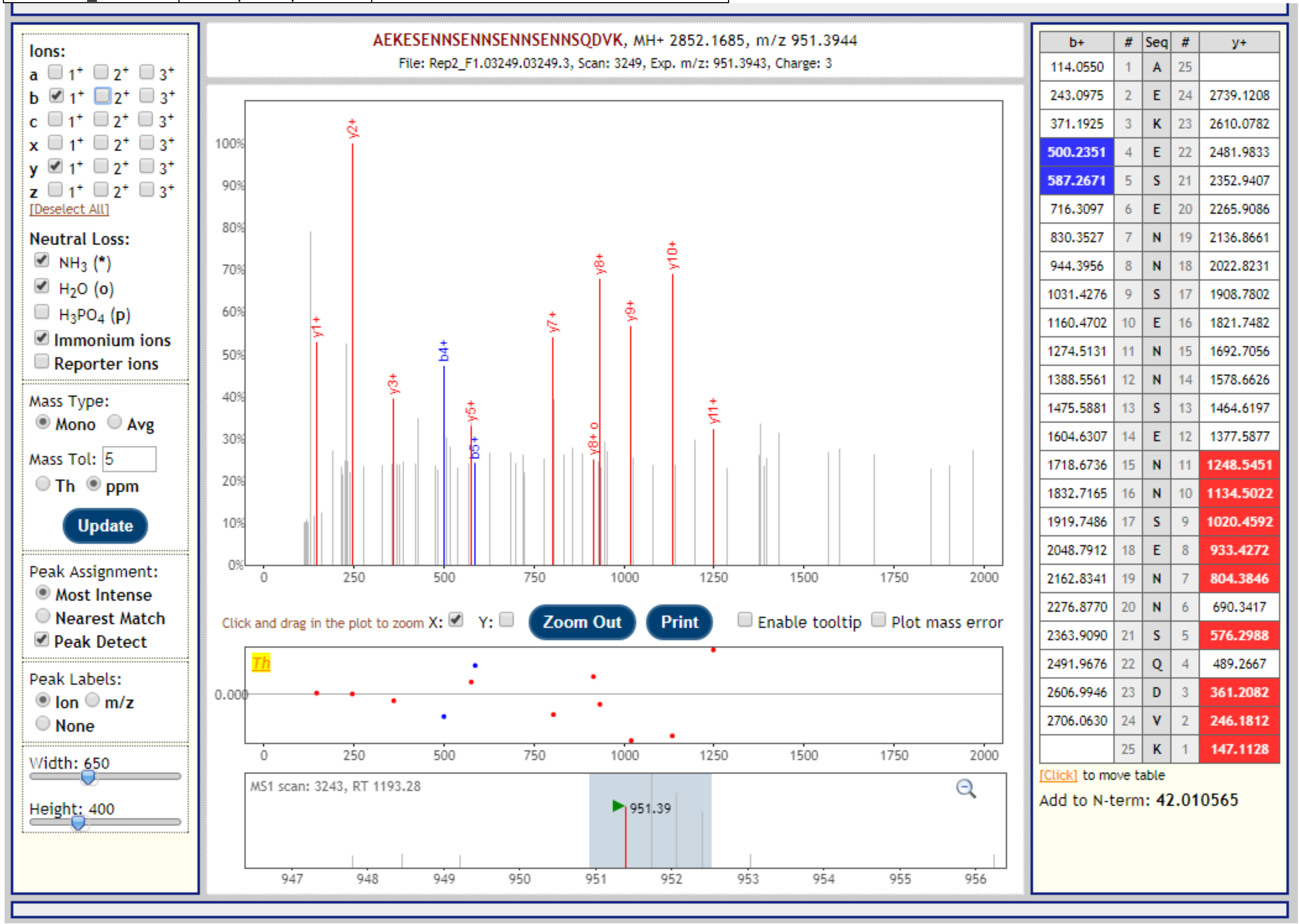

| Gene | Description | Peptide with cleaved & acetylated PEXEL |
|---|---|---|
| PBANKA_1003000 | liver specific protein 2 | RLI.AEKESENNSENNSENNSQDVK |

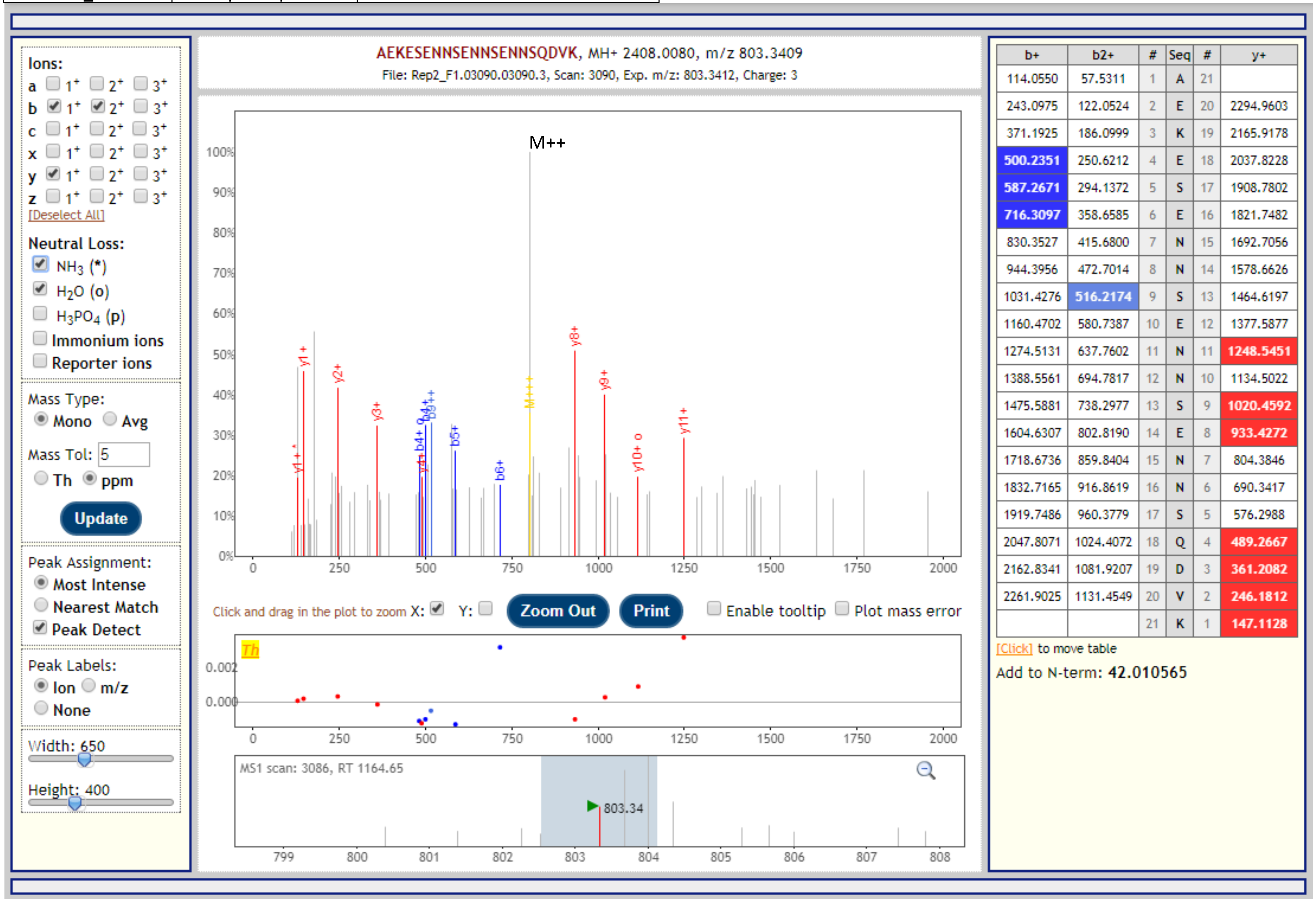


| b+ | b2+ | # | Seq | # | y+ |
|---|---|---|---|---|---|
| 114.0550 | 57.5311 | 1 | A | 21 | |
| 243.0975 | 122.0524 | 2 | E | 20 | 2294.9603 |
| 371.1925 | 186.0999 | 3 | K | 19 | 2165.9178 |
| 500.2351 | 250.6212 | 4 | E | 18 | 2037.8228 |
| 587.2671 | 294.1372 | 5 | S | 17 | 1908.7802 |
| 716.3097 | 358.6585 | 6 | E | 16 | 1821.7482 |
| 830.3527 | 415.6800 | 7 | N | 15 | 1692.7056 |
| 944.3956 | 472.7014 | 8 | N | 14 | 1578.6626 |
| 1031.4276 | 516.2174 | 9 | S | 13 | 1464.6197 |
| 1160.4702 | 580.7387 | 10 | E | 12 | 1377.5877 |
| 1274.5131 | 637.7602 | 11 | N | 11 | 1248.5451 |
| 1388.5561 | 694.7817 | 12 | N | 10 | 1134.5022 |
| 1475.5881 | 738.2977 | 13 | S | 9 | 1020.4592 |
| 1604.6307 | 802.8190 | 14 | E | 8 | 933.4272 |
| 1718.6736 | 859.8404 | 15 | N | 7 | 804.3846 |
| 1832.7165 | 916.8619 | 16 | N | 6 | 690.3417 |
| 1919.7486 | 960.3779 | 17 | S | 5 | 576.2988 |
| 2047.8071 | 1024.4072 | 18 | Q | 4 | 489.2667 |
| 2162.8341 | 1081.9207 | 19 | D | 3 | 361.2082 |
| 2261.9025 | 1131.4549 | 20 | V | 2 | 246.1812 |
| | | 21 | K | 1 | 147.1128 |

| Gene | Description | Peptide with cleaved & acetylated PEXEL |
|---|---|---|
| PBANKA_1328000 | serine/threonine protein phosphatase UIS2 | RVL.QEQNEDIKDDNDENDEGDEEDEYYSYLK |

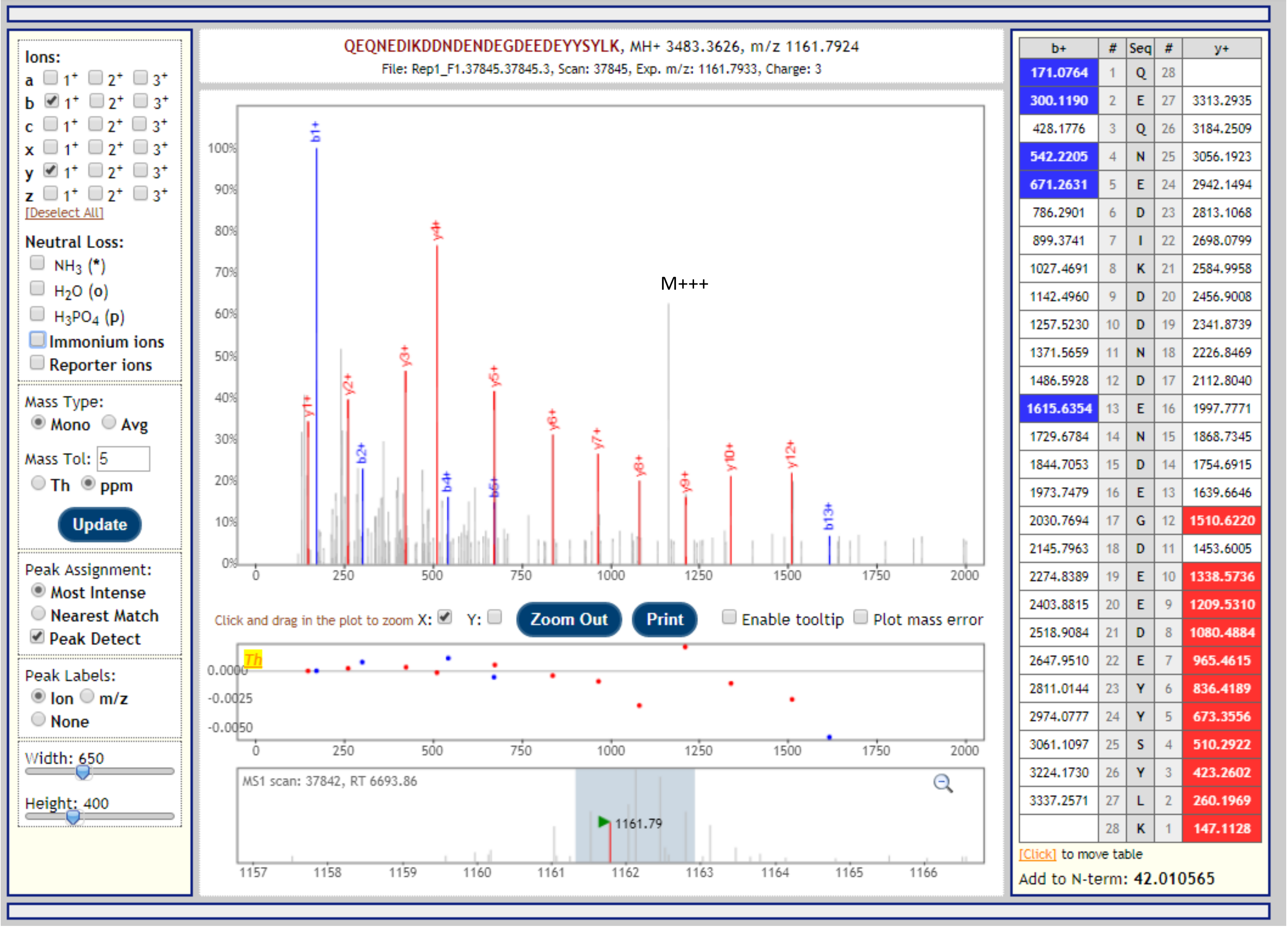


| b+ | # | Seq | # | y+ |
|---|---|---|---|---|
| 171.0764 | 1 | Q | 28 | |
| 300.1190 | 2 | E | 27 | 3313.2935 |
| 428.1776 | 3 | Q | 26 | 3184.2509 |
| 542.2205 | 4 | N | 25 | 3056.1923 |
| 671.2631 | 5 | E | 24 | 2942.1494 |
| 786.2901 | 6 | D | 23 | 2813.1068 |
| 899.3741 | 7 | I | 22 | 2698.0799 |
| 1027.4691 | 8 | K | 21 | 2584.9958 |
| 1142.4960 | 9 | D | 20 | 2456.9008 |
| 1257.5230 | 10 | D | 19 | 2341.8739 |
| 1371.5659 | 11 | N | 18 | 2226.8469 |
| 1486.5928 | 12 | D | 17 | 2112.8040 |
| 1615.6354 | 13 | E | 16 | 1997.7771 |
| 1729.6784 | 14 | N | 15 | 1868.7345 |
| 1844.7053 | 15 | D | 14 | 1754.6915 |
| 1973.7479 | 16 | E | 13 | 1639.6646 |
| 2030.7694 | 17 | G | 12 | 1510.6220 |
| 2145.7963 | 18 | D | 11 | 1453.6005 |
| 2274.8389 | 19 | E | 10 | 1338.5736 |
| 2403.8815 | 20 | E | 9 | 1209.5310 |
| 2518.9084 | 21 | D | 8 | 1080.4884 |
| 2647.9510 | 22 | E | 7 | 965.4615 |
| 2811.0144 | 23 | Y | 6 | 836.4189 |
| 2974.0777 | 24 | Y | 5 | 673.3556 |
| 3061.1097 | 25 | S | 4 | 510.2922 |
| 3224.1730 | 26 | Y | 3 | 423.2602 |
| 3337.2571 | 27 | L | 2 | 260.1969 |
| | 28 | K | 1 | 147.1128 |

| Gene | Description | Peptide with cleaved & acetylated PEXEL |
|---|---|---|
| PBANKA_1328000 | serine/threonine protein phosphatase UIS2 | RVL.QEQNEDIKDDNDENDEGDEEDEYYSYLK |

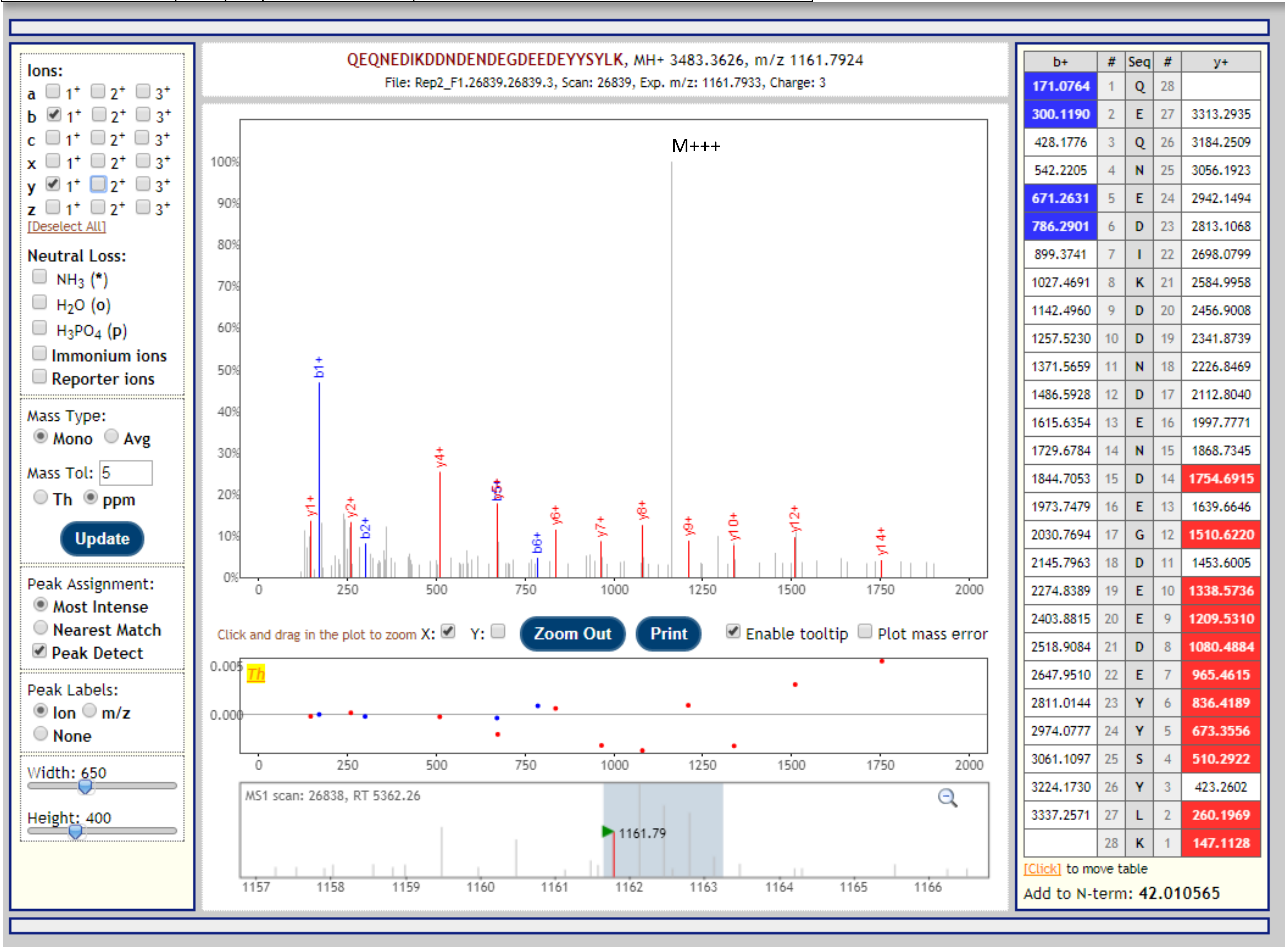


| b+ | # | Seq | # | y+ |
|---|---|---|---|---|
| 171.0764 | 1 | Q | 28 | |
| 300.1190 | 2 | E | 27 | 3313.2935 |
| 428.1776 | 3 | Q | 26 | 3184.2509 |
| 542.2205 | 4 | N | 25 | 3056.1923 |
| 671.2631 | 5 | E | 24 | 2942.1494 |
| 786.2901 | 6 | D | 23 | 2813.1068 |
| 899.3741 | 7 | I | 22 | 2698.0799 |
| 1027.4691 | 8 | K | 21 | 2584.9958 |
| 1142.4960 | 9 | D | 20 | 2456.9008 |
| 1257.5230 | 10 | D | 19 | 2341.8739 |
| 1371.5659 | 11 | N | 18 | 2226.8469 |
| 1486.5928 | 12 | D | 17 | 2112.8040 |
| 1615.6354 | 13 | E | 16 | 1997.7771 |
| 1729.6784 | 14 | N | 15 | 1868.7345 |
| 1844.7053 | 15 | D | 14 | 1754.6915 |
| 1973.7479 | 16 | E | 13 | 1639.6646 |
| 2030.7694 | 17 | G | 12 | 1510.6220 |
| 2145.7963 | 18 | D | 11 | 1453.6005 |
| 2274.8389 | 19 | E | 10 | 1338.5736 |
| 2403.8815 | 20 | E | 9 | 1209.5310 |
| 2518.9084 | 21 | D | 8 | 1080.4884 |
| 2647.9510 | 22 | E | 7 | 965.4615 |
| 2811.0144 | 23 | Y | 6 | 836.4189 |
| 2974.0777 | 24 | Y | 5 | 673.3556 |
| 3061.1097 | 25 | S | 4 | 510.2922 |
| 3224.1730 | 26 | Y | 3 | 423.2602 |
| 3337.2571 | 27 | L | 2 | 260.1969 |
| | 28 | K | 1 | 147.1128 |

| Gene | Description | Peptide with cleaved & acetylated PEXEL |
|---|---|---|
| PBANKA_1328000 | serine/threonine protein phosphatase UIS2 | RVL.QEQNEDIKDDNDENDEGDEEDEYYSYLK |

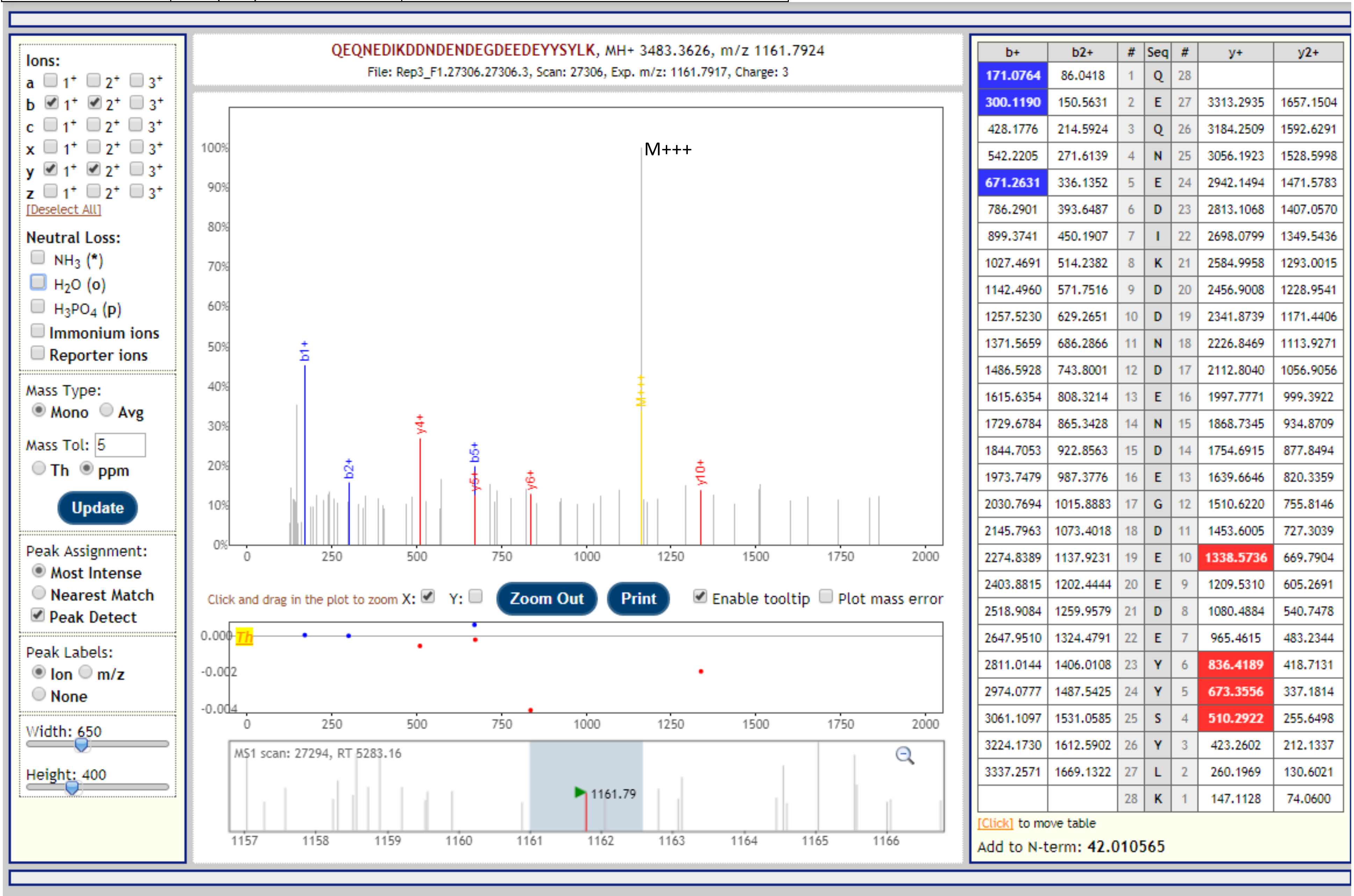

| Gene | Description | Peptide with cleaved & acetylated PEXEL |
|---|---|---|
| PBANKA_1365500 | exported protein IBIS1 | RIL.SELDQNKDQNLGYK |

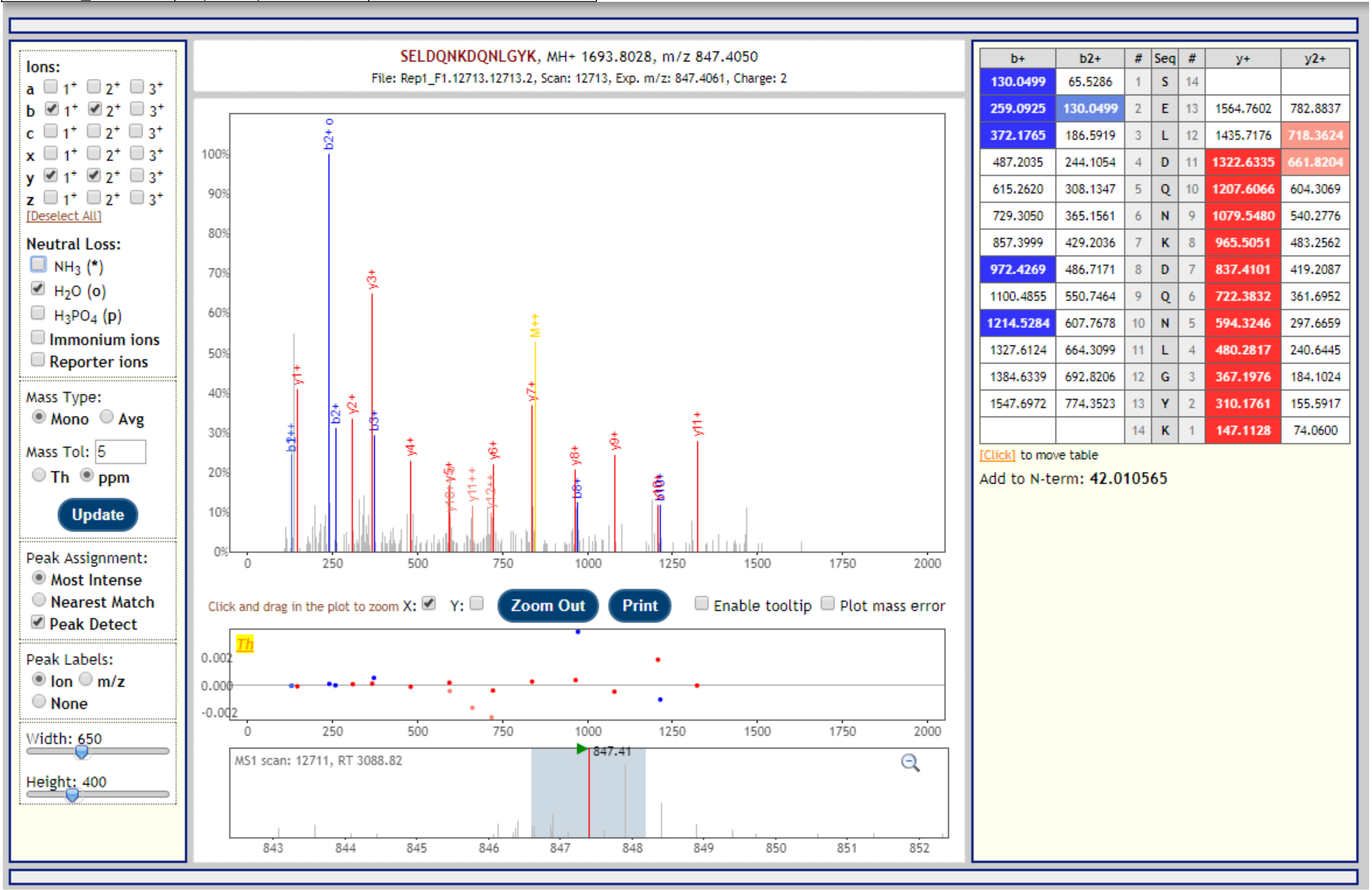


| b+ | b2+ | # | Seq | # | y+ | y2+ |
|---|---|---|---|---|---|---|
| 130.0499 | 65.5286 | 1 | S | 14 | | |
| 259.0925 | 130.0499 | 2 | E | 13 | 1564.7602 | 782.8837 |
| 372.1765 | 186.5919 | 3 | L | 12 | 1435.7176 | 718.3624 |
| 487.2035 | 244.1054 | 4 | D | 11 | 1322.6335 | 661.8204 |
| 615.2620 | 308.1347 | 5 | Q | 10 | 1207.6066 | 604.3069 |
| 729.3050 | 365.1561 | 6 | N | 9 | 1079.5480 | 540.2776 |
| 857.3999 | 429.2036 | 7 | K | 8 | 965.5051 | 483.2562 |
| 972.4269 | 486.7171 | 8 | D | 7 | 837.4101 | 419.2087 |
| 1100.4855 | 550.7464 | 9 | Q | 6 | 722.3832 | 361.6952 |
| 1214.5284 | 607.7678 | 10 | N | 5 | 594.3246 | 297.6659 |
| 1327.6124 | 664.3099 | 11 | L | 4 | 480.2817 | 240.6445 |
| 1384.6339 | 692.8206 | 12 | G | 3 | 367.1976 | 184.1024 |
| 1547.6972 | 774.3523 | 13 | Y | 2 | 310.1761 | 155.5917 |
| | | 14 | K | 1 | 147.1128 | 74.0600 |

| Gene | Description | Peptide with cleaved & acetylated PEXEL |
|---|---|---|
| PBANKA_1365500 | exported protein IBIS1 | RIL.SELDQNKDQNLGYK |

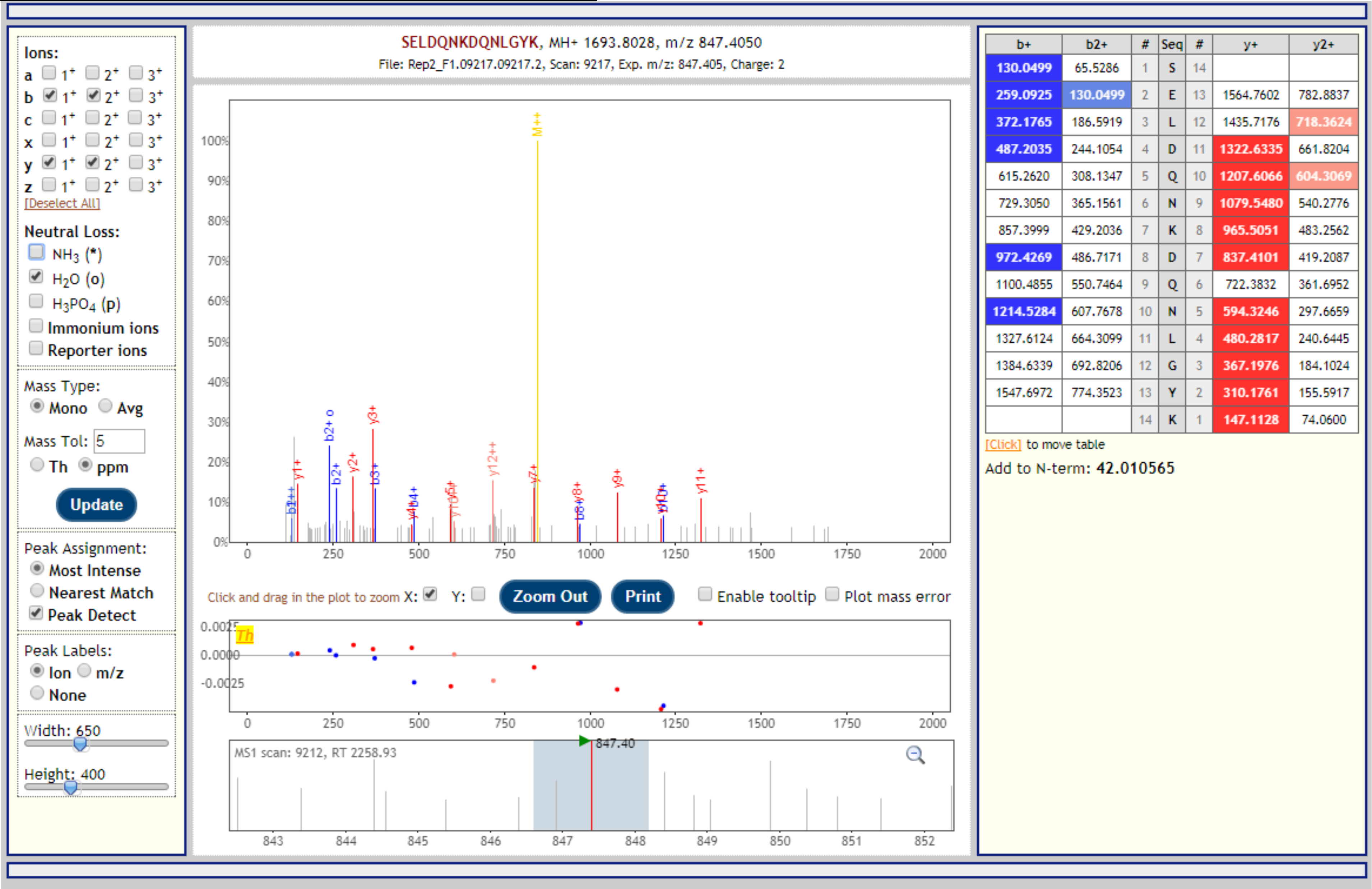

| Gene | Description | Peptide with cleaved & acetylated PEXEL |
|---|---|---|
| PBANKA_1365500 | exported protein IBIS1 | RIL.SELDQNKDQNLGYK |

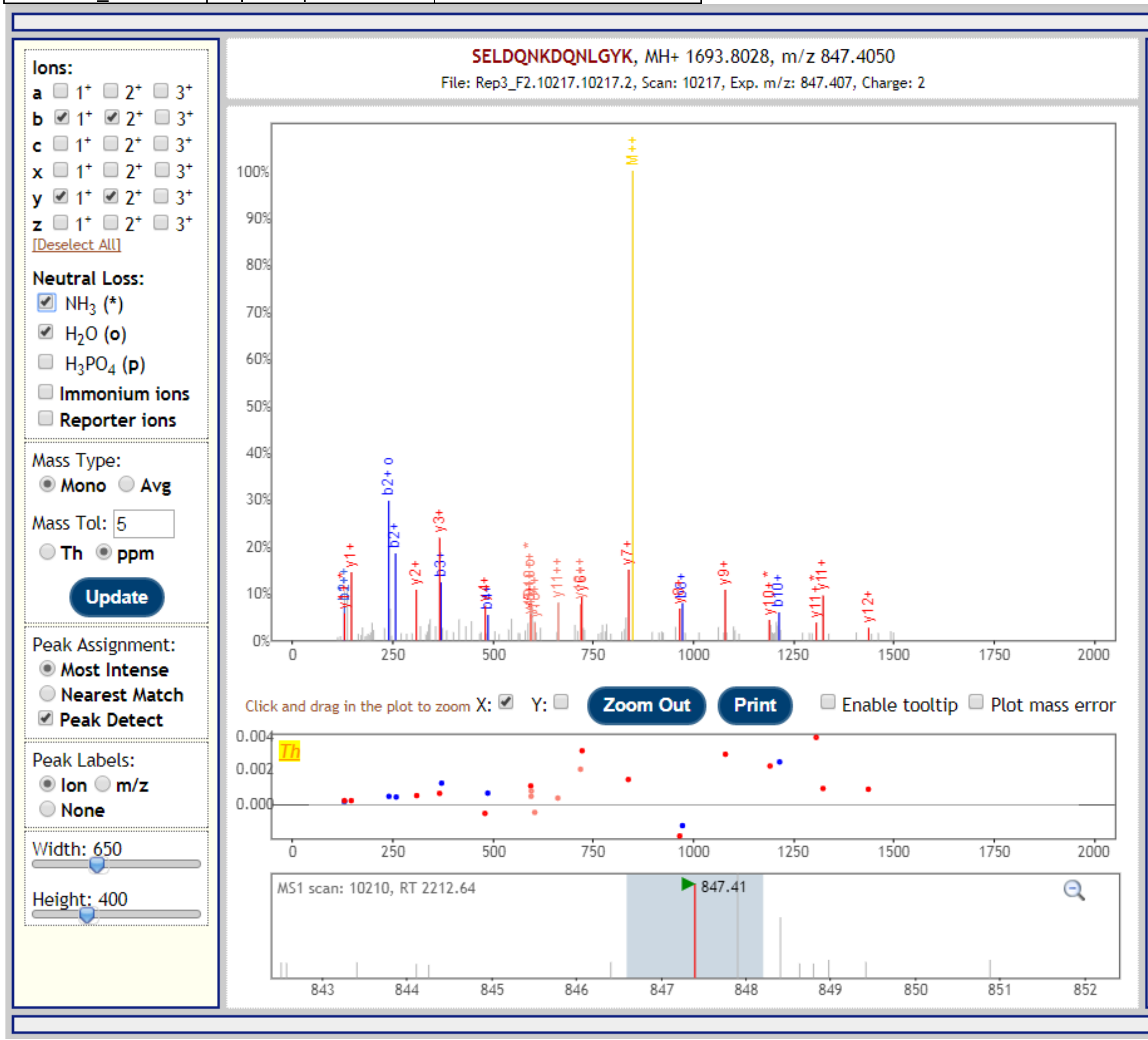


| b+ | b2+ | # | Seq | # | y+ | y2+ |
|---|---|---|---|---|---|---|
| 130.0499 | 65.5286 | 1 | S | 14 | | |
| 259.0925 | 130.0499 | 2 | E | 13 | 1564.7602 | 782.8837 |
| 372.1765 | 186.5919 | 3 | L | 12 | 1435.7176 | 718.3624 |
| 487.2035 | 244.1054 | 4 | D | 11 | 1322.6335 | 661.8204 |
| 615.2620 | 308.1347 | 5 | Q | 10 | 1207.6066 | 604.3069 |
| 729.3050 | 365.1561 | 6 | N | 9 | 1079.5480 | 540.2776 |
| 857.3999 | 429.2036 | 7 | K | 8 | 965.5051 | 483.2562 |
| 972.4269 | 486.7171 | 8 | D | 7 | 837.4101 | 419.2087 |
| 1100.4855 | 550.7464 | 9 | Q | 6 | 722.3832 | 361.6952 |
| 1214.5284 | 607.7678 | 10 | N | 5 | 594.3246 | 297.6659 |
| 1327.6124 | 664.3099 | 11 | L | 4 | 480.2817 | 240.6445 |
| 1384.6339 | 692.8206 | 12 | G | 3 | 367.1976 | 184.1024 |
| 1547.6972 | 774.3523 | 13 | Y | 2 | 310.1761 | 155.5917 |
| | | 14 | K | 1 | 147.1128 | 74.0600 |

[Click] to move table

Add to N-term: **42.010565**

| Gene | Description | Peptide with cleaved & acetylated PEXEL |
|---|---|---|
| PBANKA_1465000 | fam-c protein | RSL.SEHSTEDKEETVTISDNK |

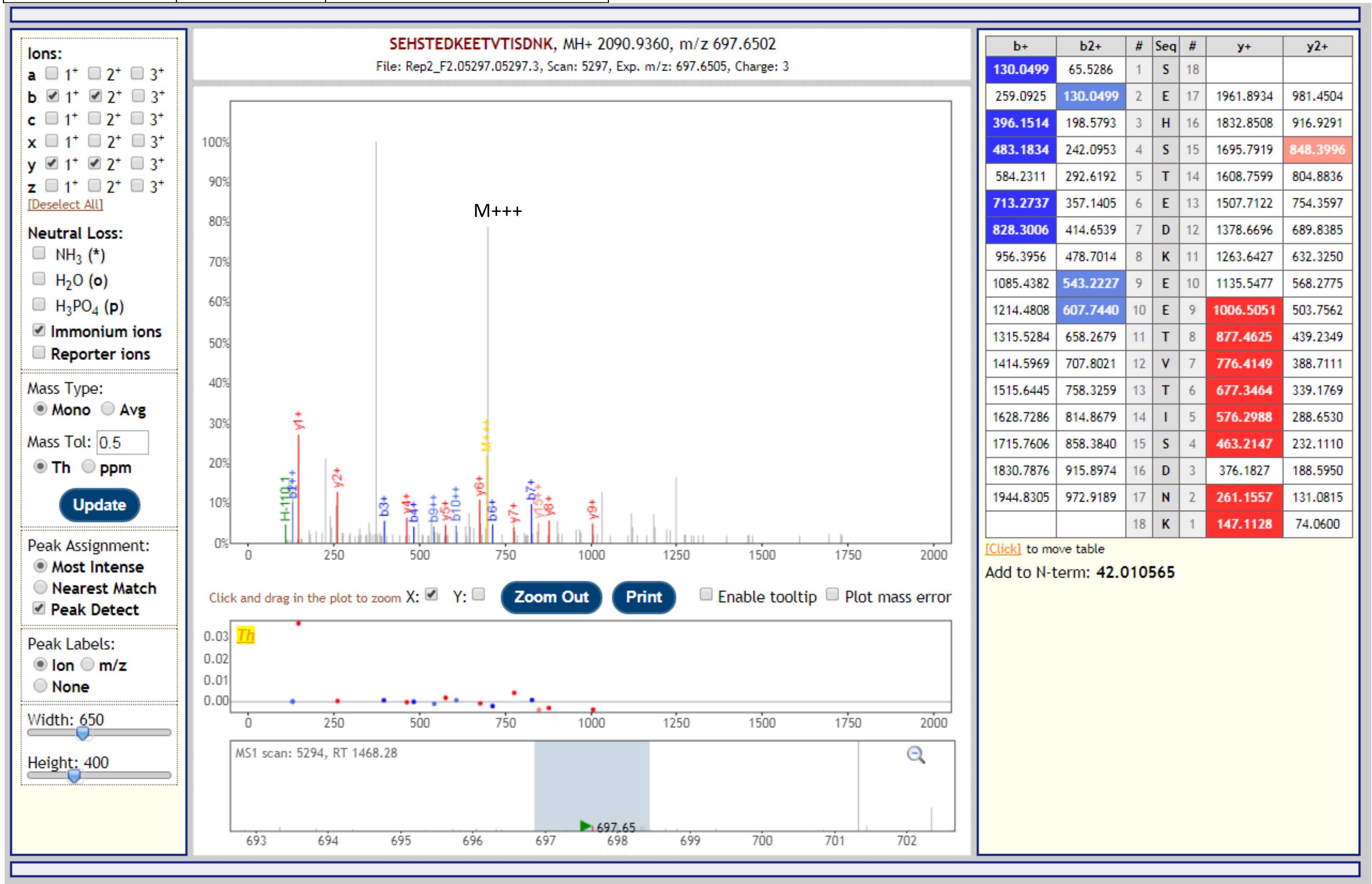


| b+ | b2+ | # | Seq | # | y+ | y2+ |
|---|---|---|---|---|---|---|
| 130.0499 | 65.5286 | 1 | S | 18 | | |
| 259.0925 | 130.0499 | 2 | E | 17 | 1961.8934 | 981.4504 |
| 396.1514 | 198.5793 | 3 | H | 16 | 1832.8508 | 916.9291 |
| 483.1834 | 242.0953 | 4 | S | 15 | 1695.7919 | 848.3996 |
| 584.2311 | 292.6192 | 5 | T | 14 | 1608.7599 | 804.8836 |
| 713.2737 | 357.1405 | 6 | E | 13 | 1507.7122 | 754.3597 |
| 828.3006 | 414.6539 | 7 | D | 12 | 1378.6696 | 689.8385 |
| 956.3956 | 478.7014 | 8 | K | 11 | 1263.6427 | 632.3250 |
| 1085.4382 | 543.2227 | 9 | E | 10 | 1135.5477 | 568.2775 |
| 1214.4808 | 607.7440 | 10 | E | 9 | 1006.5051 | 503.7562 |
| 1315.5284 | 658.2679 | 11 | T | 8 | 877.4625 | 439.2349 |
| 1414.5969 | 707.8021 | 12 | V | 7 | 776.4149 | 388.7111 |
| 1515.6445 | 758.3259 | 13 | T | 6 | 677.3464 | 339.1769 |
| 1628.7286 | 814.8679 | 14 | I | 5 | 576.2988 | 288.6530 |
| 1715.7606 | 858.3840 | 15 | S | 4 | 463.2147 | 232.1110 |
| 1830.7876 | 915.8974 | 16 | D | 3 | 376.1827 | 188.5950 |
| 1944.8305 | 972.9189 | 17 | N | 2 | 261.1557 | 131.0815 |
| | | 18 | K | 1 | 147.1128 | 74.0600 |

| Gene | Description | Peptide with cleaved & acetylated PEXEL |
|---|---|---|
| PBANKA_1465000 | fam-c protein | RSL.SEHSTEDKEETVTISDNK |

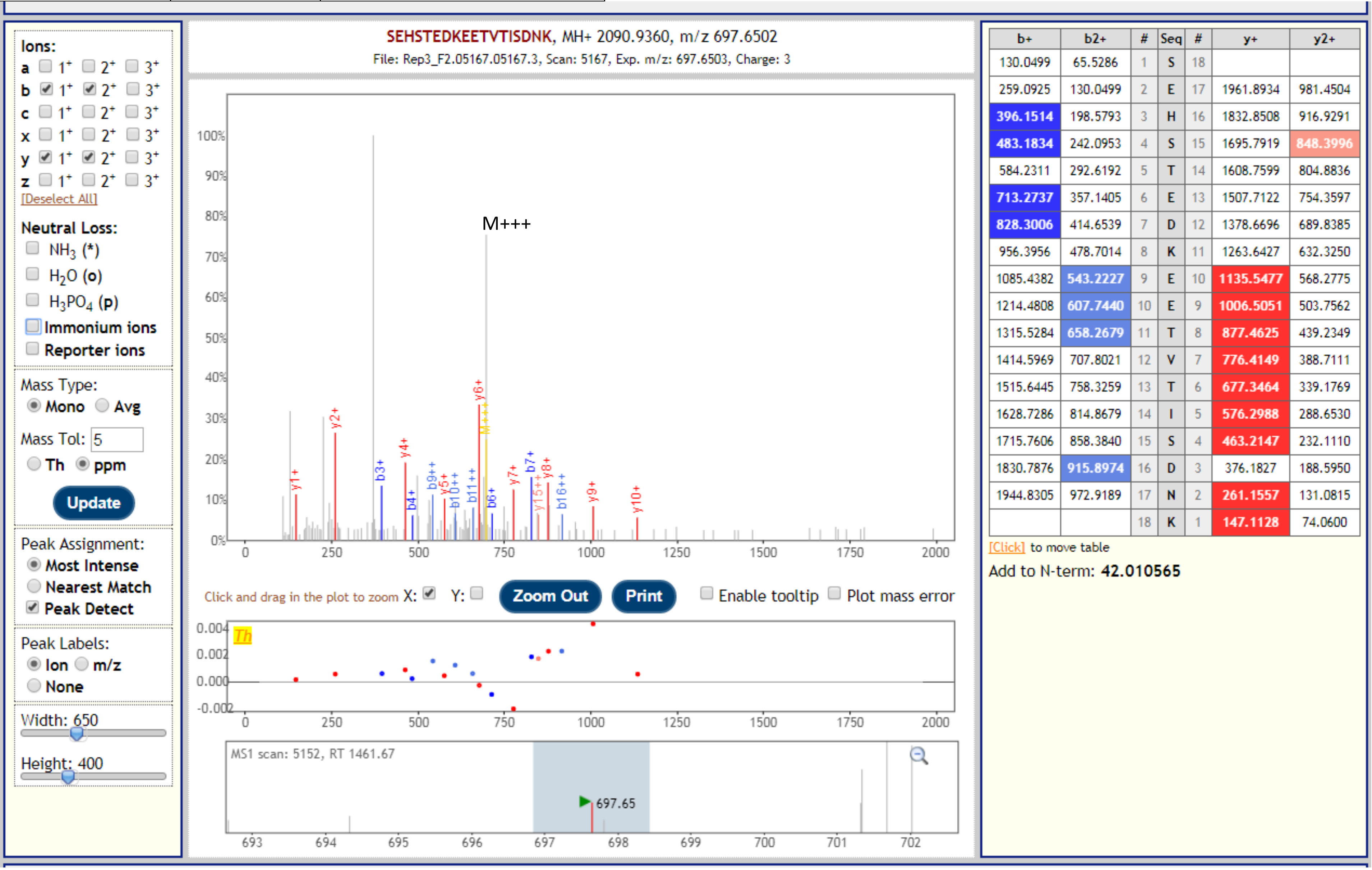


| b+ | b2+ | # | Seq | # | y+ | y2+ |
|---|---|---|---|---|---|---|
| 130.0499 | 65.5286 | 1 | S | 18 | | |
| 259.0925 | 130.0499 | 2 | E | 17 | 1961.8934 | 981.4504 |
| 396.1514 | 198.5793 | 3 | H | 16 | 1832.8508 | 916.9291 |
| 483.1834 | 242.0953 | 4 | S | 15 | 1695.7919 | 848.3996 |
| 584.2311 | 292.6192 | 5 | T | 14 | 1608.7599 | 804.8836 |
| 713.2737 | 357.1405 | 6 | E | 13 | 1507.7122 | 754.3597 |
| 828.3006 | 414.6539 | 7 | D | 12 | 1378.6696 | 689.8385 |
| 956.3956 | 478.7014 | 8 | K | 11 | 1263.6427 | 632.3250 |
| 1085.4382 | 543.2227 | 9 | E | 10 | 1135.5477 | 568.2775 |
| 1214.4808 | 607.7440 | 10 | E | 9 | 1006.5051 | 503.7562 |
| 1315.5284 | 658.2679 | 11 | T | 8 | 877.4625 | 439.2349 |
| 1414.5969 | 707.8021 | 12 | V | 7 | 776.4149 | 388.7111 |
| 1515.6445 | 758.3259 | 13 | T | 6 | 677.3464 | 339.1769 |
| 1628.7286 | 814.8679 | 14 | I | 5 | 576.2988 | 288.6530 |
| 1715.7606 | 858.3840 | 15 | S | 4 | 463.2147 | 232.1110 |
| 1830.7876 | 915.8974 | 16 | D | 3 | 376.1827 | 188.5950 |
| 1944.8305 | 972.9189 | 17 | N | 2 | 261.1557 | 131.0815 |
| | | 18 | K | 1 | 147.1128 | 74.0600 |

| Gene | Description | Peptide with cleaved & acetylated PEXEL |
|---|---|---|
| PBANKA_1465000 | fam-c protein | RSL.SEHSTEDKEETVTISDNKSSK |

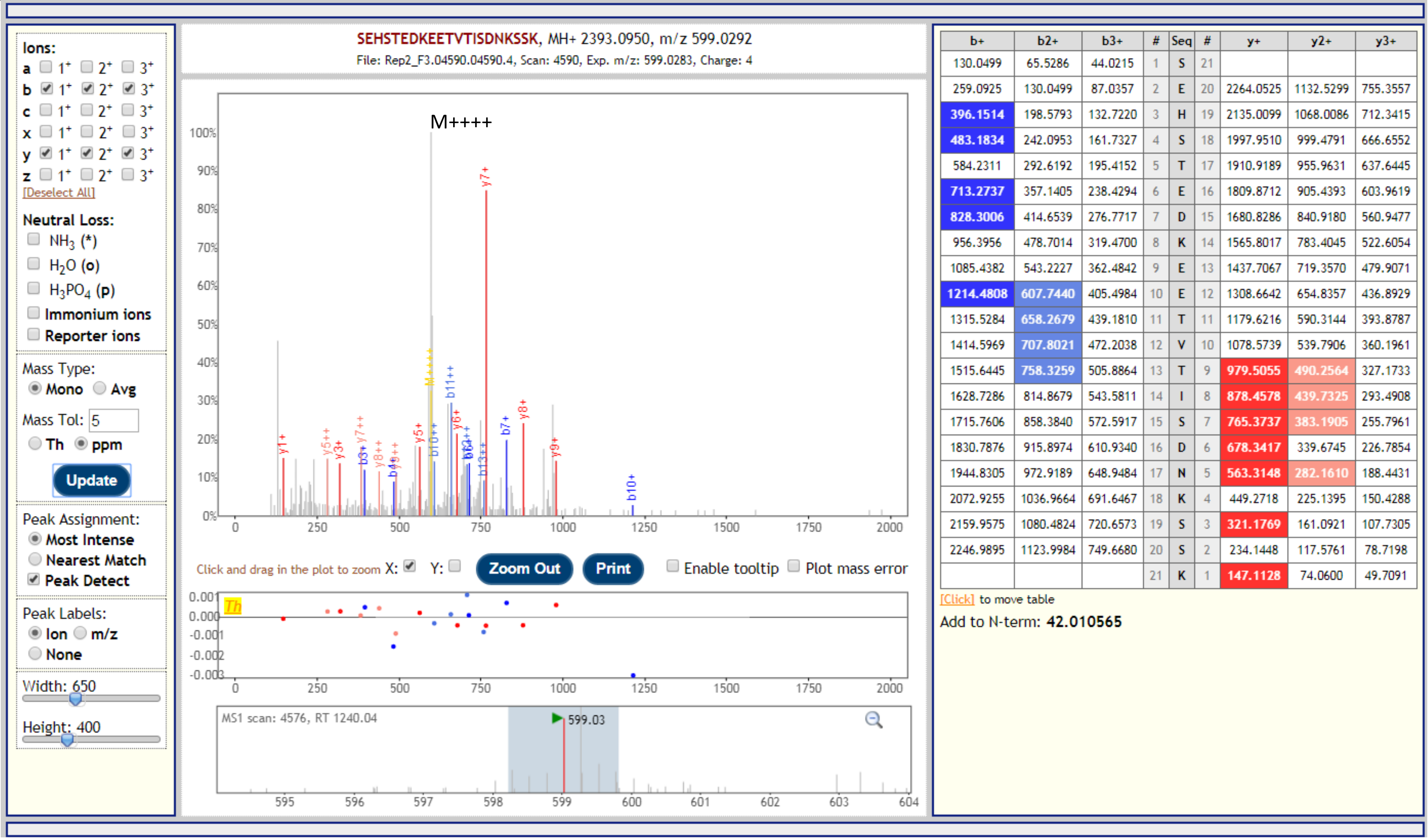


| b+ | b2+ | b3+ | # | Seq | # | y+ | y2+ | y3+ |
|---|---|---|---|---|---|---|---|---|
| 130.0499 | 65.5286 | 44.0215 | 1 | S | 21 | | | |
| 259.0925 | 130.0499 | 87.0357 | 2 | E | 20 | 2264.0525 | 1132.5299 | 755.3557 |
| 396.1514 | 198.5793 | 132.7220 | 3 | H | 19 | 2135.0099 | 1068.0086 | 712.3415 |
| 483.1834 | 242.0953 | 161.7327 | 4 | S | 18 | 1997.9510 | 999.4791 | 666.6552 |
| 584.2311 | 292.6192 | 195.4152 | 5 | T | 17 | 1910.9189 | 955.9631 | 637.6445 |
| 713.2737 | 357.1405 | 238.4294 | 6 | E | 16 | 1809.8712 | 905.4393 | 603.9619 |
| 828.3006 | 414.6539 | 276.7717 | 7 | D | 15 | 1680.8286 | 840.9180 | 560.9477 |
| 956.3956 | 478.7014 | 319.4700 | 8 | K | 14 | 1565.8017 | 783.4045 | 522.6054 |
| 1085.4382 | 543.2227 | 362.4842 | 9 | E | 13 | 1437.7067 | 719.3570 | 479.9071 |
| 1214.4808 | 607.7440 | 405.4984 | 10 | E | 12 | 1308.6642 | 654.8357 | 436.8929 |
| 1315.5284 | 658.2679 | 439.1810 | 11 | T | 11 | 1179.6216 | 590.3144 | 393.8787 |
| 1414.5969 | 707.8021 | 472.2038 | 12 | V | 10 | 1078.5739 | 539.7906 | 360.1961 |
| 1515.6445 | 758.3259 | 505.8864 | 13 | T | 9 | 979.5055 | 490.2564 | 327.1733 |
| 1628.7286 | 814.8679 | 543.5811 | 14 | I | 8 | 878.4578 | 439.7325 | 293.4908 |
| 1715.7606 | 858.3840 | 572.5917 | 15 | S | 7 | 765.3737 | 383.1905 | 255.7961 |
| 1830.7876 | 915.8974 | 610.9340 | 16 | D | 6 | 678.3417 | 339.6745 | 226.7854 |
| 1944.8305 | 972.9189 | 648.9484 | 17 | N | 5 | 563.3148 | 282.1610 | 188.4431 |
| 2072.9255 | 1036.9664 | 691.6467 | 18 | K | 4 | 449.2718 | 225.1395 | 150.4288 |
| 2159.9575 | 1080.4824 | 720.6573 | 19 | S | 3 | 321.1769 | 161.0921 | 107.7305 |
| 2246.9895 | 1123.9984 | 749.6680 | 20 | S | 2 | 234.1448 | 117.5761 | 78.7198 |
| | | | 21 | K | 1 | 147.1128 | 74.0600 | 49.7091 |

| Gene | Description | Peptide with cleaved & acetylated PEXEL |
|---|---|---|
| PBANKA_0700700 | Plasmodium exported protein, unknown function | RIL.SSMSNEKPYNILR |

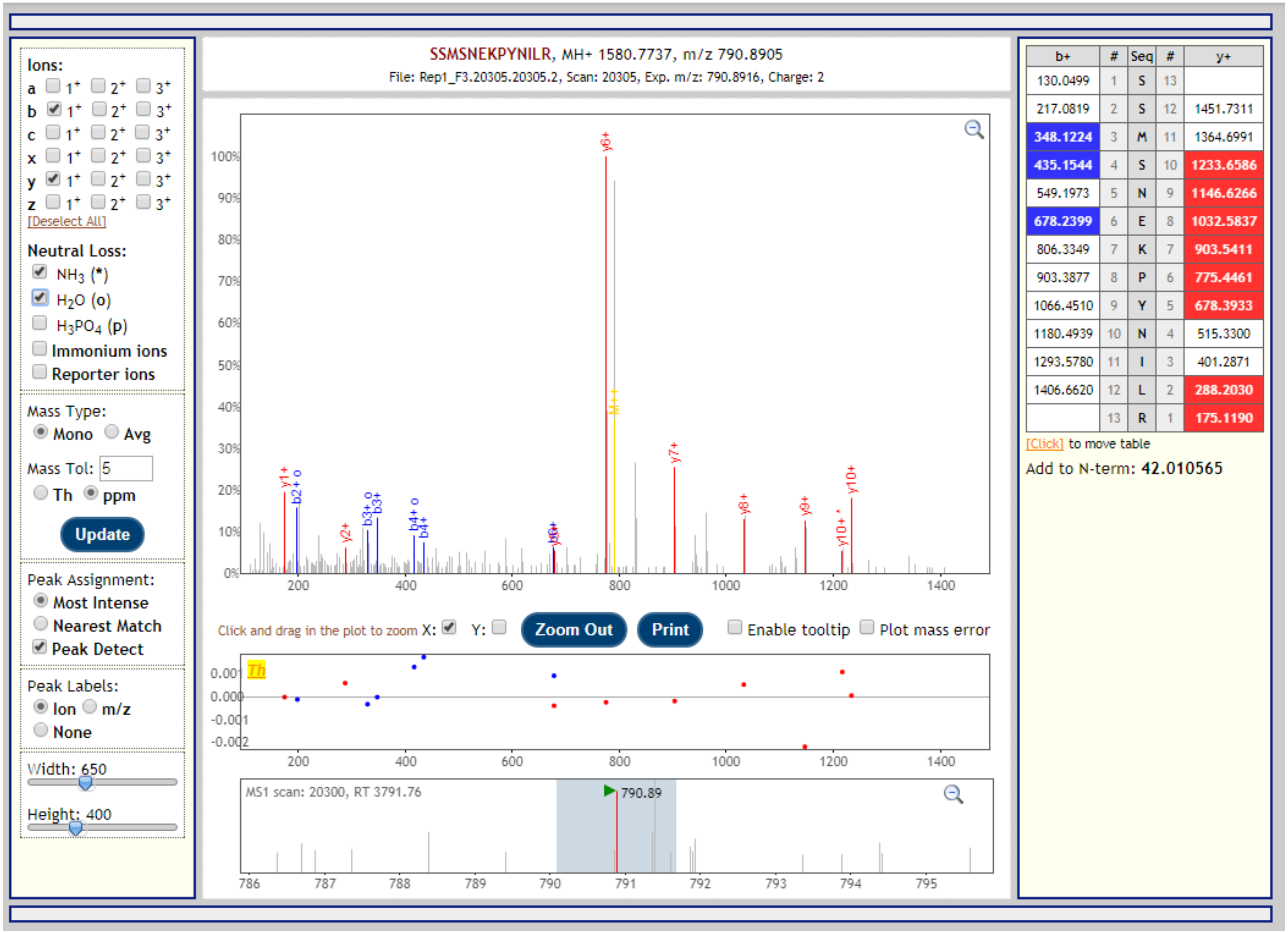

| Gene | Description | Peptide with cleaved & acetylated PEXEL |
|---|---|---|
| PBANKA_0700700 | Plasmodium exported protein, unknown function | RIL.SSMSNEKPYNILR |

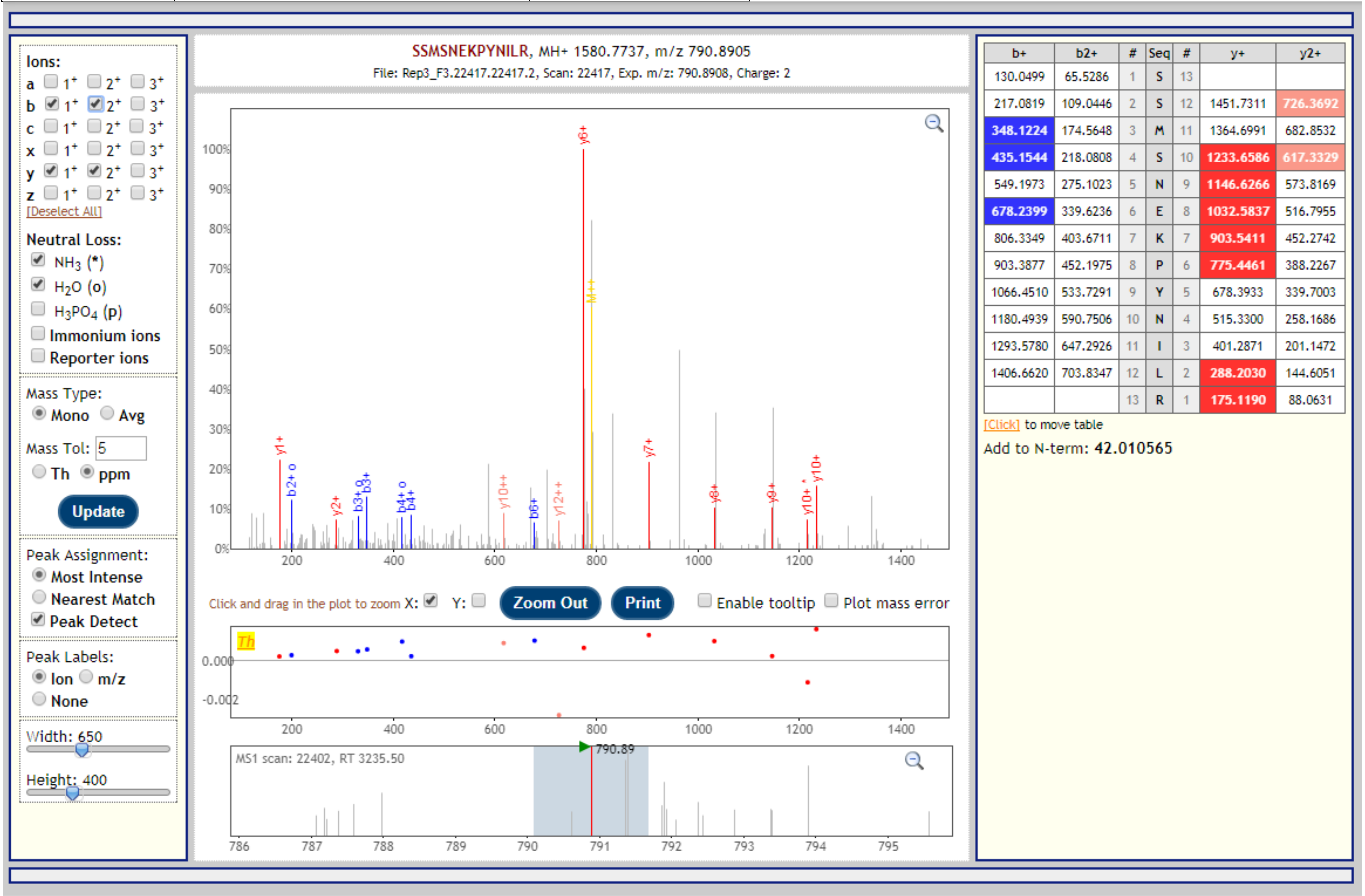


| b+ | b2+ | # | Seq | # | y+ | y2+ |
|---|---|---|---|---|---|---|
| 130.0499 | 65.5286 | 1 | S | 13 | | |
| 217.0819 | 109.0446 | 2 | S | 12 | 1451.7311 | 726.3692 |
| 348.1224 | 174.5648 | 3 | M | 11 | 1364.6991 | 682.8532 |
| 435.1544 | 218.0808 | 4 | S | 10 | 1233.6586 | 617.3329 |
| 549.1973 | 275.1023 | 5 | N | 9 | 1146.6266 | 573.8169 |
| 678.2399 | 339.6236 | 6 | E | 8 | 1032.5837 | 516.7955 |
| 806.3349 | 403.6711 | 7 | K | 7 | 903.5411 | 452.2742 |
| 903.3877 | 452.1975 | 8 | P | 6 | 775.4461 | 388.2267 |
| 1066.4510 | 533.7291 | 9 | Y | 5 | 678.3933 | 339.7003 |
| 1180.4939 | 590.7506 | 10 | N | 4 | 515.3300 | 258.1686 |
| 1293.5780 | 647.2926 | 11 | I | 3 | 401.2871 | 201.1472 |
| 1406.6620 | 703.8347 | 12 | L | 2 | 288.2030 | 144.6051 |
| | | 13 | R | 1 | 175.1190 | 88.0631 |

| Gene | Description | Peptide with cleaved & acetylated PEXEL |
|---|---|---|
| PBANKA_0100600 | schizont membrane associated cytoadherence protein | RFL.VEYYDANIDQYNGNQSK |

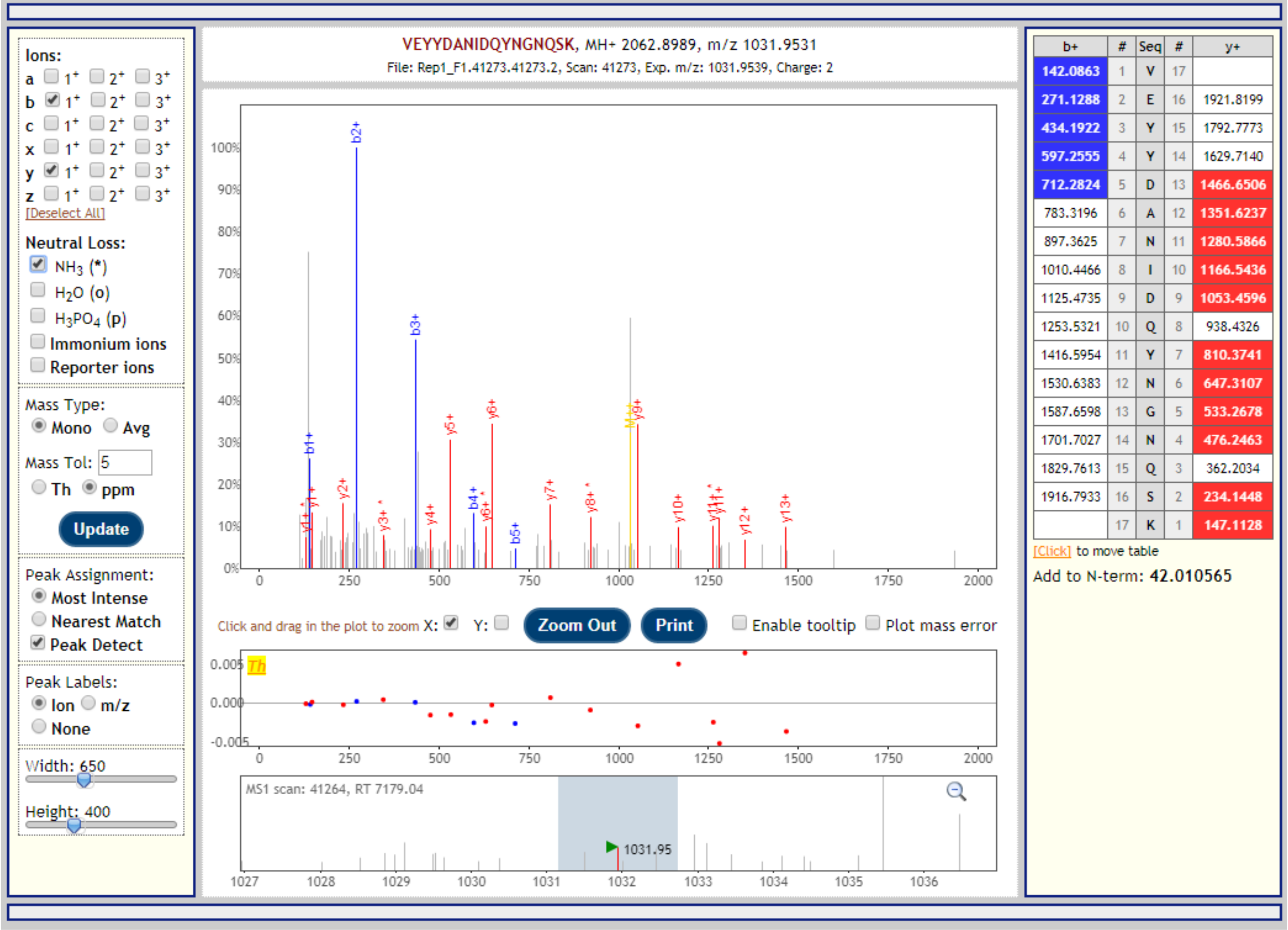

| Gene | Description | Peptide with cleaved & acetylated PEXEL |
|---|---|---|
| PBANKA_0100600 | schizont membrane associated cytoadherence protein | RFL.VEYYDANIDQYNGNQSK |

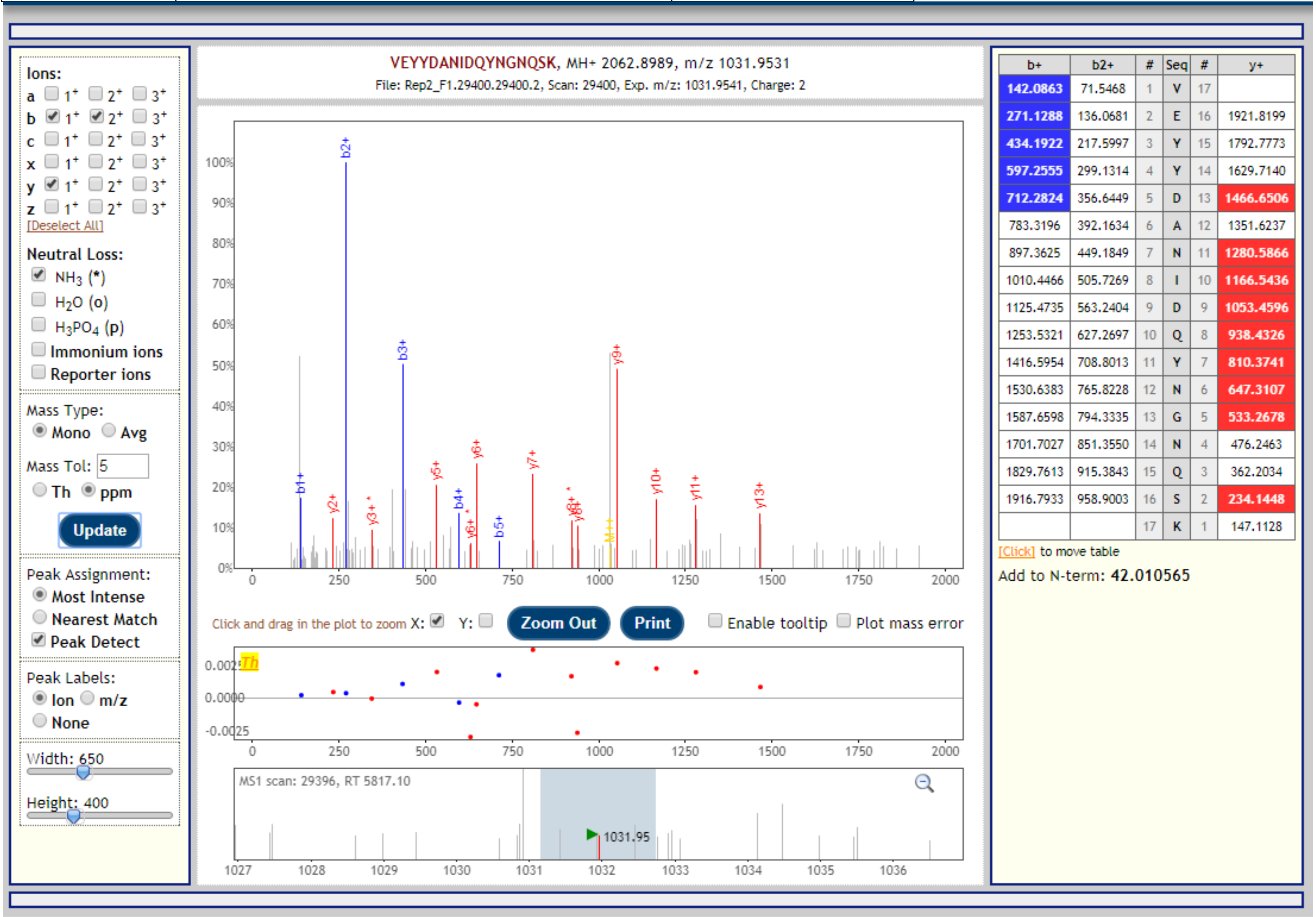


| b+ | b2+ | # | Seq | # | y+ |
|---|---|---|---|---|---|
| 142.0863 | 71.5468 | 1 | V | 17 | |
| 271.1288 | 136.0681 | 2 | E | 16 | 1921.8199 |
| 434.1922 | 217.5997 | 3 | Y | 15 | 1792.7773 |
| 597.2555 | 299.1314 | 4 | Y | 14 | 1629.7140 |
| 712.2824 | 356.6449 | 5 | D | 13 | 1466.6506 |
| 783.3196 | 392.1634 | 6 | A | 12 | 1351.6237 |
| 897.3625 | 449.1849 | 7 | N | 11 | 1280.5866 |
| 1010.4466 | 505.7269 | 8 | I | 10 | 1166.5436 |
| 1125.4735 | 563.2404 | 9 | D | 9 | 1053.4596 |
| 1253.5321 | 627.2697 | 10 | Q | 8 | 938.4326 |
| 1416.5954 | 708.8013 | 11 | Y | 7 | 810.3741 |
| 1530.6383 | 765.8228 | 12 | N | 6 | 647.3107 |
| 1587.6598 | 794.3335 | 13 | G | 5 | 533.2678 |
| 1701.7027 | 851.3550 | 14 | N | 4 | 476.2463 |
| 1829.7613 | 915.3843 | 15 | Q | 3 | 362.2034 |
| 1916.7933 | 958.9003 | 16 | S | 2 | 234.1448 |
| | | 17 | K | 1 | 147.1128 |

| Gene | Description | Peptide with cleaved & acetylated PEXEL |
|---|---|---|
| PBANKA_1145400 | Plasmodium exported protein (PHIST), unknown function | RNL.SETSVVNDNLSNNVNNLRDEPK |

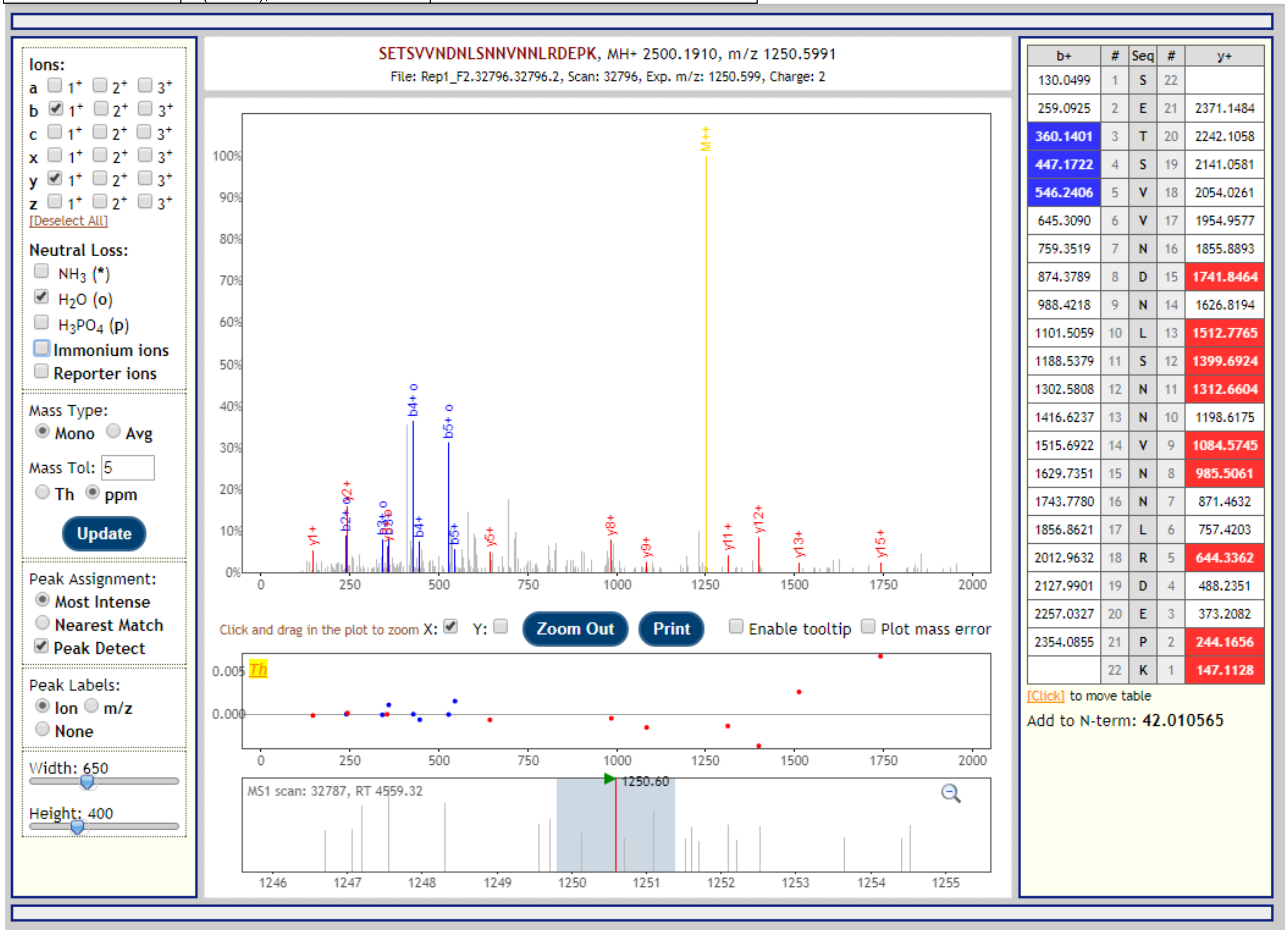

| Gene | Description | Peptide with cleaved & acetylated PEXEL |
|---|---|---|
| PBANKA_1106800 | subtilisin-like protease 3, putative | RIL.NQINNKELEINKNMK |

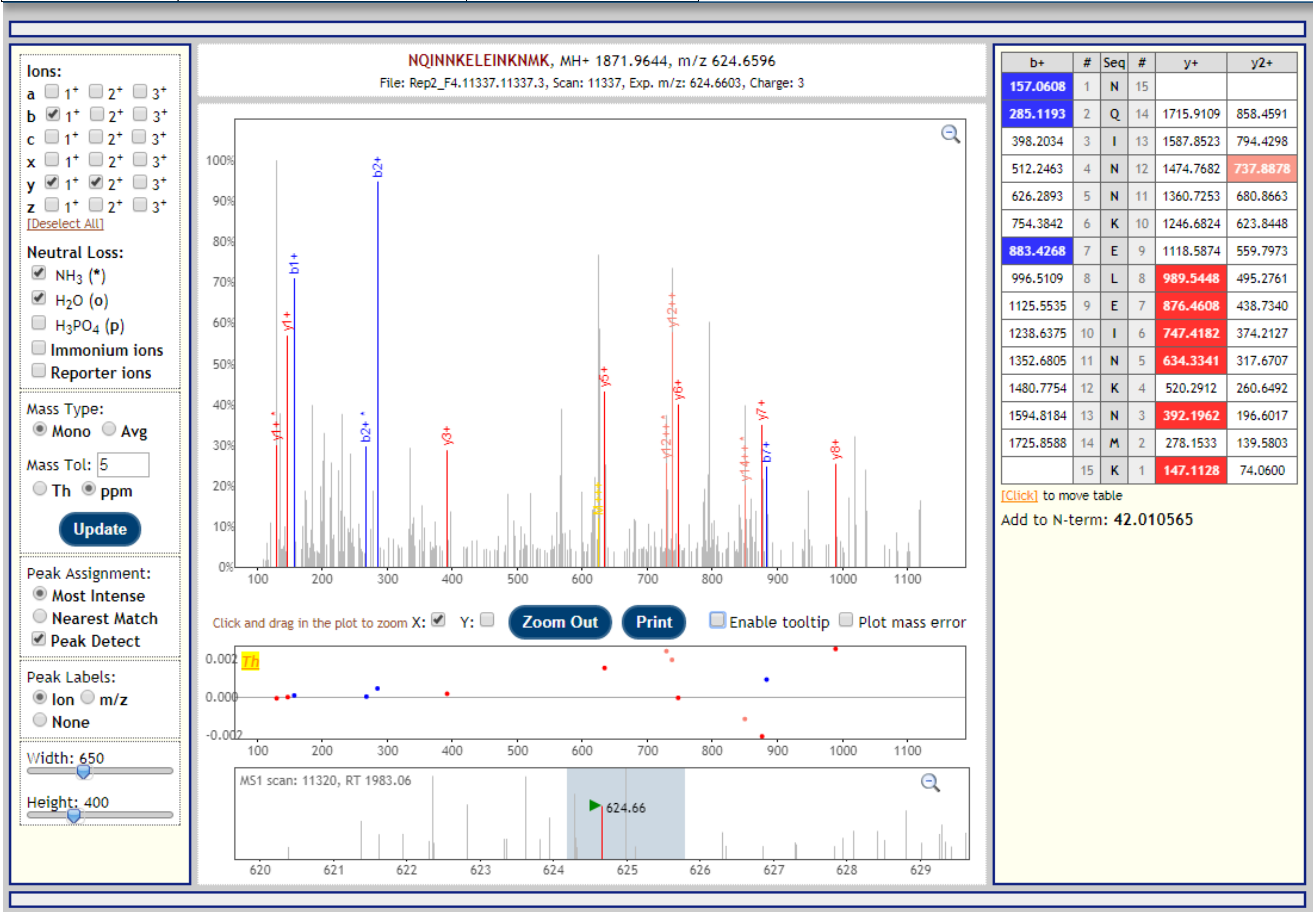

| Gene | Protein | Description | Peptide with cleaved & acetylated PEXEL |
|---|---|---|---|
| PBANKA_1106800 | SUB3 | subtilisin-like protease 3, putative | RIL.NQINNKELEINK |

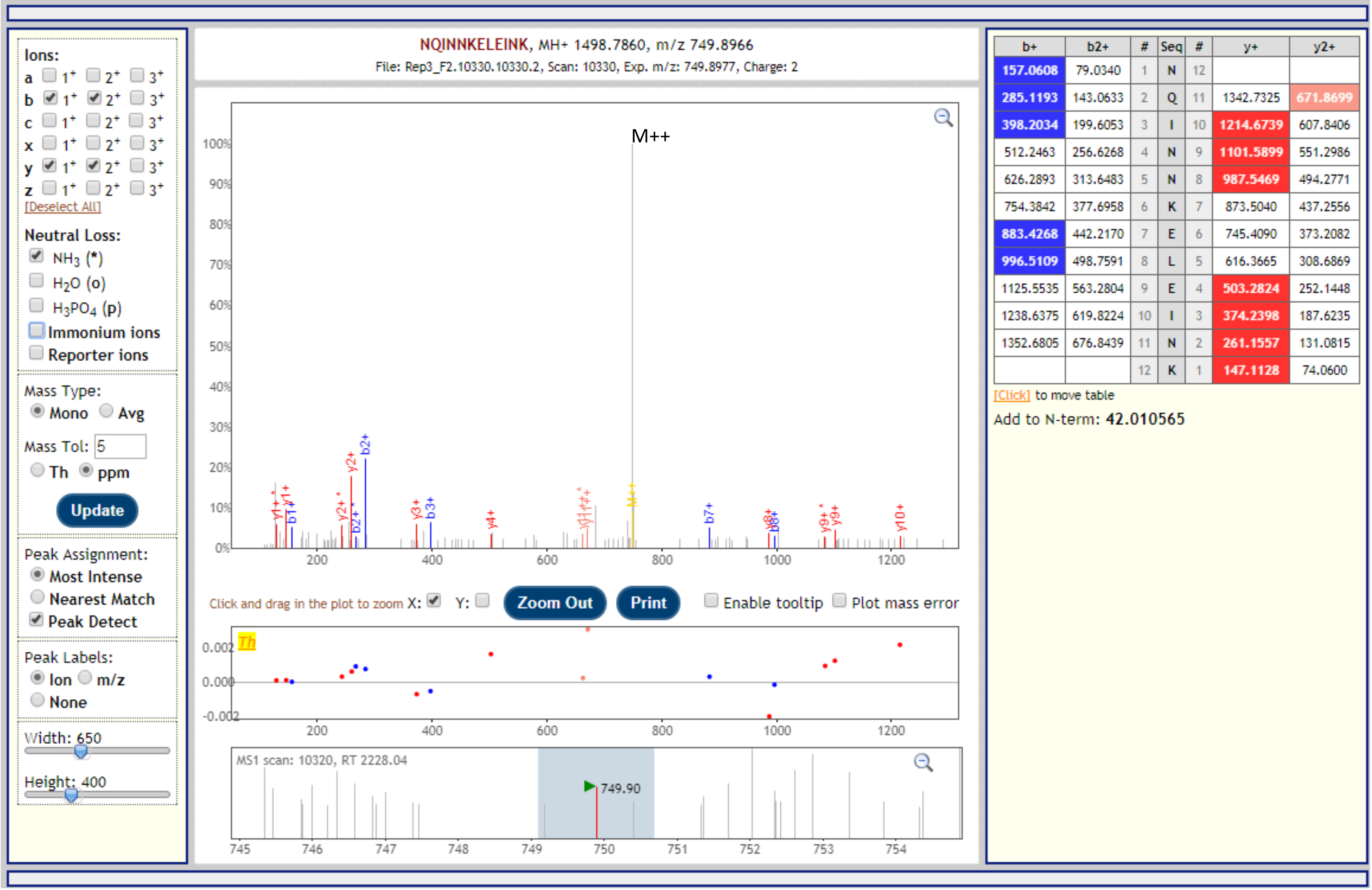


| b+ | b2+ | # | Seq | # | y+ | y2+ |
|---|---|---|---|---|---|---|
| 157.0608 | 79.0340 | 1 | N | 12 | | |
| 285.1193 | 143.0633 | 2 | Q | 11 | 1342.7325 | 671.8699 |
| 398.2034 | 199.6053 | 3 | I | 10 | 1214.6739 | 607.8406 |
| 512.2463 | 256.6268 | 4 | N | 9 | 1101.5899 | 551.2986 |
| 626.2893 | 313.6483 | 5 | N | 8 | 987.5469 | 494.2771 |
| 754.3842 | 377.6958 | 6 | K | 7 | 873.5040 | 437.2556 |
| 883.4268 | 442.2170 | 7 | E | 6 | 745.4090 | 373.2082 |
| 996.5109 | 498.7591 | 8 | L | 5 | 616.3665 | 308.6869 |
| 1125.5535 | 563.2804 | 9 | E | 4 | 503.2824 | 252.1448 |
| 1238.6375 | 619.8224 | 10 | I | 3 | 374.2398 | 187.6235 |
| 1352.6805 | 676.8439 | 11 | N | 2 | 261.1557 | 131.0815 |
| | | 12 | K | 1 | 147.1128 | 74.0600 |

| Gene | Description | Peptide with cleaved & acetylated PEXEL |
|---|---|---|
| PBANKA_1200041, PBANKA_1246961, PBANKA_1300041, PBANKA_0007801 | fam-b protein | RIL.AYADNEFDLNGFYQSTLNLASQLGDCVEGNKEIAHLR |

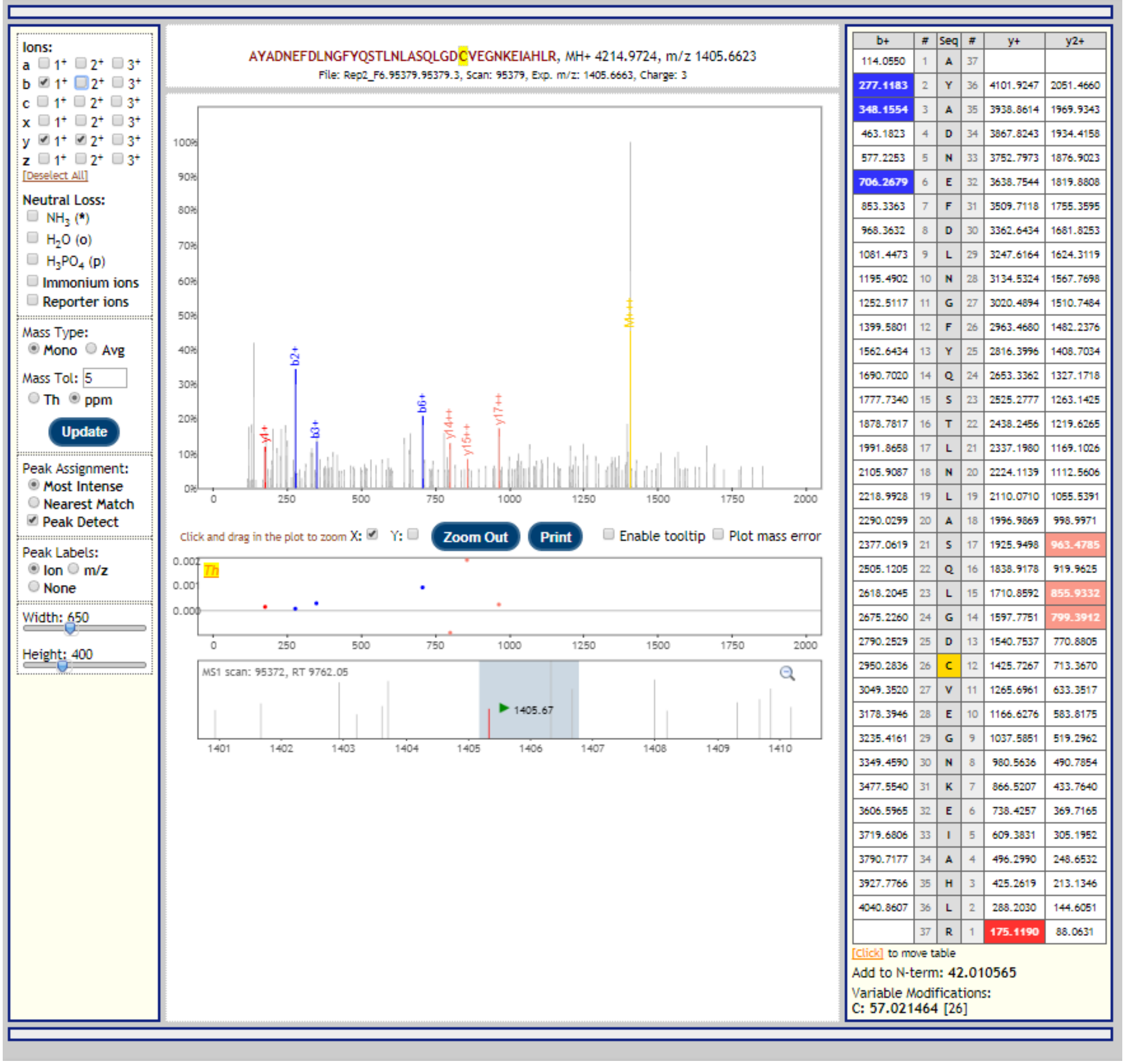


| b+ | # | Seq | # | y+ | y2+ |
|---|---|---|---|---|---|
| 114.0550 | 1 | A | 37 | | |
| 277.1183 | 2 | Y | 36 | 4101.9247 | 2051.4660 |
| 348.1554 | 3 | A | 35 | 3938.8614 | 1969.9343 |
| 463.1823 | 4 | D | 34 | 3867.8243 | 1934.4158 |
| 577.2253 | 5 | N | 33 | 3752.7973 | 1876.9023 |
| 706.2679 | 6 | E | 32 | 3638.7544 | 1819.8808 |
| 853.3363 | 7 | F | 31 | 3509.7118 | 1755.3595 |
| 968.3632 | 8 | D | 30 | 3362.6434 | 1681.8253 |
| 1081.4473 | 9 | L | 29 | 3247.6164 | 1624.3119 |
| 1195.4902 | 10 | N | 28 | 3134.5324 | 1567.7698 |
| 1252.5117 | 11 | G | 27 | 3020.4894 | 1510.7484 |
| 1399.5801 | 12 | F | 26 | 2963.4680 | 1482.2376 |
| 1562.6434 | 13 | Y | 25 | 2816.3996 | 1408.7034 |
| 1690.7020 | 14 | Q | 24 | 2653.3362 | 1327.1718 |
| 1777.7340 | 15 | S | 23 | 2525.2777 | 1263.1425 |
| 1878.7817 | 16 | T | 22 | 2438.2456 | 1219.6265 |
| 1991.8658 | 17 | L | 21 | 2337.1980 | 1169.1026 |
| 2105.9087 | 18 | N | 20 | 2224.1139 | 1112.5606 |
| 2218.9928 | 19 | L | 19 | 2110.0710 | 1055.5391 |
| 2290.0299 | 20 | A | 18 | 1996.9869 | 998.9971 |
| 2377.0619 | 21 | S | 17 | 1925.9498 | 963.4785 |
| 2505.1205 | 22 | Q | 16 | 1838.9178 | 919.9625 |
| 2618.2045 | 23 | L | 15 | 1710.8592 | 855.9332 |
| 2675.2260 | 24 | G | 14 | 1597.7751 | 799.3912 |
| 2790.2529 | 25 | D | 13 | 1540.7537 | 770.8805 |
| 2950.2836 | 26 | C | 12 | 1425.7267 | 713.3670 |
| 3049.3520 | 27 | V | 11 | 1265.6961 | 633.3517 |
| 3178.3946 | 28 | E | 10 | 1166.6276 | 583.8175 |
| 3235.4161 | 29 | G | 9 | 1037.5851 | 519.2962 |
| 3349.4590 | 30 | N | 8 | 980.5636 | 490.7854 |
| 3477.5540 | 31 | K | 7 | 866.5207 | 433.7640 |
| 3606.5965 | 32 | E | 6 | 738.4257 | 369.7165 |
| 3719.6806 | 33 | I | 5 | 609.3831 | 305.1952 |
| 3790.7177 | 34 | A | 4 | 496.2990 | 248.6532 |
| 3927.7766 | 35 | H | 3 | 425.2619 | 213.1346 |
| 4040.8607 | 36 | L | 2 | 288.2030 | 144.6051 |
| | 37 | R | 1 | 175.1190 | 88.0631 |

| Gene | Description | Peptide with cleaved & acetylated PEXEL |
|---|---|---|
| PBANKA_0500741, PBANKA_1000071, PBANKA_1146781 | fam-b protein | RIL.SYADNEFDLNGFYQSTLNLANQLGDCVEGKKEIEHLR |

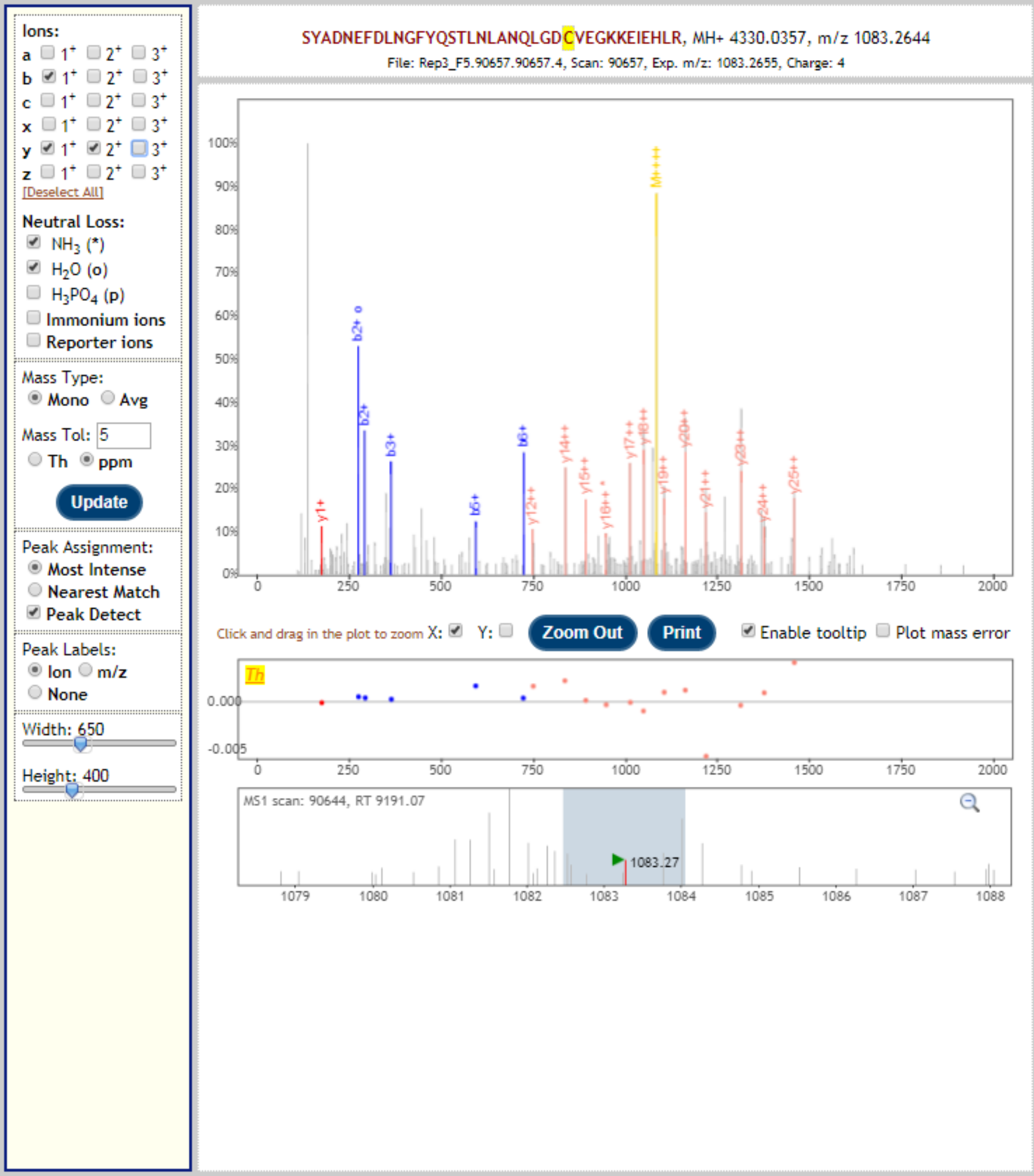


| b+ | # | Seq | # | y+ | y2+ |
|---|---|---|---|---|---|
| 130.0499 | 1 | S | 37 | | |
| 293.1132 | 2 | Y | 36 | 4200.9931 | 2101.0002 |
| 364.1503 | 3 | A | 35 | 4037.9298 | 2019.4685 |
| 479.1773 | 4 | D | 34 | 3966.8927 | 1983.9500 |
| 593.2202 | 5 | N | 33 | 3851.8657 | 1926.4365 |
| 722.2628 | 6 | E | 32 | 3737.8228 | 1869.4150 |
| 869.3312 | 7 | F | 31 | 3608.7802 | 1804.8937 |
| 984.3581 | 8 | D | 30 | 3461.7118 | 1731.3595 |
| 1097.4422 | 9 | L | 29 | 3346.6849 | 1673.8461 |
| 1211.4851 | 10 | N | 28 | 3233.6008 | 1617.3040 |
| 1268.5066 | 11 | G | 27 | 3119.5579 | 1560.2826 |
| 1415.5750 | 12 | F | 26 | 3062.5364 | 1531.7718 |
| 1578.6383 | 13 | Y | 25 | 2915.4680 | 1458.2376 |
| 1706.6969 | 14 | Q | 24 | 2752.4047 | 1376.7060 |
| 1793.7289 | 15 | S | 23 | 2624.3461 | 1312.6767 |
| 1894.7766 | 16 | T | 22 | 2537.3140 | 1269.1607 |
| 2007.8607 | 17 | L | 21 | 2436.2664 | 1218.6368 |
| 2121.9036 | 18 | N | 20 | 2323.1823 | 1162.0948 |
| 2234.9877 | 19 | L | 19 | 2209.1394 | 1105.0733 |
| 2306.0248 | 20 | A | 18 | 2096.0553 | 1048.5313 |
| 2420.0677 | 21 | N | 17 | 2025.0182 | 1013.0127 |
| 2548.1263 | 22 | Q | 16 | 1910.9753 | 955.9913 |
| 2661.2104 | 23 | L | 15 | 1782.9167 | 891.9620 |
| 2718.2318 | 24 | G | 14 | 1669.8326 | 835.4200 |
| 2833.2588 | 25 | D | 13 | 1612.8112 | 806.9092 |
| 2993.2894 | 26 | C | 12 | 1497.7842 | 749.3957 |
| 3092.3578 | 27 | V | 11 | 1337.7536 | 669.3804 |
| 3221.4004 | 28 | E | 10 | 1238.6852 | 619.8462 |
| 3278.4219 | 29 | G | 9 | 1109.6426 | 555.3249 |
| 3406.5168 | 30 | K | 8 | 1052.6211 | 526.8142 |
| 3534.6118 | 31 | K | 7 | 924.5261 | 462.7667 |
| 3663.6544 | 32 | E | 6 | 796.4312 | 398.7192 |
| 3776.7385 | 33 | I | 5 | 667.3886 | 334.1979 |
| 3905.7811 | 34 | E | 4 | 554.3045 | 277.6559 |
| 4042.8400 | 35 | H | 3 | 425.2619 | 213.1346 |
| 4155.9240 | 36 | L | 2 | 288.2030 | 144.6051 |
| | 37 | R | 1 | 175.1190 | 88.0631 |

[Click] to move table

Add to N-term: **42.010565**

Variable Modifications:
C: **57.021464** [26]

| Gene | Protein | Description | Peptide with cleaved & acetylated PEXEL |
|---|---|---|---|
| PBANKA_1342800 | NMNAT | nicotinamide/nicotinic acid mononucleotide adenylyltransferase, putative | KDL.ESENTTATYDLLNMLKK |

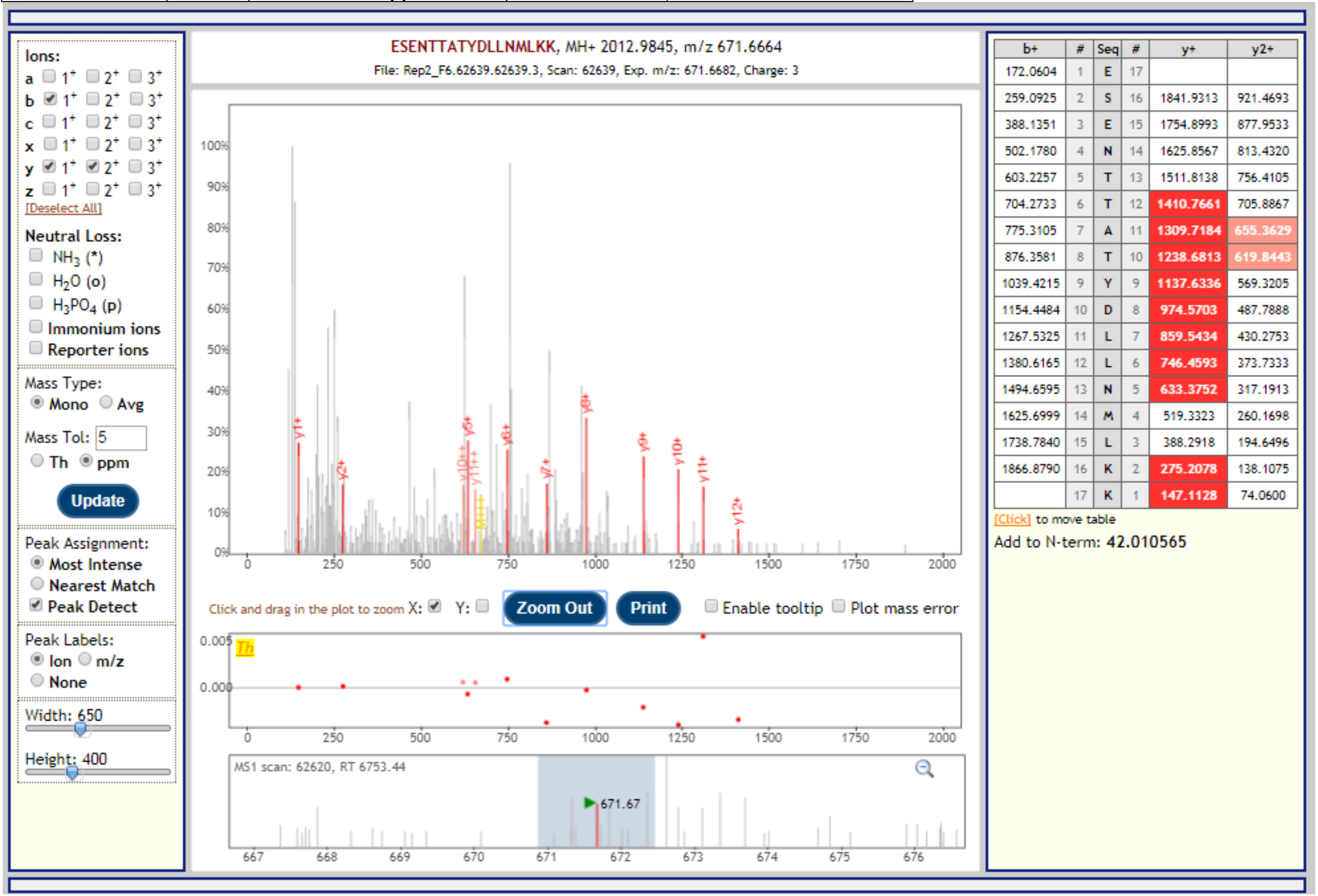


| b+ | # | Seq | # | y+ | y2+ |
|---|---|---|---|---|---|
| 172.0604 | 1 | E | 17 | | |
| 259.0925 | 2 | S | 16 | 1841.9313 | 921.4693 |
| 388.1351 | 3 | E | 15 | 1754.8993 | 877.9533 |
| 502.1780 | 4 | N | 14 | 1625.8567 | 813.4320 |
| 603.2257 | 5 | T | 13 | 1511.8138 | 756.4105 |
| 704.2733 | 6 | T | 12 | 1410.7661 | 705.8867 |
| 775.3105 | 7 | A | 11 | 1309.7184 | 655.3629 |
| 876.3581 | 8 | T | 10 | 1238.6813 | 619.8443 |
| 1039.4215 | 9 | Y | 9 | 1137.6336 | 569.3205 |
| 1154.4484 | 10 | D | 8 | 974.5703 | 487.7888 |
| 1267.5325 | 11 | L | 7 | 859.5434 | 430.2753 |
| 1380.6165 | 12 | L | 6 | 746.4593 | 373.7333 |
| 1494.6595 | 13 | N | 5 | 633.3752 | 317.1913 |
| 1625.6999 | 14 | M | 4 | 519.3323 | 260.1698 |
| 1738.7840 | 15 | L | 3 | 388.2918 | 194.6496 |
| 1866.8790 | 16 | K | 2 | 275.2078 | 138.1075 |
| | 17 | K | 1 | 147.1128 | 74.0600 |

| Gene | Description | Peptide with cleaved & acetylated PEXEL |
|---|---|---|
| PBANKA_1437200 | conserved Plasmodium protein, unknown function | RIL.VGSSKSHDIKEYDTNNFELIK |

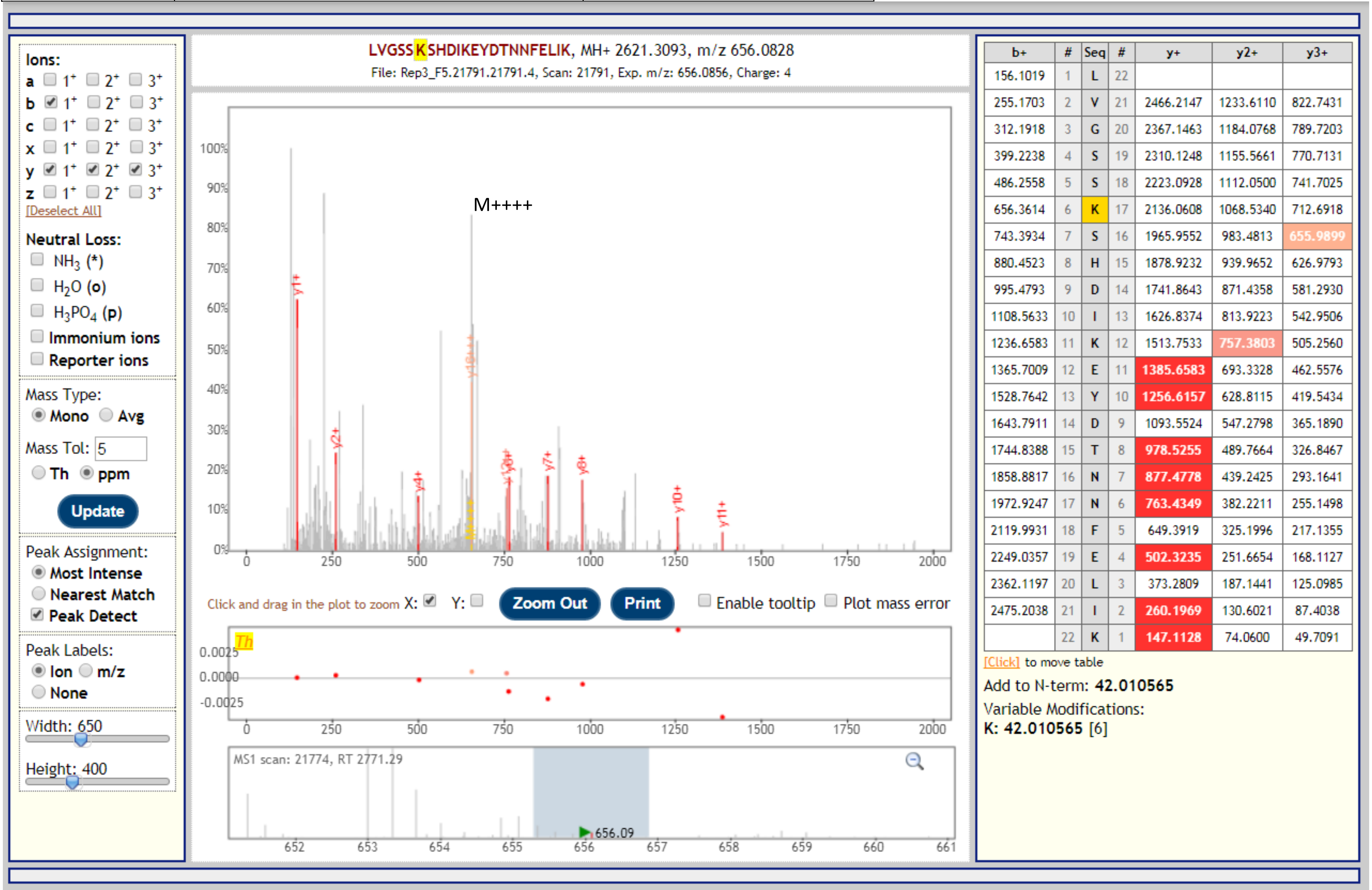


| b+ | # | Seq | # | y+ | y2+ | y3+ |
|---|---|---|---|---|---|---|
| 156.1019 | 1 | L | 22 | | | |
| 255.1703 | 2 | V | 21 | 2466.2147 | 1233.6110 | 822.7431 |
| 312.1918 | 3 | G | 20 | 2367.1463 | 1184.0768 | 789.7203 |
| 399.2238 | 4 | S | 19 | 2310.1248 | 1155.5661 | 770.7131 |
| 486.2558 | 5 | S | 18 | 2223.0928 | 1112.0500 | 741.7025 |
| 656.3614 | 6 | K | 17 | 2136.0608 | 1068.5340 | 712.6918 |
| 743.3934 | 7 | S | 16 | 1965.9552 | 983.4813 | 655.9899 |
| 880.4523 | 8 | H | 15 | 1878.9232 | 939.9652 | 626.9793 |
| 995.4793 | 9 | D | 14 | 1741.8643 | 871.4358 | 581.2930 |
| 1108.5633 | 10 | I | 13 | 1626.8374 | 813.9223 | 542.9506 |
| 1236.6583 | 11 | K | 12 | 1513.7533 | 757.3803 | 505.2560 |
| 1365.7009 | 12 | E | 11 | 1385.6583 | 693.3328 | 462.5576 |
| 1528.7642 | 13 | Y | 10 | 1256.6157 | 628.8115 | 419.5434 |
| 1643.7911 | 14 | D | 9 | 1093.5524 | 547.2798 | 365.1890 |
| 1744.8388 | 15 | T | 8 | 978.5255 | 489.7664 | 326.8467 |
| 1858.8817 | 16 | N | 7 | 877.4778 | 439.2425 | 293.1641 |
| 1972.9247 | 17 | N | 6 | 763.4349 | 382.2211 | 255.1498 |
| 2119.9931 | 18 | F | 5 | 649.3919 | 325.1996 | 217.1355 |
| 2249.0357 | 19 | E | 4 | 502.3235 | 251.6654 | 168.1127 |
| 2362.1197 | 20 | L | 3 | 373.2809 | 187.1441 | 125.0985 |
| 2475.2038 | 21 | I | 2 | 260.1969 | 130.6021 | 87.4038 |
| | 22 | K | 1 | 147.1128 | 74.0600 | 49.7091 |

| Gene | Description | Peptide with cleaved & acetylated PEXEL |
|---|---|---|
| PBANKA_0316200 | Plasmodium exported protein, unknown function | RIL.SELDEINGFTSEIR |

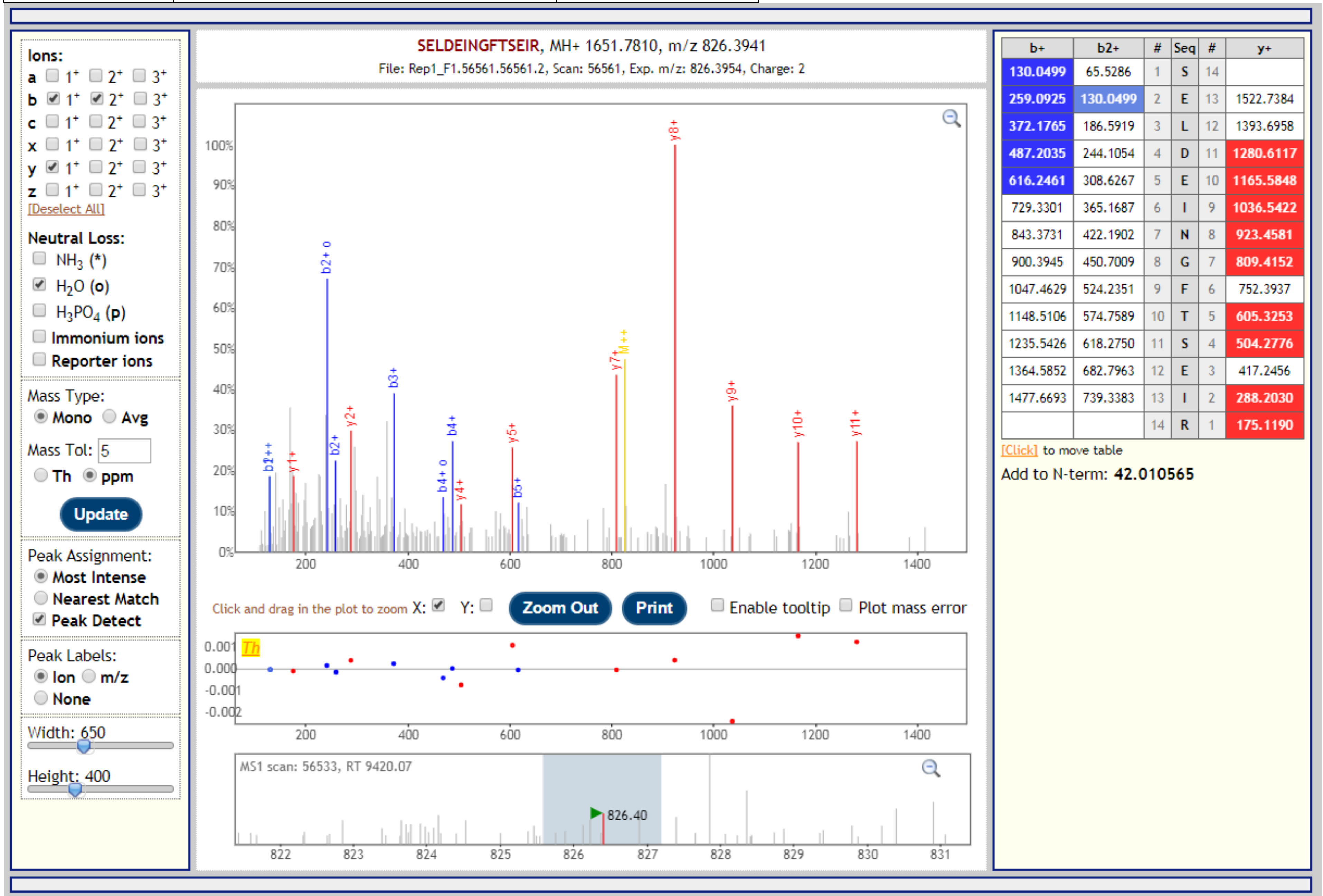

| Gene | Description | Peptide with cleaved & acetylated PEXEL |
|---|---|---|
| PBANKA_0100700 | Plasmodium exported protein, unknown function | RLL.AEPTNDLGVNK |

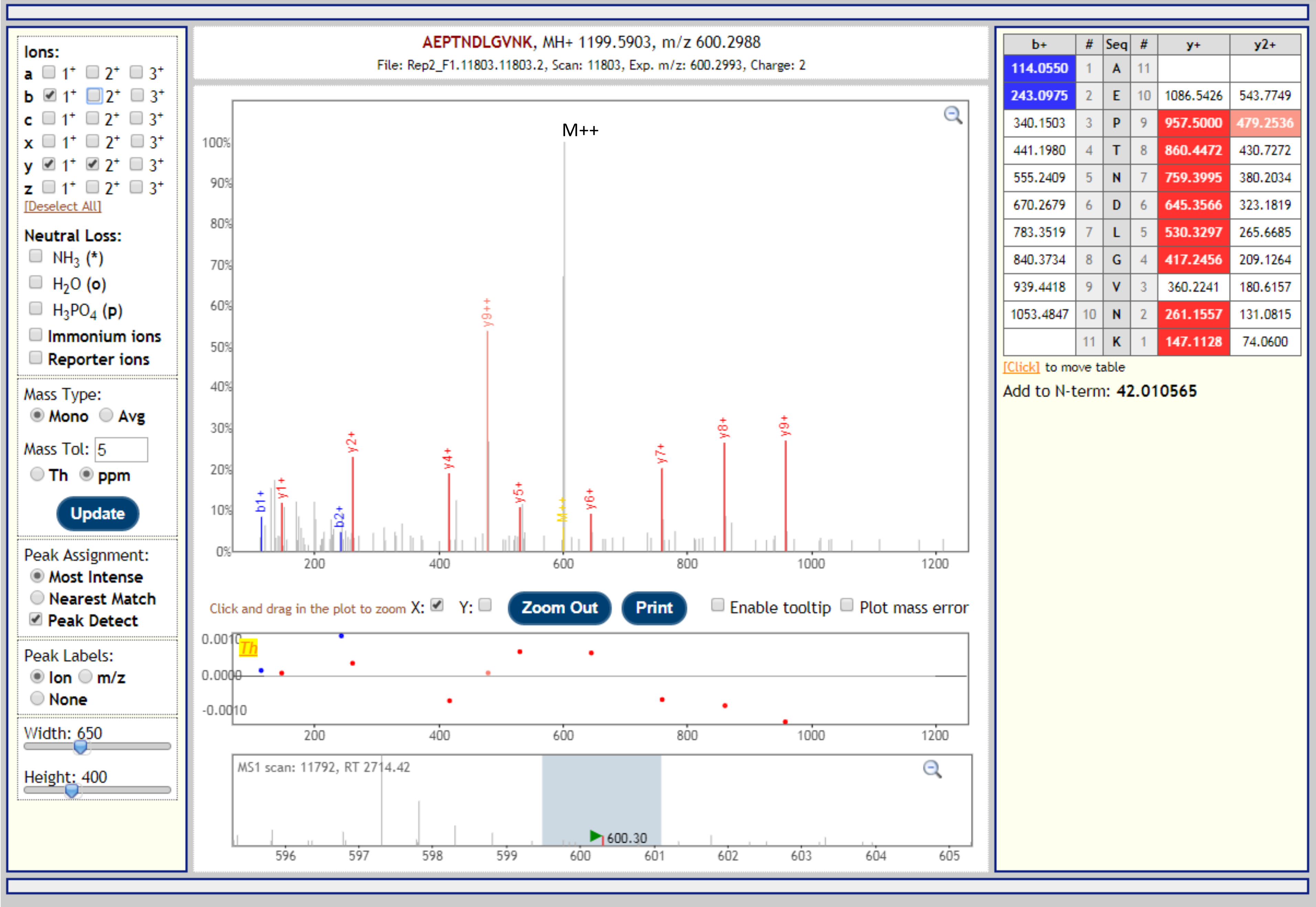

| Gene | Description | Peptide with cleaved & acetylated PEXEL |
|---|---|---|
| PBANKA_0715300 | conserved Plasmodium protein, unknown function | RSL.NENTTPNVMPIPDSKNEIINTESTISDIAEK |

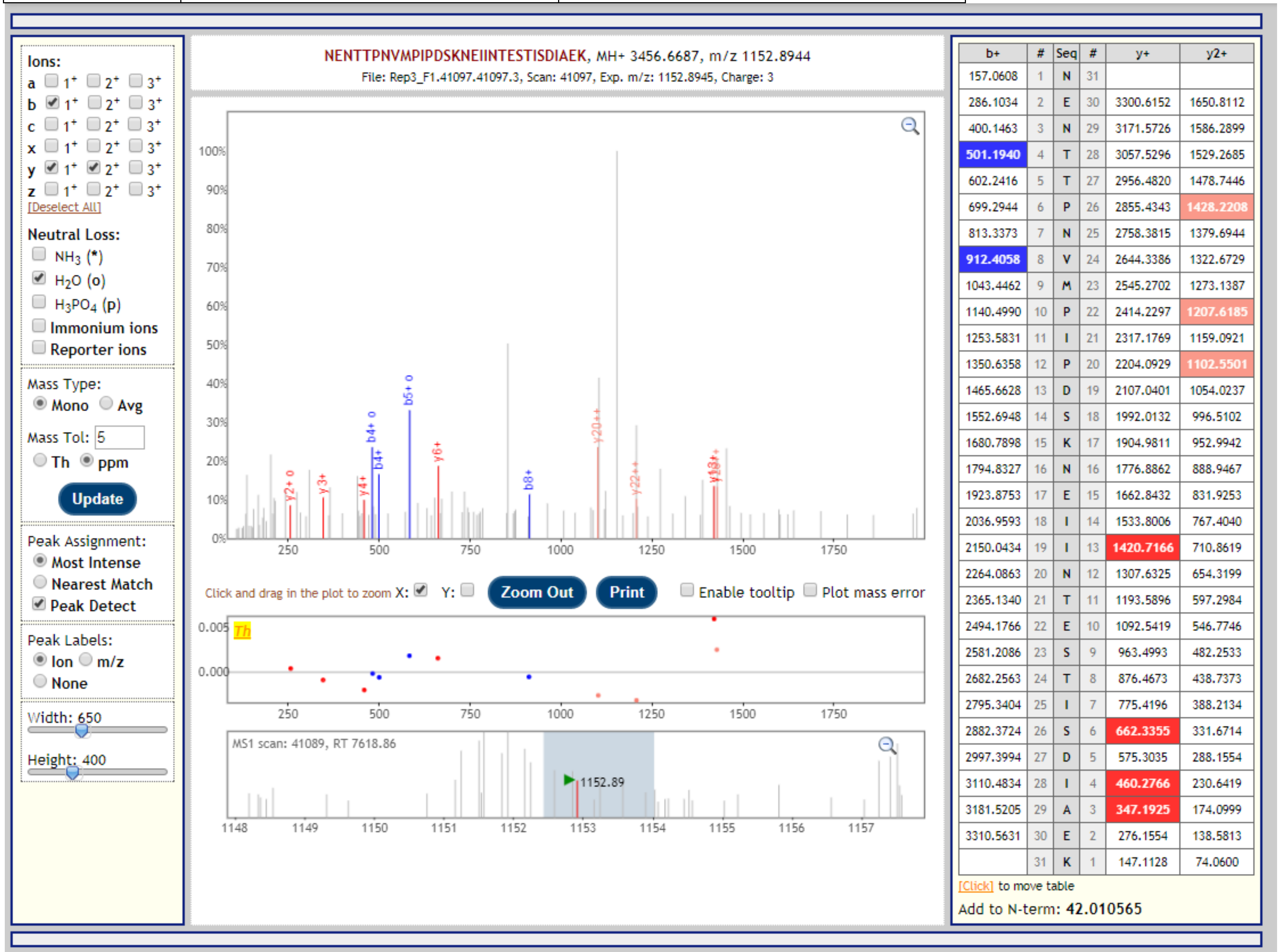


| b+ | # | Seq | # | y+ | y2+ |
|---|---|---|---|---|---|
| 157.0608 | 1 | N | 31 | | |
| 286.1034 | 2 | E | 30 | 3300.6152 | 1650.8112 |
| 400.1463 | 3 | N | 29 | 3171.5726 | 1586.2899 |
| 501.1940 | 4 | T | 28 | 3057.5296 | 1529.2685 |
| 602.2416 | 5 | T | 27 | 2956.4820 | 1478.7446 |
| 699.2944 | 6 | P | 26 | 2855.4343 | 1428.2208 |
| 813.3373 | 7 | N | 25 | 2758.3815 | 1379.6944 |
| 912.4058 | 8 | V | 24 | 2644.3386 | 1322.6729 |
| 1043.4462 | 9 | M | 23 | 2545.2702 | 1273.1387 |
| 1140.4990 | 10 | P | 22 | 2414.2297 | 1207.6185 |
| 1253.5831 | 11 | I | 21 | 2317.1769 | 1159.0921 |
| 1350.6358 | 12 | P | 20 | 2204.0929 | 1102.5501 |
| 1465.6628 | 13 | D | 19 | 2107.0401 | 1054.0237 |
| 1552.6948 | 14 | S | 18 | 1992.0132 | 996.5102 |
| 1680.7898 | 15 | K | 17 | 1904.9811 | 952.9942 |
| 1794.8327 | 16 | N | 16 | 1776.8862 | 888.9467 |
| 1923.8753 | 17 | E | 15 | 1662.8432 | 831.9253 |
| 2036.9593 | 18 | I | 14 | 1533.8006 | 767.4040 |
| 2150.0434 | 19 | I | 13 | 1420.7166 | 710.8619 |
| 2264.0863 | 20 | N | 12 | 1307.6325 | 654.3199 |
| 2365.1340 | 21 | T | 11 | 1193.5896 | 597.2984 |
| 2494.1766 | 22 | E | 10 | 1092.5419 | 546.7746 |
| 2581.2086 | 23 | S | 9 | 963.4993 | 482.2533 |
| 2682.2563 | 24 | T | 8 | 876.4673 | 438.7373 |
| 2795.3404 | 25 | I | 7 | 775.4196 | 388.2134 |
| 2882.3724 | 26 | S | 6 | 662.3355 | 331.6714 |
| 2997.3994 | 27 | D | 5 | 575.3035 | 288.1554 |
| 3110.4834 | 28 | I | 4 | 460.2766 | 230.6419 |
| 3181.5205 | 29 | A | 3 | 347.1925 | 174.0999 |
| 3310.5631 | 30 | E | 2 | 276.1554 | 138.5813 |
| | 31 | K | 1 | 147.1128 | 74.0600 |
